# Amyloidogenic proteolysis of APP regulates glutamatergic presynaptic function

**DOI:** 10.1101/2025.08.01.667924

**Authors:** Akshay Kapadia, Fabian Schuhmann, Ezgi Daskin, Jochen Walter, Isabell Lindahl, Neda Rahmani, Weria Pezeshkian, Anne-Sophie Hafner

## Abstract

Disease causing mutations of Alzheimer’s disease (AD) point to dysregulations of APP proteolysis. During asymptomatic and early stages of AD, brain recordings revealed hyperexcitation reverting into over-inhibition as dementia progresses. Here, we show that endogenous APP and its proteolytic product APP-CTFβ, the precursors of Aβ, accumulate preferentially at excitatory synapses. Using pharmacological treatments to modulate physiological concentrations of APP-CTFβ and Aβ, we identify APP-CTFβ as a key regulator of glutamatergic synaptic transmission. Accumulation of APP-CTFβ increases the release probability of synaptic vesicles. Strikingly, monomeric Aβ counteracts this APP-CTFβ-driven hyperexcitability. This suggests that therapeutic strategies clearing monomeric Aβ could be detrimental during the early hyperexcitability phase of AD.

## Introduction

Decades of research have centered on amyloid-β (Aβ) owing to its aggregation propensity and extracellular deposition as plaques, a key hallmark of Alzheimer’s disease (AD) pathology (*1*, *2*). Amyloid-beta precursor protein (APP), a transmembrane protein, expressed highly in neurons and regulating neuronal function (*3*), is the source of Aβ peptides. However, APP undergoes complex proteolysis, generating multiple fragments beyond Aβ, including extracellular soluble APP species (sAPP), membrane-bound APP C-terminal fragments (APP-CTFs) and cytosolic APP intracellular domain (AICD) (*4*, *5*), whose contributions to synaptic physiology and neurodegeneration remain relatively underexplored (*6–8*). Nevertheless, emerging evidence suggests that APP-CTFs, in particular APP-CTFβ (intermediate of the amyloidogenic pathway), independent of Aβ, could play an active role in triggering neuronal dyshomeostasis (*9–15*). Thus, understanding the precise roles of these fragments in physiological and pathological contexts is critical.

AD transgenic animal models, which encompass multiple familial AD mutations, are mainly developed on recapitulating Aβ plaque pathology, increasing the complexity of APP proteolysis (*16*). Furthermore, non-physiological levels (APP overexpression models), aberrant localization, interactions and proteolysis; potentially lead to artifactual phenotypes (*16–19*). Similarly, the use of synthetic APP-derived peptides often fails to replicate biological conformations or endogenous concentrations as well as cellular localization, complicating the interpretation of functional studies (*20*, *21*). These methodological and technical limitations have contributed to the persistent knowledge gap regarding the physiological and pathological roles of APP proteolytic products beyond Aβ, despite their important contributions to neuron pathophysiology.

Therapeutic strategies singularly aimed at reducing Aβ levels or its aggregation have faced significant challenges in clinical trials, making AD drug development a pharmaceutical graveyard. Aβ-directed therapies aiming at lowering Aβ production (β- or γ-secretase inhibitors), or clearing its excess (antibody-based immunotherapy), or preventing its aggregation (small molecule/peptide-based β-sheet breakers) will inadvertently interfere with its physiological function (*22–26*). Moreover, APP and its cleavage products have been shown to play roles in synaptic function (*6–8*), and targeting Aβ possibly disrupts APP-dependent processes that are essential for synaptic homeostasis. Addressing these concerns requires a more comprehensive understanding of APP processing and its precise impact on synaptic physiology, in an attempt to better delineate the multifaceted AD pathology.

In this study, we investigate the differential localization, site of action and function of endogenous APP and its proteolytic fragments in synapse subtypes. Using a combination of fluorescence-activated synapse sorting, biochemical fractionations, high-resolution imaging, functional assays *in vivo* and *in vitro* models as well as *in silico* computational modelling of the human C99, we examine how they influence synaptic activity and neuronal network dynamics, with a particular focus on APP-CTFβ. Our findings reveal distinct presynaptic roles for APP-CTFs/β, challenging the conventional Aβ-centric paradigm and providing new insights into the physiological functions of APP proteolysis.

## Results

### APP and enzymes of its amyloidogenic proteolysis predominantly localize in excitatory presynapses

Using biochemistry and immunofluorescence assays, APP and its processing enzymes have been shown to be present and locally synthesized at synapses (*27–31*). Nonetheless, efforts to localize those proteins in different synaptic compartments of animal brains have been challenged by the small size (a few micrometers) and density/proximity of these compartments in tissue (fig. S1). To circumvent these challenges, we optimized the method of fluorescence-activated synapse sorting (FASS) (*32*) to obtain two pools of synapses, respectively enriched in excitatory [GFP-positive, containing vesicular glutamate transporter 1 (vGLUT1)] and inhibitory synapses [GFP-negative, containing vesicular GABA transporter (vGAT)] from 1-year old mice (Fig. 1B-E, fig. S2A-C). The “presorted” and “sorted” synaptosomes constitute resealed presynaptic compartments, sometimes associated with “open” postsynaptic membranes (*32*). We validated the enrichment of both pools using immunocytochemistry on synaptosomes plated at low density on coverslips (Fig. 1C, fig. S2O) and ELISA-assays (Fig. 1D-E, fig. S2D-G).

**Figure 1.**
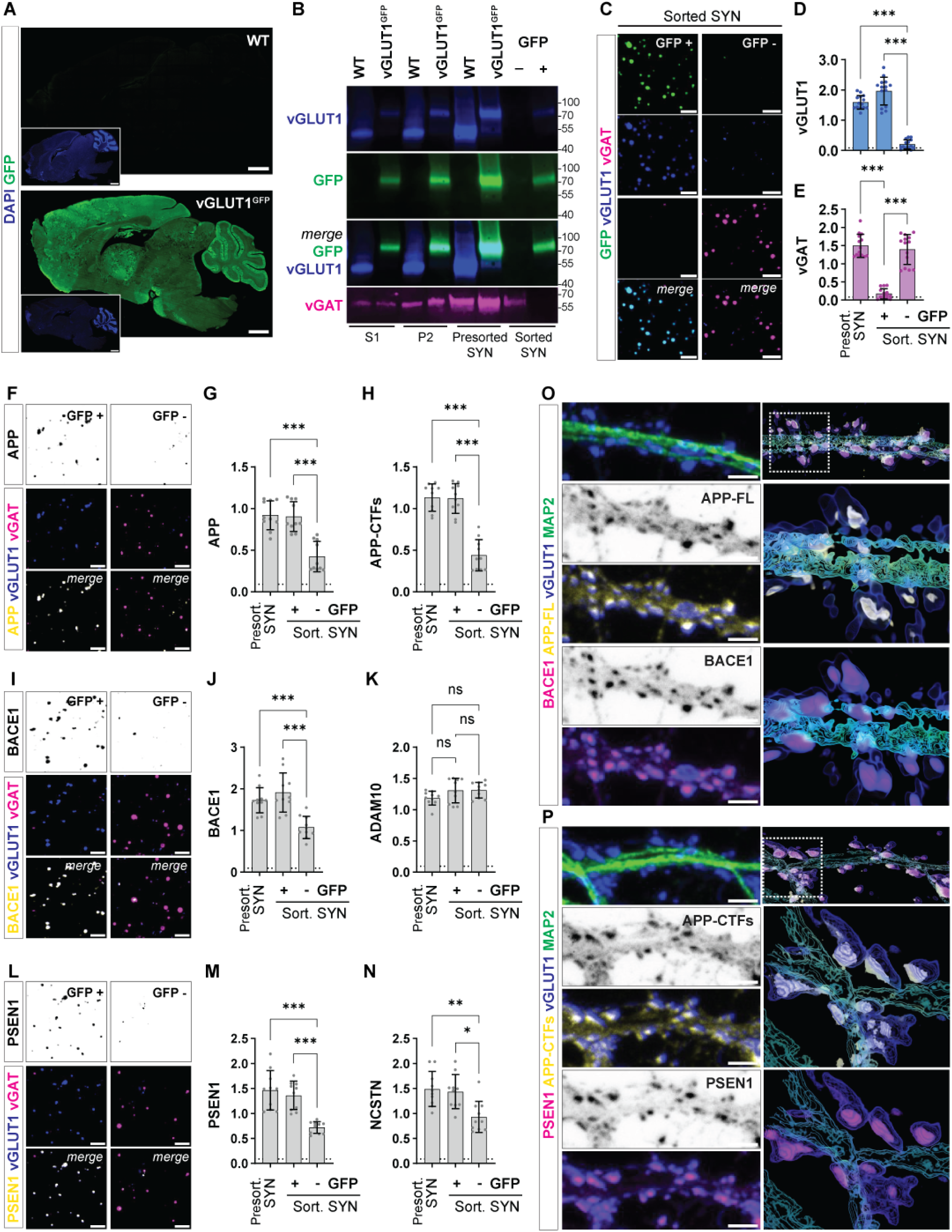
Amyloidogenic processing of APP predominantly occurs at the excitatory synapse. **(A)** Immunohistochemical analyses of wild type and vGLUT1^GFP^ transgenic mice sections (male, 6 weeks) showing GFP positive vGLUT1 (vesicular glutamatergic transporter 1, *green channel*) signals. Sections were co-stained for DAPI to visualize the nucleus (*blue channel*). Scale bar = 100 μm, N = 2. (**B**) Western immunoblots depicting the differential stages of synaptosome enrichment from WT and vGLUT1^GFP^ mice forebrain (male, 1 year); and sorting (*Fig. S2A, scheme*) of GFP+ (GFP positive) and GFP-(GFP negative) synapses. Blots were probed for α-vGLUT1 (*blue*), α-GFP (*green*) and α-vGAT (vesicular GABAergic transporter, *magenta*). Protein load-S1, P2, presort. SYN = 20 μg, Sort. SYN = 1 μg. N = 5. (**C**) Immunocytochemical (ICC) staining of sorted synapses co-stained for vGLUT1 (*blue*) and vGAT (*magenta*). Scale bar = 1 μm, n = 10, N = 5. (**D-E**) Indirect ELISA analyses depicting enrichment of vGLUT1 in GFP+ synapses (D) and vGAT in GFP-synapses (E), respectively. n = 15, N = 5. (**F, I, L**) ICC of sorted synapses stained for APP-FL (beta-amyloid precursor full-length protein; E, *gray, yellow-merge*), BACE1 (beta-amyloid precursor protein cleaving enzyme 1; H, *gray, yellow-merge*), and PSEN1 (presenilin-1, catalytic component of γ-secretase complex, K, *gray, yellow-merge*) along with vGLUT1 (*blue*) and vGAT (*magenta*). Scale bar = 1 μm, n = 10, N = 5. (**G, H, J, K, M, N**) Indirect ELISA analyses of presorted synapses (*P3 fraction*), and sorted GFP+ and GFP- synapses quantifying relative levels of APP-FL (F), APP C-terminal fragments (APP-CTFs, G), ADAM10 (metalloproteinase-10, α-secretase; J), BACE1 (I), PSEN1 (L) and NCSTN (Nicastrin, component of γ-secretase complex, M) in presorted and sorted - GFP+ and GFP-synapse fractions. n = 10, N = 5. (**O-P**) Immunocytochemistry and Imaris 3D reconstruction of rat primary cortical neurons (DIV21) depicting localization of APP-FL (N, *grayscale, yellow-merge*); BACE1 (N, *magenta*) and APP-CTFs (O, *grayscale, yellow-merge*) and PSEN1 (O, *magenta*) in excitatory synapses (vGLUT1+, *blue*). Neurons were co-stained with MAP2 to visualize the dendrites (*green*). Zoomed in sections have been indicated with a white box. Scale bar = 5 μm, N = 3. ns (*p* > 0.05), * (*p* ≤ 0.05), ** (*p* ≤ 0.01) , *** (*p* ≤ 0.01); One-way ANOVA.

Using these approaches, we then investigated the relative enrichment of endogenous levels of membrane bound APP, APP-CTFs - precursors of Aβ, as well as enzymes involved in APP proteolysis in presorted and sorted synapses (Fig. 1F-N, fig. S2H-O). We found that APP, its proteolytic products APP-CTFs, and enzymes of the amyloidogenic pathway β-secretase (BACE1) and 𝛾𝛾-secretase (PSEN1 and NCSTN) are present in excitatory synapses. While some signals were detectable for vGAT-enriched (GFP-negative) synapses immunocytochemistry revealed that most vGAT-positive synapses did not contain those proteins as opposed to vGLUT1-positive synapses (Fig. 1F, I, L; fig. S2O). Consequently, the metalloproteinase ADAM10, which cleaves APP *via* the non-amyloidogenic pathway is equally abundant in all synapse types (Fig. 1K, fig. S2J). We further confirmed the presence of APP, APP-CTFs, β-secretase and 𝛾𝛾-secretase in synapses enriched from dissociated rat neurons (fig. S1E-I) by performing immunocytochemistry (Fig. 1O- P, fig. S1C-D). Our results confirmed their accumulation in excitatory synapses detected using an antibody against vGLUT1.

### Accumulation of APP-CTFβ drives network hyperexcitability

A remarkable number of studies have linked APP and its proteolytic products, to a wide range of fundamental processes to maintain neuronal homeostasis (*6–8*). Since we found the amyloidogenic proteolysis of APP to be occurring at excitatory synapses, we investigated the effects of manipulating the relative abundance of those products using pharmacology, to systematically disentangle the role of each component. We note that our manipulations rely on endogenous levels of rodent APP, wherein the levels of APP and its proteolytic products remain within a plausible physiological range. Our treatments consisted of a 4 h incubation with one of the following drugs applied directly in the cell culture media: (i) LY2886721 to inhibit β-secretase (BSI), (ii) DAPT to inhibit 𝛾𝛾-secretase (GSI), and (iii) Aftin-4 to promote the entire pathway in order to increase the production of Aβ (APA) (Fig. 2A-E, fig. S3A-H). BSI led to a predicted decrease in soluble extracellular products, Aβ and sAPPβ; and a slight accumulation of intracellular (full length) APP-FL. Changes in APP might result from its endogenous high expression in neurons and/or compensatory mechanisms. In our experiments, β-secretase inhibition decreased synaptic but not global APP-CTFβ levels (Fig. 2C, fig. S3G, H, J, K). GSI, however, caused a sharp reduction in Aβ levels (∼50% decrease in detectable Aβ in media) and an accumulation of synaptic APP-CTFβ (∼2.5-fold increase, fig. S3J-K) in excitatory boutons (Fig. 2F, H). Finally, APA led to an increase in sAPPβ and Aβ (∼2-fold increase). Importantly, each treatment causes a specific molecular signature (Fig. 2B-E, fig. S3A-K), and differently impacts synapses (Fig. 2F-G, fig. S3I) along with spontaneous network activity measured by imaging neuronal calcium dynamics [GCaMP6 response] (Fig. 2I-J, fig. S4A-C, video S1).

**Figure 2.**
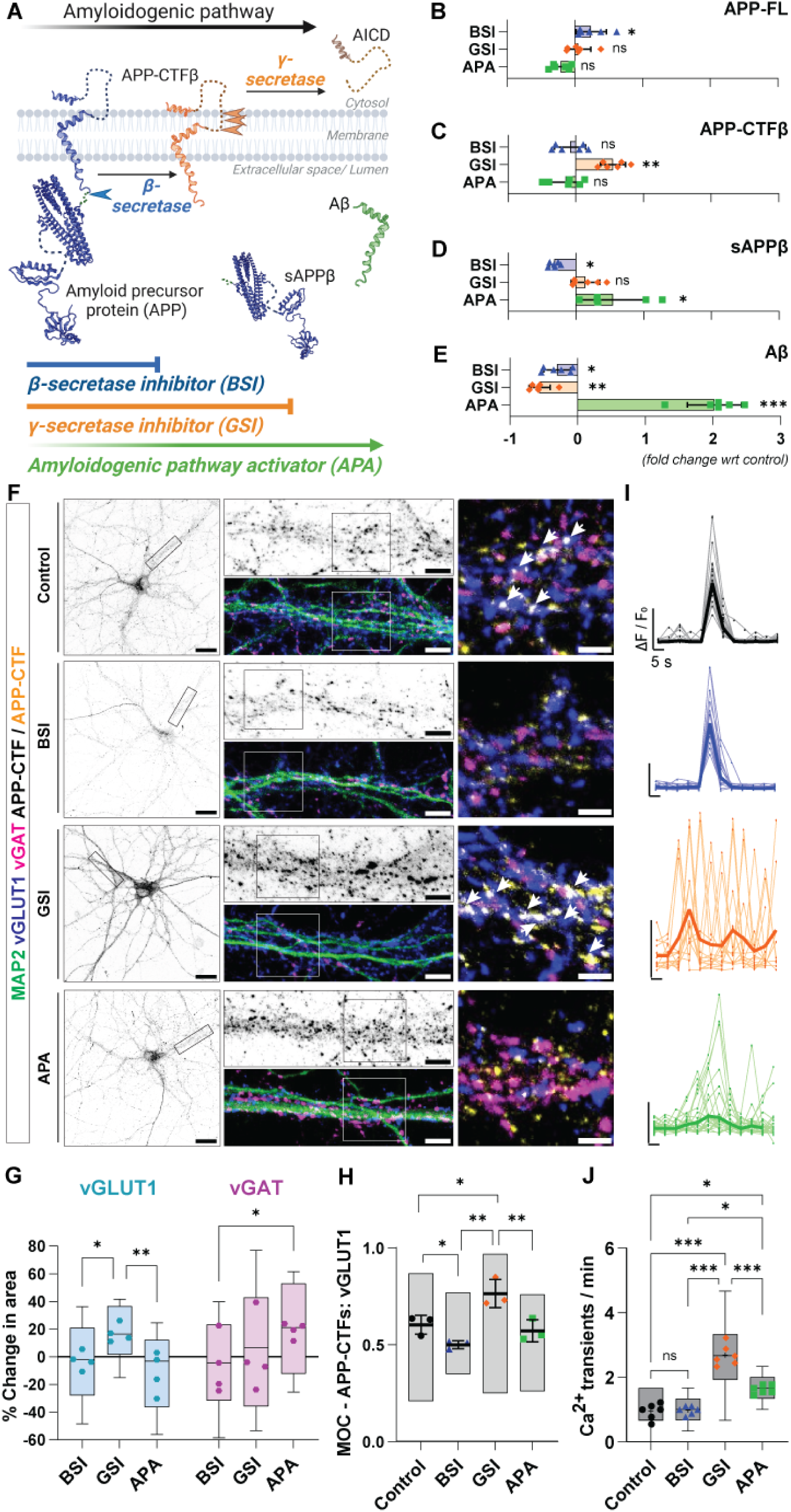
Pharmacological manipulation of amyloidogenic proteolysis has differential impact of synaptic activity. (**A**) Schematic representation of APP proteolysis *via* the amyloidogenic pathway. Initial cleavage by BACE1 results secretion of soluble APP species (sAPPβ) leaving a membrane bound C-terminal fragment (APP-CTFβ). γ-secretase cleavage and sequential trimming of APP-CTFβ additionally releases Aβ peptides in the extracellular space or lumen and intracellular AICD (amyloid intracellular C-terminal domain), respectively. Rat cortical neurons (DIV21) were treated with LY2886721 (1 μM; BACE1 inhibition, BSI), DAPT (10 μM; γ-secretase inhibition, GSI) or Aftin-4 (5 μM; activator of APP processing, APA) for 4 h, to pharmacologically manipulate various stages of APP proteolysis. (**B-E**). Quantitative analyses from western immunoblots (B, C) ELISA (D, E) and depicting fold change in the levels of membrane-bound APP-FL (B), APP-CTFβ (C) and secreted sAPPβ (D), Aβ (E) detected in neuronal homogenate and culture media, respectively. n = 6, N = 3. (**F**) ICC depicting relative changes in the levels of APP-CTFs upon treatment with different compounds (*gray, yellow-merge*). Neurons were also co-stained with vGLUT1 (excitatory synapses, *blue*), vGAT (inhibitory synapses, *magenta*) and MAP2 (dendrites, *green*). Box selection depicts the zoomed in panel. Arrows depict colocalization of APP-CTFs with vGLUT1+ synapse. Scale bar = 1 μm, N = 4. (**G**) Percentage change in vGLUT1 and vGAT area following pharmacological modulation of APP processing was quantified and presented as a box plot. N = 4. (**H**) Scatter plots depict Mander’s overlap coefficient of colocalization between APP-CTFs and vGLUT1 positive boutons. N = 4. (**I-J**) Neurons expressing Camk2-GCaMP6 (AAV transduction, DIV3) were subjected to treatment with different modulators (DIV21, indicated above). Spontaneous calcium transients were recorded in the absence and presence of Ca^2+^ in Tyrode buffer. Ca^2+^ traces per minute have been depicted respectively as (*individual traces-thin lines; average-thick lines*; H). Representative video files have been provided in SI. Box plots (I) depict average number of spontaneously evoked somatic or dendritic Ca^2+^ transients per minute, following pharmacological modulation of APP proteolysis. Each data point indicates an experimental replicate. n = 6; N = 3. ns (*p* > 0.05), * (*p* ≤ 0.05), ** (*p* ≤ 0.01) , *** (*p* ≤ 0.01); One-way ANOVA.

BSI had no strong effect on synapse size but slightly altered calcium transients in individual neurons (Fig. 2F-J, fig. S4A-C). The lower synaptic APP-CTFβ levels correlated with decreased colocalization of APP-CTFβ with vGLUT1 (Fig. 2H; fig. S3J-K). On the other hand, GSI resulted in a roughly 20% increase in excitatory synapse size [vGLUT1 positive] without affecting inhibitory synapses [vGAT positive] (Fig. 2F-G, fig. S3I). To ensure that excitatory synapse enlargement was due to APP-CTFβ accumulation and/or Aβ decrease and not to another substrate of 𝛾𝛾-secretase, we repeated this experiment while knocking down APP using siRNA (fig. S8). The effect was not observed in neurons transfected with siRNA targeting APP indicating that APP-CTFβ accumulation and/or Aβ decrease are driving the increase in excitatory synapse size (fig. S8C, H; GSI alters synapse number in absence of APP). The large vGLUT1 presynaptic boutons observed after GSI had elevated levels of APP-CTFβ when compared with control conditions (Fig. 2F, H; fig. S3J-K). Using puromycin to label ongoing protein synthesis, we also found an increase in protein synthesis after GSI (fig. S5). Since only excitatory synapses were affected by GSI we predicted an E/I imbalance that should drive the network of neurons toward hyperexcitation. Indeed, when recording calcium transients after 4h of GSI, we observed a sharp increase from 1 to almost 3 calcium transients per minute (Fig. 2I-J). While APA still led to a slight increase in calcium transient frequency, mild compared to GSI, we observed no significant change in ΔF/F0 intensity of those transients (Fig. 2I-J, fig. S4C). When compared to GSI, APA had strikingly opposite effects, resulting in an enlargement of inhibitory synapses while leaving excitatory synapses unaffected (Fig. 2F-G, fig. S3I). Altogether, our results suggest an opposite effect on E/I imbalance of GSI and APA. While accumulation of APP-CTFβ in Aβ-reduced conditions leads to hyperexcitability, over-production of Aβ reduces network activity (Video S1).

To dissect the mechanisms by which APP-CTFβ and Aβ affect E/I balance, we conducted live imaging experiments during which we measured in parallel dendritic calcium transients and synaptic vesicle release using SynaptoRed (Fig. 3A, *scheme*, video S2) under stimulation conditions. The entire culture was stimulated using an external electrode at intervals of 30 seconds. Dendritic calcium transient confirmed the general trend of our findings recording spontaneous calcium activity (Fig. 2I, J; fig. S4, video S3). Nonetheless, BSI, which did not affect whole-neuron spontaneous calcium transients, resulted in a sharp reduction in the peak amplitudes (Fig. 3B-C, fig. S6A) and decay-times (Fig. 3D) of individual stimulation-induced transients. Interestingly, under GSI conditions, transients’ decay-time became so slow that calcium concentration appeared not to go back to baseline between two consecutive stimulations (Fig. 3B-C). Since amyloidogenic APP proteolysis appears to take place in excitatory boutons, we investigated the impact of the modulation of APP proteolysis on presynaptic vesicle release. SynaptoRed (SR) integrates into membranes and accumulates into synaptic vesicles (*33*). After it is washed from the cell media and in absence of stimulation, SR fluorescence slowly decays in synapses as labeled synaptic vesicles fuse with the plasma membrane. Under control conditions SR fluorescence decreases drastically immediately after stimulation (Fig. 3A, F-G). Surprisingly, under GSI, SR fluorescence decreased much faster in absence of external stimulation (Fig. 3I), wherein stimulations failed to trigger the drop in SR fluorescence. This indicates an increase in the number of vesicles released in absence of stimulation and a failure of stimulation to induce synchronous release - because the ready-to-be-released pool of synaptic vesicles has been depleted (fig. S6C-D). To confirm the specificity of the effects induced by amyloidogenic APP products, we also treated our culture with alpha secretase modulator GI25423X (ASM) and Notch signaling inhibitor FLI-06 (NSI, fig. S3A-E). The effect on dendritic calcium transients and presynaptic vesicle release (fig. S4D and S6E-I) induced by either treatments could be attributed to alternative signaling pathways (metalloproteinases and notch), plausible independent of APP proteolysis. Additionally, Compound E, another γ-secretase inhibitor (GSI2) showed comparable results to DAPT (GSI), validating our previous observations (Fig. S3; fig. S4D, S6E-I). Furthermore, GSI induced effects on neuronal and synaptic activity were absent in APP deficient neurons (fig. S8D-G). Altogether, this strongly suggests that GSI directly interferes with the synaptic vesicle release. But is it the accumulation of APP-CTFβ or the reduction in Aβ that mediates this effect?

**Figure 3.**
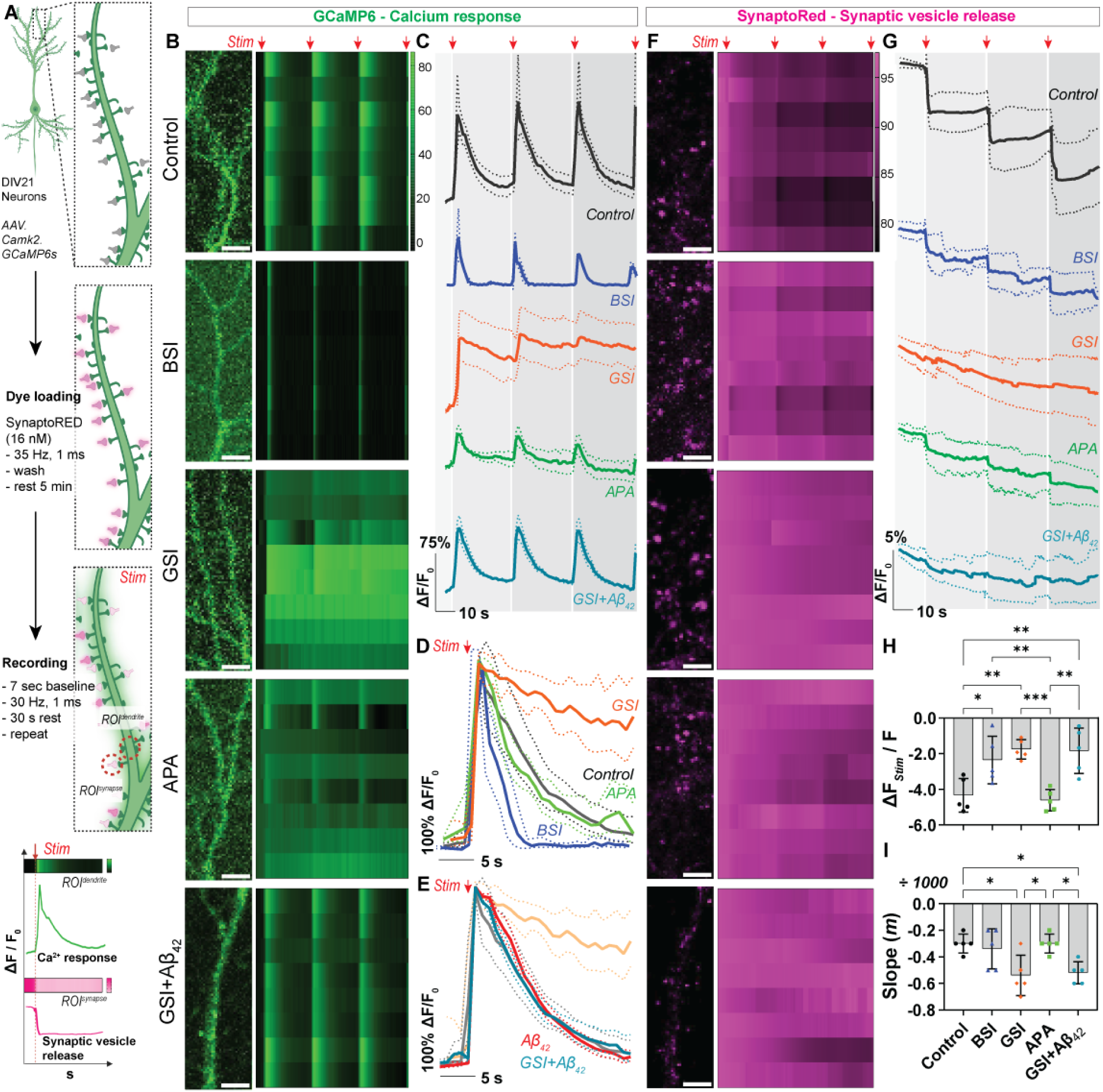
APP and its proteolytic fragments have distinct yet interconnected effects in regulating neuronal network activity. (**A**) Schematic representation depicting experimental flow for dual imaging of neuronal Ca^2+^ response and synaptic vesicle release events. Neurons expressing AAV.Camk2.GCaMP6 (DIV3) were subjected to treatment with different modulators (DIV21). Post treatment synaptic vesicles were loaded with SynaptoRed (SR), excess dye was washed off and neurons were rested for 5 min for stable baseline recordings. Neurons were stimulated with 30 Hz, 4 repetitions, starting 7^th^ s, with 30 s recovery period. Data was recorded for 2 min with 125 ms intervals. (**B-F**) Changes in fluorescence intensity of GCaMP6 (*green*) and SR (*magenta*) were quantified for eight individual ROIs (aligned dendrite, **B** and synapses, **F**), presented as heat map. Overall changes in neuronal Ca^2+^ response (**C-E**) and SR fluorescence intensity (**G***, quantified from multiple selected ROIs*) is depicted as line plots, respectively. Traces depict mean (*bold*) and S.E.M (*dotted lines*), n = 15, N = 5. (**H**) % Decrease in fluorescence intensity of SR (ΔF/F) depicting synaptic vesicle release (just after stimulation) has been shown as a bar graph. n = 15, N = 5. (**I**) SR fluorescence values recorded during the initial 6 s before stimulation was linear fitted, slopes calculated and plotted as bar graphs. n = 15, N = 5. ns (*p* > 0.05), * (*p* ≤ 0.05), ** (*p* ≤ 0.01) , *** (*p* ≤ 0.01); One-way ANOVA.

### Protective role of Aβ

We have shown that after 4 h of GSI both increases APP-CTFβ concentration and decreases the extracellular Aβ (>50%) (Fig. 2B-E). We assessed the effect of Aβ alone and found that it did not alter the intensity of the calcium transients nor affect their shapes (Fig. 3E; fig. S6A,E,G; video S4). To dissect the individual contribution of APP-CTFβ and Aβ, we exposed GSI pre-treated neurons with monomeric Aβ extracellularly for 5 min, before recording again calcium transients and vesicle fusion using SR. Surprisingly, Aβ treatment was able to abolish effects of GSI on calcium transients and rescue neuronal network hyperactivity (Fig. 3B-I; fig. S6A-B). Despite a slight decrease in ΔF/F_0_ intensity all other parameters were comparable to control untreated/Aβ alone treated neurons (Fig. 3B-C, E; fig. S6A, video S4). However, here Aβ did not significantly affect synaptic vesicle defects triggered by APP-CTF accumulation (Fig. 3F-I; fig. S6E, G-I) at the presynapse. These results suggest that Aβ mainly acts post-synaptically. Indeed, many reports have shown that Aβ regulates neuronal activity by binding to postsynaptic glutamate receptors (*34–37*). This means that presynaptic defects induced by accumulation of APP-CTFβ could be masked by postsynaptic counteracting effects of Aβ. Supporting this observations, we could mimic acute Aβ treatment effect on GSI treated neurons by shortly exposing them to CNXQ (AMPA receptor antagonist, fig. S7A) or AP5 (NMDA receptor antagonist, fig. S7B). Of note, this effect was only seen with Aβ_1-42_ species, Aβ_1-40_ and Aβ_42-1_ did not show this effect for the acute treatment timeframe examined in this study (fig. S7C-E). ICC and WB analyses post-acute exposure of Aβ_1-42_ to untreated or GSI pre-treated neurons, showed that Aβ peptides remain bound to the neuronal membranes and negligible signals are seen in the juxtanuclear space, independent to GSI pre-treatment (fig. S7F-G).

### Presynaptic APP-CTFβ interacts with synaptic vesicle release machinery

Experimental evidence from sorted excitatory synapses (Fig. 1F-H, O-P; figs. S2H-O, S3J-K), calcium recordings (Figs. 2I-J, 3B-D; figs. S4, S6) and synaptic activity (Fig. 3F-I; fig. S6) pinpoint that the site of action of APP and APP-CTF is at the presynapse. Thus, our first approach was to identify if there are key differences in the synaptic location of APP-FL and APP-CTFs. We performed immunogold electron microscopy (EM) after live-labelling with antibodies targeting the APP-FL ectodomain and APP-CTFβ epitopes in rat and mouse cortical neuronal cultures (Fig. 4A, fig. S9A). APP-CTFβ was more frequently detected at the presynaptic membrane— particularly within the active zone—compared to full-length APP. This was further validated by ICC of cultured neurons wherein presynaptic APP-CTFs colocalize with the active zone protein RIM1 (regulating synaptic membrane exocytosis protein 1, Fig. 4B). What is the function of APP-CTF at the active zone? Does it interact with other synaptic proteins?

**Figure 4.**
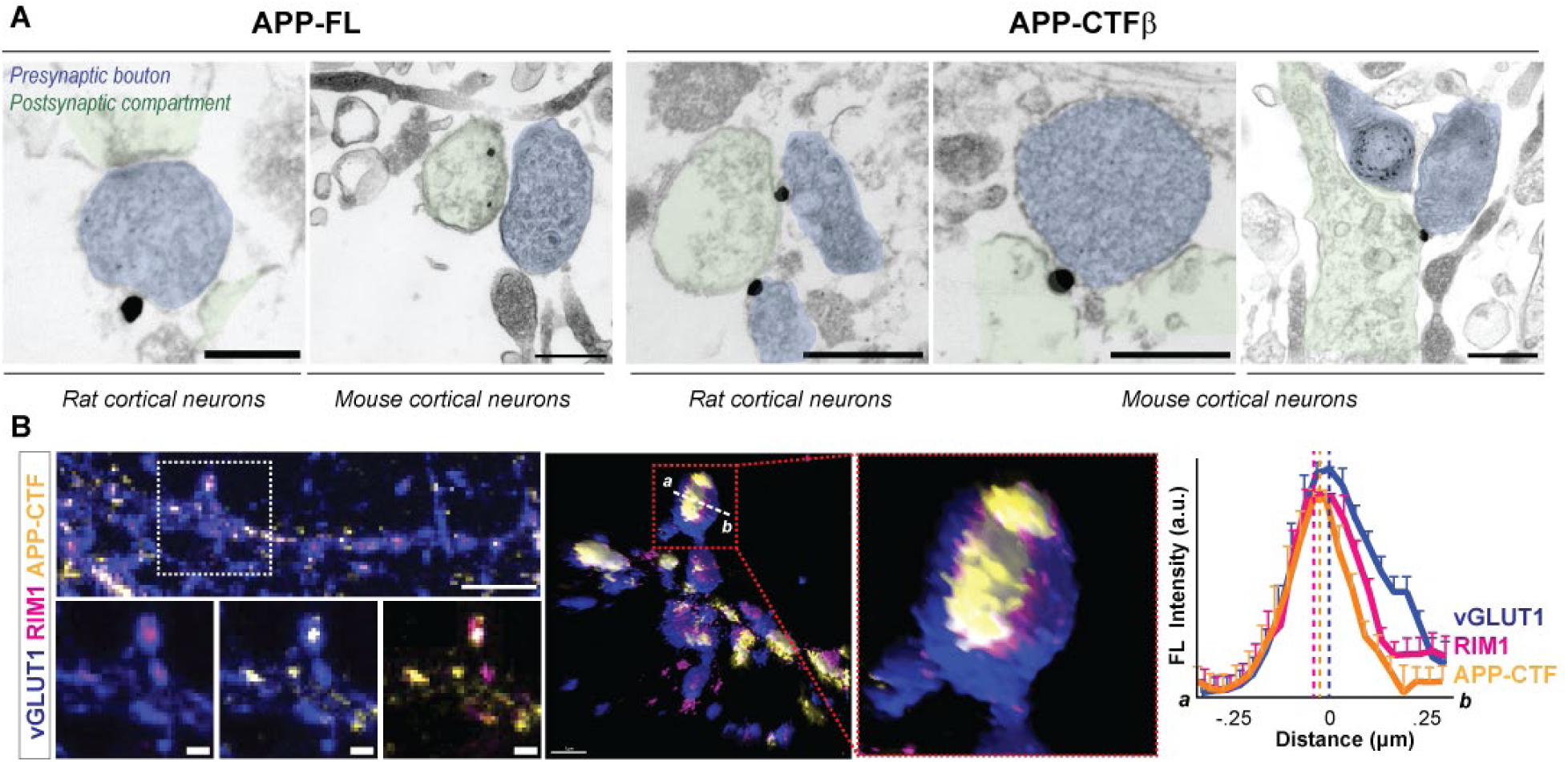
Presynaptic APP-CTF is present in the active zone. (**A**) Immunogold labelling of full-length APP (APP-FL) and C-terminal fragment-β (APP-CTFβ) and EM imaging of synapses in cultured rat and mouse cortical neurons. Presynaptic boutons have been highlighted in blue, wherein their postsynaptic counterpart has been indicated in green. (**B**) Immunocytochemical analysis and Imaris3D reconstruction depicting presynaptic APP-CTF is present in the active zone colocalizes with presynaptic active zone protein RIM1 (inset depicts zoom in, red dotted box). N = 2.

APP has been shown to interact with SYT1 (synaptotagmin 1) *via* its intracellular /luminal C-terminal domain (*29*, *38–41*). Using sandwich ELISA (Fig. 5A-B) and immunoisolation (WB, Fig. 5C) techniques, along with SYT1, we additionally identified STX1 (syntaxin 1), Syb (synaptobrevin 1), SNAP25 (synaptosomal associated protein 25) as interactors of the presynaptic APP intracellular C-terminal domain within synaptosomes isolated from adult rat brain. Since both the full length and APP-CTFs share the same C-terminal domain (Fig. 4C), making them indistinguishable in proteomics and mass spectroscopy methods, we set out to examine if there are any differences between the two, using a dual-immunoisolation based ELISA technique (Fig. 5D, *scheme*; fig. S9B-J). We primarily enriched the full-length protein using an antibody against the ectodomain (APP N-terminal, APP-NT; fig. S9B). The APP-FL depleted material was then enriched for APP-CTFβ (fig. S9B), and the levels of synaptic vesicle proteins were then compared between the two fractions (Fig. 5D, fig. S9C-J). Synaptic vesicle proteins preferentially interact with APP-CTFβ (>1.5-fold enrichment) as compared to the full-length protein.

**Figure 5.**
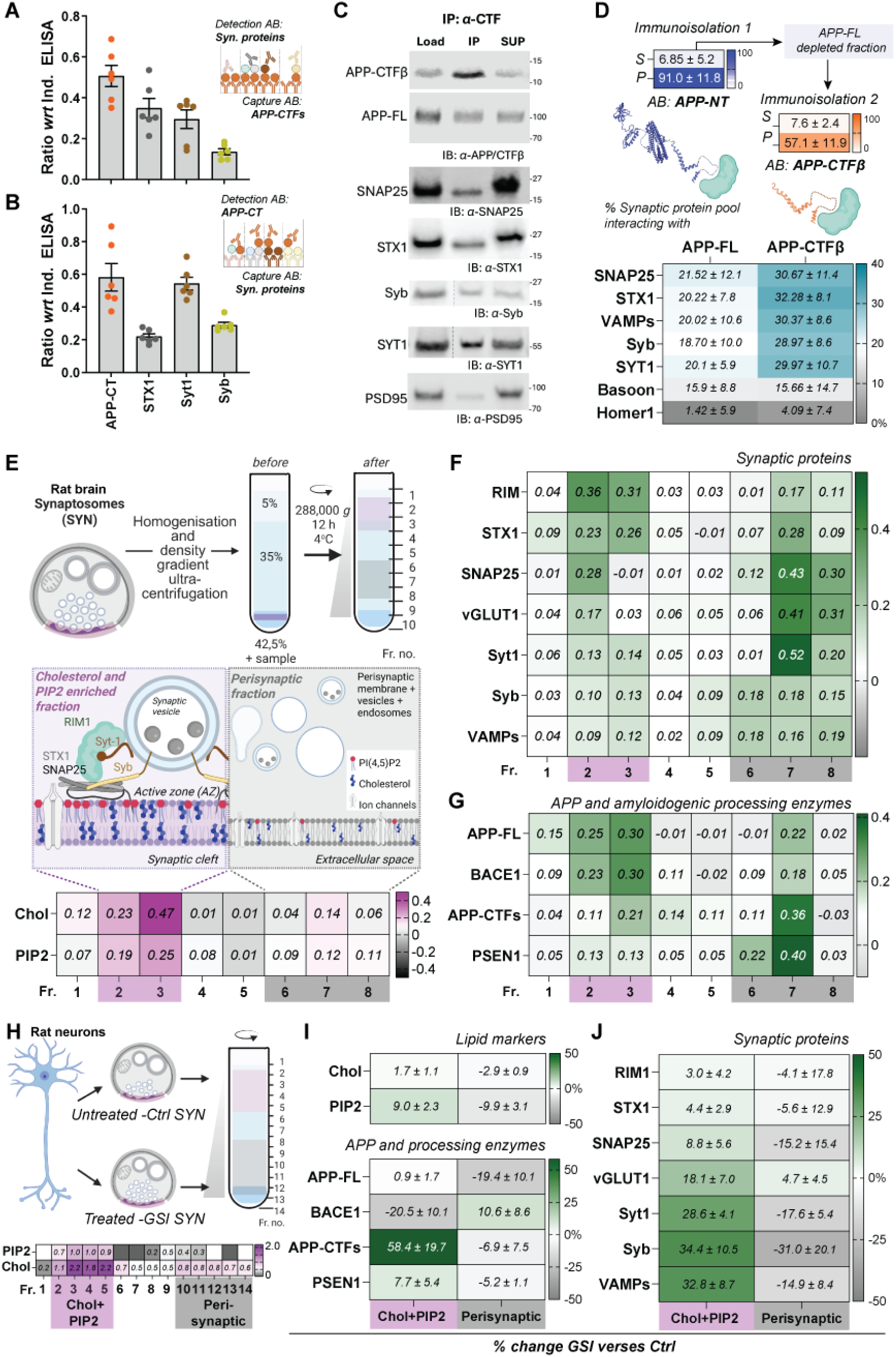
Presynaptic APP-CTFβ interacts with vesicle release machinery and facilitates docking of synaptic vesicles at the active zone. (**A-B**) Quantitative analyses from *in-house* sandwich ELISA showing interaction of APP-C-terminal with different synaptic proteins examined from synaptosome homogenate. n = 6, N = 3. Inset scheme depicts the capture and detection AB used in the experiments. (**C**) Rat synaptosome homogenate was immunoisolated with α-APP-C terminus antibody and interacting proteins were detected *via* western immunoblotting. N = 3. (**D**) ELISA analyses dual immunoisolation from rat synaptosome homogenate, first with α-APP-N terminus antibody and later with α-APP-C terminus antibody depict % relative abundance of synaptic proteins interacting with APP-C terminal domain is significantly enriched with the APP-CTFs rather than the full-length protein. n = 6, N = 3. (**E, H**) Schematic representation depicting fractionation of synaptosomal membranes into cholesterol and PI(4,5)P2 enriched (*highlighted in magenta*) and the perisynaptic membrane fraction (*grays*) from rat brain (**E**) and cultured neurons (**H**). Heat map depicts the normalized levels of individual lipid in each fraction. n = 8, N = 4. (**F-G**) Heat maps depicting relative pool of APP, APP-CTFs and its processing enzymes (**F**) synaptic proteins (**G**) as well as within isolated fractions. n = 8, N = 4. **H-J.** Cultured rat neurons were either treated with GSI (DAPT, 10um, 4 h) or untreated were subjected to synaptosome isolation and lipid fractionation. Collected fractions 1-14 were analyzed for cholesterol and PIP2 content (heat map, **H**). Heat maps (**I-J**) depict percentage change of lipids, APP, APP-CTFs and processing enzymes (**I**) and different synaptic proteins (**J**) in the two distinct pools. n = 4, N = 2. ns (*p* > 0.05), * (*p* ≤ 0.05), ** (*p* ≤ 0.01) , *** (*p* ≤ 0.01); One-way ANOVA.

Does the localization of APP-CTFs at the active zone and its interaction with synaptic vesicle proteins influence vesicle docking? This was examined by homogenizing synaptosomes and separating lipid microdomains (cholesterol and PIP2 enriched fraction) versus the perisynaptic fraction *via* density gradient ultracentrifugation (Fig. 5E). Fractionated samples were validated by examining the levels of cholesterol (Chol) and PIP2 (Fig. 5E) along with FLOT-1 (flotillin) and TFR (transferrin receptors, fig. S9K-L). Enrichment of RIM1 and STX1 within fraction 2-3 indicated that Chol and PIP2 enriched fractions indeed contain active zone membranes. SNAP25, vGLUT1, SYT1, Syb , VAMPs (vesicle-associated membrane proteins) significantly enriched in the perisynaptic fraction indicating the major synaptic vesicle pool as compared to the docked vesicles at the active zone (Fig. 5F). Interestingly, the levels of APP-FL and APP-CTF show differential distribution between the two fractions, this is also comparable to their immediate cleaving enzyme -BACE1 and PSEN, respectively (Fig. 5G). Of note, APP-CTFs are primarily detected in vesicular endo-lysosomal membranes (*13*, *14*), which are enriched within the perisynaptic fraction (Fig. 5G), suggesting a transient stay at the active zone. Next, we isolated synaptosomes from untreated (ctrl) and GSI-treated neurons (Fig. 5H) and used a similar protocol to separate Chol and PIP2-enriched active zone and perisynaptic membrane fractions (Fig. 5H, heat map; fig. S9M). Comparing the protein levels between GSI treated and untreated conditions, we observed an approximately 60% increase in APP-CTFs within the chol+PIP2-enriched fraction, while APP-FL levels remained unchanged (Fig. 5I, fig. S9N), in line with our previous observations (fig. S3J-K). Notably, APP-CTF-interacting synaptic vesicle proteins—Syt1, Syb, and VAMPs—showed a ∼30% increase in the active zone fraction, whereas levels of RIM1, STX1, and SNAP25 were comparable across conditions (<10% change; Fig. 5J, fig. S9O). These findings suggest that accumulation of APP-CTFβ at the active zone enhances the pool of docked synaptic vesicles. Yet, the question remains, could the accumulation of APP-CTFβ directly enhance synaptic vesicle release?

### APP-CTFβ oligomerization increases vesicle release probability

To further expand on the presynaptic role of APP-CTFβ and understand the link between its accumulation at the excitatory synapse and increase in release probability, we modelled APP-CTFβ for the first time in a realistic presynaptic membrane approximating lipid concentrations at presynapse (*42*, *43*) (Fig. 6A, tab. S5, video S5), or POPC membranes, and performed all atom MD simulations for 500 ns. Our primary observations are in line with previously published work wherein the transmembrane domain (TMD, residues 28-52) remain suspended within the membrane bilayer, N-terminal arm remains unstructured, wherein the C-terminal helix dips and remains at the membrane surface (fig. S12) (*44*). Our interest was piqued to notice that Arg76 interacted with PIP2 head groups (stoichiometry of 1:2) -only present in realistic presynaptic membranes-within the first 100 ns of the simulations and remained quite stable through the end of it (Fig. 6B, replicas 2 & 3 in SI, fig. S13). We confirmed the proximity of PIP2 and APP-CTFβ at the active zone (facing the dendritic arm) of vGLUT1 positive excitatory presynapse using co-immunostaining in primary neurons (Fig. 6C). The strength of this interaction was examined by computing the interaction energy indicating the strength of the non-bonded interactions (Fig. 6D). Strong energetic non-covalent bonds are measurable throughout the protein between not bound adjacent residues (amino acid position ±2). Surprisingly, the interaction energy between Arg76 (*yellow circle*) and the two closest PIP2 head groups (*red circles*) is comparable to the ones of intramolecular interaction within APP-CTFβ (fig. S14A). Thus, we hypothesize that the Arg76-PIP2 head group interaction could be relevant in regulating the relative position of the C-terminal helix of APP-CTFβ toward or away from the plasma membrane. This could in turn modulate its binding to synaptic vesicles *via* Synaptotagmin-1 or synaptobrevin as shown previously [(Fig. 4E-H) and (*29*, *38–41*, *45*)].

**Figure 6:**
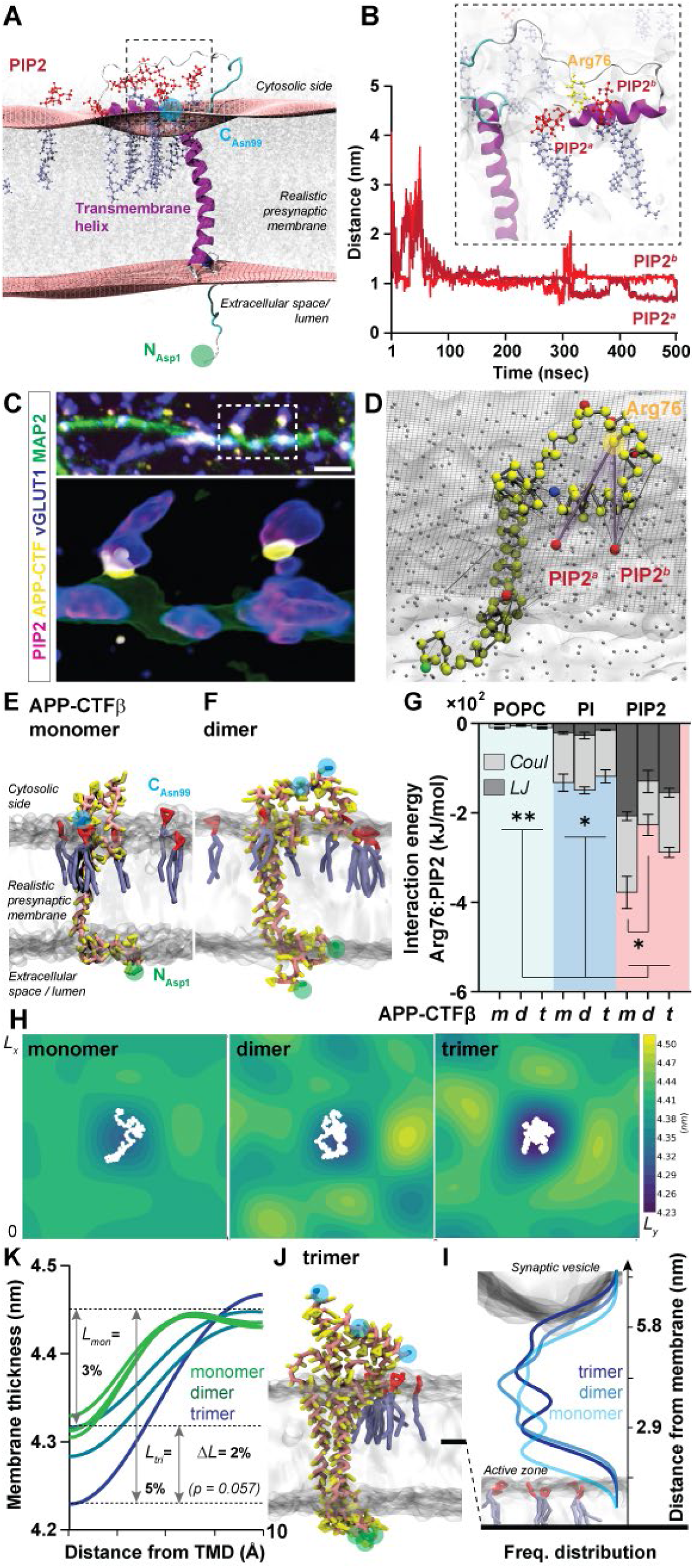
APP-CTFβ interacts with PIP2 and shows propensity of membrane thinning capacity. (**A-B, D**) All atom MD simulation of APP-CTFβ (C99) in a membrane that approximates the realistic presynaptic lipid concentrations (tab. S3; **A**, *snapshot*). The surface of each monolayer was obtained by fitting phosphorus atoms positions of each lipid to an analytical function (Chol not used in the fitting). The protein is shown in secondary structure representation and motifs are colored, respectively. α-helix, *purple;* turns, *cyan;* coils, *white.* N terminal Asp1, *green;* C-terminal Asn99, *blue,* PIP2 head group, *red* and tails, *blue.* (**B**) The distance between PO_4_ headgroup of the two closest PIP2 head groups and the furthest carbon atom (C*Z*) in the side chain of Arg76 as a function of time. The results were consistent across three independent replicas. A simulation snapshot depicting the interaction of PIP2 with Arg76 of APP-CTFβ. (**C**) Immunocytochemical analysis and Imaris3D reconstruction depicting presynaptic APP-CTF (*yellow*) colocalizes with PIP2 (*magenta*). Neurons were co-stained with MAP2 (dendrites, *green*) and vGLUT1 (excitatory presynapse, *blue*). Inset depicts zoom in, *red dotted box*). N = 2. (**D**) Interaction energy network mapped onto the all-atom structure. Every residue is shown as its center of geometry. Width of a connection correlates to the strength of the non-bonded interaction energies. Arg76 (*yellow circle*), two closest PIP2 (*red*) Arg76:PIP2 interaction (*highlighted in light red*). (**E-K**) Coarse-grained MD simulation of 1, 2 or 3 APP-CTFβs in a realistic presynaptic membrane. Renderings of monomer (**E**), dimer (**F**) and trimer (**J**) APP-CTFβ in the elastic network model; protein backbone, *light orange*; side chains, *yellow*; PIP2 head group, *red*; tails, *light blue*; N-Asp1, *green* and C-Asn99 terminal, *blue*, analogous to panel A. (**G**) Total average interaction energy contributions of Arg76 to all lipids of a specific type split into Van der Waals (LJ, Lennard-Jones) and electrostatic (Coulomb) contributions. The contributions have been scaled/averaged by the abundance of the lipids. Note, since some of the PIP2 lipids are located far from Arg76, the interaction energy with the bound PIP2 is higher. Statistical analysis compared between the lipid types, two-way ANOVA, interaction of Arg76 with PIP2, one-way ANOVA, N = 3. * (*p* ≤ 0.05), ** (*p* ≤ 0.01). (**H-I**) Thickness profile of the lipid membrane quantified radially from the APP-CTFβ transmembrane domain (1 protein, n = 3) or cluster (2 protein, n = 2; 3 protein, n = 1). Heat maps (**H**) depict the changes in membrane thickness in 1-3 APP-CTFβ systems, protein depicted in white (BB beads). (**J**) Line plots show the height of the last backbone bead of the C-terminal to/from the center of the membrane leaflet as obtained as a fitting of the PO_4_ headgroup for all simulation models. Distance between the active zone and synaptic vesicle membranes depicted as a cartoon representation. All data are representative of three independent simulations runs (three replica simulations), depicted in SI.

Recently, there have been reports about the propensity of APP-CTFβ to form dimers or oligomers (*44*, *46*, *47*). To examine the behavior of individual domains of APP-CTFβ we additionally employed to coarse-grained MD models using Martini3 force field (*48–50*) to capture a longer time and length scale. To model APP-CTFβ secondary structure we used two popular approaches:

i. the elastic network (*51*) and (ii) the GO models (*52*). For both membrane types described above, we introduced 1, 2 or 3 peptides of APP-CTFβ (SI, figs. S15-S20, Tab, S6) in a single membrane created with TS2CG 2.0 (*53*). During the course of 5 μs simulations we observed the spontaneous formation of dimers and trimers of APP-CTFβ (renderings of mono-, di- and trimers shown in (Fig. 6E, F and J). We computed the interaction energy contributions (*54*) of Arg76 to the different lipid types, i.e., POPC, PI and PIP2 (Fig. 6G), other lipids (figs. S14-15). In line with all atom simulations, we observed for a single motif of APP-CTFβ, that Arg76 made the strongest interaction with PIP2. Notably, dimerization (*p* = 0.034) as well as trimerization of APP-CTFβ decreased this interaction. This reduction in Arg76–PIP2 interactions further influencing the positioning of the C-terminal helix-away from the inner membrane leaflet. Like the all-atom simulations (Fig.5A; phosphorus atoms fitted monolayer, *red*), we noticed that APP-CTFβ induced ∼3% thinning of the membrane bilayer around its transmembrane domain (Fig. 5H–I, figs. S16- 19). This deformation was also observed for dimers of APP-CTFβ but importantly was accentuated when APP-CTFβ formed trimers (*L* = 5%; Fig. 6H-I).

Finally, we examined the distance of the APP-CTFβ C-terminal residue (Asn99) to the middle of the bilayer membrane (Fig. 6K, fig. S20). During our coarse-grained simulations we observed that the C-terminal domains of APP-CTFβ oscillated between locations from within the plasma membrane up to 9 nm from the center of the membrane bilayer -exploring the cytosol. Interestingly, APP-CTFβ C-terminal domains in dimers and trimers are spending more time further away from the membrane, during the simulations; than the ones of monomers, due to the Arg76-PIP2 tether. This suggests that APP-CTFβ oligomerization makes its C-terminus more prone to recruit docked synaptic vesicles. Indeed, the distance separating docked vesicle membrane to the surface of the inner side of the plasma membrane is approximately 5.8 nm (*55*) - exactly in the range of APP-CTFβ C-terminus cytosolic exploration (Fig. 6K). Both capacities of APP-CTFβ oligomers to thin the plasma membrane and to favor the C-terminal helix cytosolic exploration will positively impact synaptic vesicles fusion. Thus, molecular crowding of APP-CTFβ at the active zone - as in GSI induced increase in local concentration-could reduce the energy barrier for synaptic fusion as the spontaneous formation of oligomers by recruiting docked synaptic vesicle to active sites where the plasma membrane thickness is further reduced.

## Discussion

Aβ, in its oligomeric form, has been shown to have toxic effects on neuronal and synaptic function (*56*, *57*). Notably, the soluble Aβ oligomers can bind to excitatory synapses/neurons (*58–60*) and induce neuronal hyperexcitability, a phenomenon observed both in animal models and early-stage AD patients (*61–65*). This aberrant increase in neuronal activity contributes to network dysfunction and impaired synaptic plasticity (*66*, *67*). These events furthermore drive Aβ production (*62*, *63*) and exacerbate tau pathology (*68*, *69*), creating a vicious cycle of neurotoxicity, neuronal loss and neuroinflammation (*65*, *70*). Nonetheless, given the polymorphic nature of AD pathology, is Aβ the earliest and solo toxic trigger of the disease? To address this question, we decided to delineate the roles of APP and its proteolytic fragments, particularly APP-CTFs/β, in synaptic and neuronal function, and its extension to its defects in early AD and disease onset.

We show that in endogenous conditions, APP and its amyloidogenic processing components are predominantly localized to excitatory presynaptic terminals. Acute pharmacological modulation of APP proteolysis results in synaptic APP-CTFβ accumulation driving excitatory synapse enlargement, spontaneous neurotransmitter release, and neuronal network hyperexcitability. While Aβ overproduction modulated network activity, it suggests a distinct compensatory role in maintaining excitatory/inhibitory balance. Our results from acute treatment could be back-extrapolated to literature published evidences of chronic GSI treatment (*71*), familial mutations of APP (*72*, *73*) , BACE1 (*74–76*) and PSEN1 (*77*, *78*) in disrupting synaptic function. Now, the retention of AICD within membrane-bound APP-CTFβ at excitatory synapses (GSI treatment) — rather than its nuclear translocation (APA condition)—may differentially regulate translation aspects, which could contribute to the enlargement of synapse size. These observations pave the way to a non-canonical, local translation-linked function of AICD (*79*, *80*) in early synaptic remodeling during AD onset, which should be investigated in the future.

One of the key observations from our study is that APP-CTFs/β localized at presynaptic active zones, interact with components of the synaptic machinery and local lipids. Previous studies have reported binding between APP C-terminal and synaptic vesicle protein-synaptotagmin using high end quantitative mass spectrometry and proteomic approaches (*29*, *38–41*, *45*). However, these approaches are not able to unambiguously distinguish APP-CTFs from APP full length and as they share the same protein sequence (APP-CTFs lack the large N-terminal ectodomain of APP full length). Thus, we used more classical biochemical approaches making use of antibodies recognizing specifically the two proteins. Indeed, we could show that in brain tissue the interaction with Synaptotagmin1 is mainly driven by the C-terminal domain of APP-CTFβ, rather than the full length APP employing simpler ELISA based immunoisolation and western blotting. Could it be that APP N-terminal extracellular domain prevents full length APP from entering synaptic clefts? Of note, the APP N-terminal domain has two glycosylation sites enhancing even more its apparent bulkiness.

Strikingly, when we treated living neurons with an antibody targeting the short extracellular sequence of APP-CTFβ (AB#1), we observed a rapid (few min) silencing of neuronal and dendritic calcium traces and a sharp decrease in SR fluorescence responses (fig. S10A, *scheme* and S10E). This effect was partially conserved when using an antibody recognizing both APP full-length and APP-CTFβ (AB#2). However, incubation with an antibody targeting the ectodomain of full-length APP protein had no effect (fig. S10A-D).

These observations further warrant in-depth investigation of APP C-terminal dependent –mediated calcium signaling events *via* Syt1 and/or GTP-binding protein G(o) (*81*, *82*), that modulate release probability. In our simulations, we uncovered a strong interaction between Arg76 of APP-CTFβ and PIP2 lipid head group. This interaction sequesters the C-terminal helix of APP-CTFβ in the plasma membrane, preventing its binding to Syt1. Accumulation of APP-CTFβ facilitates the formation of dimers or trimers which in turn decrease its interaction with the PIP2 head group, favoring its interaction with synaptic vesicle proteins. A second key finding of our study is the impact of APP-CTFβ on synaptic membrane thickness. Using coarse-grained simulations, we showed that APP-CTFβ spontaneously oligomerizes *in silico*. Trimers of APP-CTFβ induce a local thinning of the plasma membrane. This could lead to a decreased energy barrier for synaptic vesicle fusion in the region. Together, the interaction of APP-CTFβ with the synaptic vesicles and its oligomerization seem critical in regulating neurotransmitter release. These findings could be extrapolated to literature reports highlighting BACE1 activity being essential for optimal release of synaptic vesicles, and its deficiency or pharmacological inhibition decreases the number of docked synaptic vesicles at the active zones (*74*, *75*, *83*).

In the context of early synaptic deficits observed in AD, altered APP processing due to either increased β-secretase activity or γ-secretase deficits could result in APP-CTFβ accumulation, contributing to the impairment of synaptic vesicle fusion dynamics; preceding the more widely recognized toxicity associated with Aβ species. Supporting our findings, both APP and PSEN1 FAD mutations that result in loss of function of γ-secretase (*i.e.* stalling of PSEN-APP-CTFβ complexes and/or membrane retention of longer forms of Aβ) show augmented synaptopathy and neurodegeneration phenotypes, triggering early onset of AD (*15*, *71*). Multiple reports have shown that during ageing (a risk factor implicated in AD), amyloidogenic APP proteolysis is augmented *via* increase in β-secretase activity, this increases APP-CTFβ accumulation within neuronal sub-compartments, disrupting neuronal activity (*84–86*). Furthermore, recent evidence has shown that APP-CTFβ alone, in the absence of Aβ, is sufficient to induce neuronal dyshomeostasis (*9–15*). Thus, one could speculate that these effects are independent of extracellular Aβ peptides. Taken together, we propose that APP-CTFβ, rather than Aβ alone, is central to early synaptic disruptions, an understudied avenue in the pathophysiology of AD.

While much of the AD research has been focused on Aβ species, our findings highlight the significant contributions of presynaptic APP-CTFβ to regulate synaptic activity and trigger its dysfunction. Now, acute treatment of Aβ (5 min), particularly the 42 variant, rescues GSI induced network hyperactivity *via* postsynaptic mechanisms. While others have reported synaptogenic properties of Aβ (or APP amyloidogenic proteolytic products) (*87*, *88*); literature evidences support the fact that there are beneficial effects of lower concentrations of Aβ in maintaining neuron/synapse health (*88–92*). The functional transition of Aβ from physiological to pathological states is concentration dependent, wherein subtle changes in Aβ levels potentially tips the balance between normal synaptic function to neurodegeneration. These results challenge the prevailing Aβ-centric perspective and underscore the necessity of expanding our focus to encompass the broader spectrum of APP processing products, which may have substantial implications for both synaptic physiology and disease progression, as also reported recently (*93–95*). To further substantiate this we also remark that the balance of α- or β-secretase dependent processing of APP can either offer protection (e.g. Icelandic mutation) or drive the disease pathogenesis (e.g. Swedish mutation), suggesting that alterations in APP metabolism rather than Aβ, triggers AD onset (*72*, *96–99*). Indeed, upon careful screening of literature we find evidences of APP-CTFβ accumulation with synaptic structures/neurons in both transgenic mouse models (*11*, *47*) as well as human AD patients (*100–102*).

Our study also raises a concern about current therapeutic approaches targeting monomeric Aβ clearance in the early stages of AD. While Aβ targeted antibody treatments have shown to successfully reduce amyloid burden, their impact on improving cognitive performance in patients has been less clear (*23*, *25*, *103–107*). Yet, authorities are approving testing those treatments in patients with mild dementia. According to our results, antibody-based clearing techniques of monomeric/unaggregated Aβ could exacerbate synaptic dysfunction driven by APP-CTFs in the early stages of AD. Given the integral role of APP in regulating synaptic function, therapeutic strategies that selectively clear Aβ may be misguided. One could also speculate this to be a major drawback overlooked in the repeated failures of γ-secretase inhibitor-focused clinical trials (*108*, *109*). Our study underscores the need for a paradigm shift in the field of Alzheimer’s disease research. Effective therapies must consider the physiological role of APP processing when targeting its components. Importantly, a more holistic approach that considers the role of both Aβ and APP-CTFβ at the distinct stages of AD would lead to treatments that preserve and restore synaptic function.

## Methods

### Synaptosome isolation

Animals were handled and maintained according to the guidelines laid down by the Animal Welfare Body (AWB) (Instantie voor Dierenwelzijn IvD) in line with the animal experimentation policy within Radboud University and RadboudUMC; under the license/protocol numbers 2021-0040-001/002 to Dr. Anne-Sophie Hafner. Synaptosomes were enriched from forebrains of 1-year-old wild-type and vGLUT1^GFP^ knock-in mice. To favor the isolation of synaptosomes with presynaptic compartments with closed membrane bilayers and postsynaptic compartments with open membrane bilayers (*110*), we used the adapted protocol, previously described (*32*, *111*). Forebrain from a single mouse were homogenized in 2 ml of ice-cold homogenization buffer [0.32 M sucrose, 4 mM HEPES (pH 7.4), EGTA-free protease inhibitor cocktail (Calbiochem, 1:1000), and RNAase inhibitor (Promega,1:1000)] by using a 2-ml glass-Teflon homogenizer with 15-17 gentle strokes. An additional 2 ml of buffer was used for rinsing, and the combined 4 ml was centrifuged at 1000 × g for 8 min at 4°C. The supernatant (S1) was centrifuged against 12,500 × g for 15 min at 4°C. The synaptoneurosome-enriched pellet (P2) was then resuspended in 1 ml of homogenization buffer. This fraction was finally layered on top of a two-step sucrose density gradient (5 ml of 1.2 M sucrose and 5 ml of 0.8 M sucrose, 4 mM HEPES, and EGTA-free protease inhibitor cocktail, as described above). The gradient was ultracentrifuged at 40,000 × g for 70 min at 4°C. Synaptosome fractions were recovered through the tube wall, at the interface of 0.8 and 1.2 M sucrose, by using a syringe to minimize contamination with lighter fractions enriched in myelin. The resulting fraction is referred to as presorted synaptosomes (or presort. SYN).

### Fluorescence-activated synaptosome sorting (FASS)

Synaptosome sorting was performed as previously described (*32*, *111*). The Cytek® AuroraCS was operated using a 70-mm nozzle. Briefly, presort. SYN were stored on ice, diluted in PBS containing protease and RNAase inhibitor as described above. Dilution was optimized to obtain an event rate of 15,000 to 20,000 events/s with an optimal flow rate to maintain single particle per droplet. A threshold was set on SSC-A to discriminate against the noise. A dilution range was made to confirm the correct settings. A first gate delineated small particles (“singlets”) and excluded events showing correlated high values for forward scatter and side scatter areas (aggregates and large particles). The singlets gate was sub gated according to vGLUT1^GFP^ fluorescence intensity by using the 488-laser line. Thus, singlets were sorted into two fractions, the vGLUT1^GFP^-negative (GFP −) fraction and the vGLUT1^GFP^-positive (GFP+) fraction (fig. S2A*, scheme*). These two fractions were either collected on to filters and processed for homogenate preparation or plated onto gelatinized coverslips at a density of 1 Mio particles per 12-mm coverslip in 24-well plates by centrifugation at 5,000 × g for 30 min and then processed for ICC as described ahead.

### Cultured neurons

Dissociated primary rat cortical neuron cultures were prepared and maintained as described previously. Briefly, cortices were dissected from embryonic (day 18) rat pups of either sex (Sprague-Dawley strain; Janvier), trypsinised (1x, 15-20 min), mechanically dissociated in 1% normal horse serum (NHS; ThermoFischer Scientific, #16050130) containing Basal Media Eagle (Life technologies, #21010046), containing supplements like B27 plus (Gibco, #A3582801), penicillin-streptomycin (1x PS, Gibco, #15140122), non-essential amino acids (1x NEAA, ThermoFischer, #11140050), sodium pyruvate (1x, NaPy, ThermoFischer, #11360070). Dissociated cells were plated on poly-D-lysine coated (Sigma Aldrich, #A-003-E) glass coverslips (VWR, #631-1577/ 79/ 84) or 35 mm glass bottom dishes (Mattek, #P35G-1.5-14-C-HA) at specific density to achieve optimal neuron network connections. Neurons were maintained and matured in a humidified atmosphere at 37°C and 5% CO_2_ in growth medium (described above, without serum). Neurons were fed by replacing 1:1 media replacement with BrainPhys™ neuronal medium (StemCell technologies, # 05790) supplemented with NeuroCult™ SM1 neuronal supplement (StemCell technologies, # 05711) and PS (1x) starting DIV4 until DIV21-22 to ensure synapse maturation. All experiments complied with national animal care guidelines and the guidelines issued by the Radboud UMC, Nijmegen and were approved by local authorities (mentioned earlier). For transduction, 3-5-DIV neurons were transfected with AAV-Camk2-GCaMP6s, and transfected cells were maintained until 18 to 21 DIV for further experiments.

### Pharmacological treatments

Primary cortical neurons (DIV21-22) were treated with LY2886721 (1 μM; BSI; Cayman chemicals, #21599-1), DAPT (10 μM; GSI; Cayman chemicals, #13197-5) or Aftin-4 (5 μM; APA; Cayman chemicals, #27341-1) for 4 h in culture media, to manipulate different stages of APP proteolysis. Post-treatment neurons were either fixed with 4% PFA+4% sucrose in PBS for 15 min at RT for ICC or subjected to synaptoneurosomes isolation or lysate/ homogenate preparation, respectively. Treatment with Aβ peptides (100 nM, 5 min) was done following with or without prior GSI treatment and excess peptides were washed with PBS (thrice) followed by live imaging or fixation for ICC.

### Immunocytochemistry

Neurons after fixation were permeabilized with permeabilization buffer (0.025% Triton X-100 (Merck, #T8787-250ML) in blocking buffer) for 2 min, followed by blocking (2.5% NHS, 2.5% bovine serum albumin (BSA; Carl Roth, #8076-4), 0.0125% Triton X-100 in 1x PBS) for 1 h at RT. An antigen retrieval step was performed to augment and facilitate binding of APP-CT/ APP antibodies. Incubation with primary antibodies (respective dilutions in blocking buffer, tab. S2) was performed overnight at 4°C. Next day, coverslips were washed thrice with PBS and incubated with respective secondary antibodies (respective dilutions in blocking buffer, tab. S2) for 2 h at RT. Removal of the secondary antibody solution was followed by washes of PBS (thrice) and distilled water (once) and coverslips were mounted on the slides using Moviol mounting reagent (41.67% Glycerol, 16.67% Moviol 4-88 (CarlRoth# 0713.1) in ddH₂O).

### ELISA

Equal amounts of protein from various brain, synaptosome or cellular fractions (based on antibody LOD quantifications, tab. S4) were added to Nunc® MaxiSorp™ 384 well plates (Sigma Aldrich, #P6366) and incubated overnight at 4°C. Plates were washed, blocked with BSA, and incubated with primary antibodies (dilutions mentioned in tab. S2) at room temperature or 4°C. After washing, plates were incubated with respective HRP-conjugated secondary antibodies, followed by TMB substrate development (ThermoFischer, #34029) . The reaction was stopped with sulfuric acid, and the absorbance was measured at 450 nm with background correction at 620 nm. Samples were analyzed in technical duplicates or triplicates. Detailed experimental protocols for sandwich or dual immunoisolation ELISA, and signal amplification using a tyramide-based system, please refer to the SI file.

### Western blotting

Equal amount of protein (10-20 μg) was loaded protein per lane, except Fig. 1B (S1, P2, presort. SYN = 20 μg, Sort. SYN = 1 μg). Briefly, proteins were separated on a precast 4–15% or 4-20% NuPage^Ⓡ^ Bis-Tris gels (Bio-Rad, #4561083EDU, 4561096EDU), and electro-transferred onto a 0.2-μm nitrocellulose (NT) membrane (Amersham, #1060001) for 10 min using the quick transfer protocol (Trans-Blot turbo transfer system, Bio-Rad). The blots were then incubated in 5% skim milk (Cal Roth, #T145-1) in TBS-T containing 0.1% Tween 20 (ThermoFischer Scientific, #J20605.AP)) for 1 h at RT. Blots were incubated with primary antibodies (overnight at 4°C); followed by respective secondary antibodies (2 h at RT). Antibody dilutions have been summarized in tab. S2. Proteins were detected either by ECL imager (Bio-Rad, Germany) or fluorescent imager(LiCOR Biosciences, Germany). Detailed descriptions have been provided in the SI. Values from each sample were normalized to the values from the control cells in that experiment and fold-change with respect to control was indicated as graphs.

### Microscopy and image processing

Confocal microscopy images were acquired on Nikon live-cell SoRa spinning disk super-resolution microscope (Nikon technologies, Japan). or Leica SP8 Liachroic confocal microscope (Germany) or Leica Widefield fluorescence microscope equipped with Thunder computational clearing module (Germany). Imaging parameters have been mentioned in detail in the SI file.

### Calcium and SynaptoRed imaging

Neurons expressing Camk2-GCaMP6 (AAV transduction, DIV3) were subjected to treatment with different modulators (DIV21, as indicated previously). Spontaneous calcium transients were recorded in the absence and presence of Ca^2+^ in Tyrode buffer for 5 min continuously in 500 ms intervals. ROIs were either derived from the complete neuron (Fig. 2I-J) or segmented to differentiate between soma and dendrite (SI, fig. S2A-B), and Ca^2+^ traces per minute have been depicted to show the number of spontaneously evoked Ca^2+^ transients per minute. Dual imaging of neuronal Ca^2+^ response and synaptic vesicle release events by SynaptoRed C2 (Sigma Aldrich, #S6689-5MG, 16 nM in culture media), wherein synaptic vesicles were loaded with the dye *via* stimulation at 35 Hz, excess dye was washed off with fresh media thrice and neurons were rested for 5 min to ensure stable baseline recordings. This was quantified in a pilot experiment and thus the same timeline was maintained for the entirety of the experiments henceforth. Calcium and synaptic vesicle dynamics were imaged and recorded during the stimulation protocol with 30 Hz, 4 repetitions, starting 7^th^ s, with 30 s recovery period. Data was recorded for 2 min at every 125 ms intervals. Time lapse images were then processed to either quantify the total Ca^2+^ response (Fig. 3C) or just dendrite (ROIs, Fig. 3B). Fluorescence intensities of SR loaded within synaptic vesicles were then quantified for the specific dendritic ROI (ROIs, Fig. 3F) or just complete frame (Fig. 3G). Detailed analysis has been explained in video 2, and representative data has been depicted in videos 2-3.

### Aβ treatment

Untreated or γ-secretase inhibitor (GSI) pretreated neurons were treated with or without synthetic Aβ peptides (100 nM) for 5 min at 37°C. Following incubation, cells were washed three times with pre-warmed culture medium to get rid of excess Aβ and used for live imaging experiments as described earlier. Preparation of monomeric Aβ peptides is described in the SI, along with other experimental protocols.

### Electron microscopy

Neurons cultured on glass-bottom dishes (Mattek, #P35G-1.5-14-C-HA) were live-labeled with or without an antibody targeting the extracellular N-terminal domain of APP-CTFβ (AB#1) or full length APP (AB#2/3) then washed and fixed with PFA (Electron Microscopy Sciences #157-4). After blocking, cells were incubated with a secondary gold-conjugated antibody (1:50; Aurion, #800.022/11), followed by a second fixation with glutaraldehyde (Carl Roth #4157.1). Excess reagents were washed off, and samples were quenched using sodium borohydride and glycine/ammonium chloride. After additional washes, silver enhancement (Aurion, #500.044) was performed. Cells were then osmicated, stained with uranyl acetate, and dehydrated through graded ethanol. Resin infiltration and polymerization followed. Blocks were detached from dishes, trimmed, and ultrathin-sectioned. Sections were post-stained with uranyl acetate and lead citrate, then imaged using STEM mode on a Zeiss Crossbeam 550. Multiple regions per grid were imaged to ensure unbiased representation. Detailed experimental protocol and additional information have been summarized in the SI. Images were minimally processed using Fiji/ImageJ.

### Synaptosome lipid and protein fractionation

Synaptosome fraction (tab. S3, adult rat brains or cultured primary neurons) was homogenized in SynPer reagent (without any surfactant) by multiple freeze thaw cycles to maintain protein-protein interaction complexes. Samples were then loaded on a density gradient (rat brain, 5% - 2.5 ml, 35% - 6 ml and 42.5% 1.5 ml; cultured neurons, 5% - 1 ml, 35% - 3 ml, 42.5% 1 ml and 60% - 6 ml sucrose in HEPES buffer) ultracentrifugation (288,000 × *g*, 12 h, 4°C) to fractionate synaptic membranes into cholesterol and PI(4,5)P2 enriched and the perisynaptic membrane fraction. Next day, 10- 1 ml or 14- 0.3 ml fractions were collected, and the material was subjected to ELISA analysis to examine the levels of different proteins and PI(4,5)P2 lipids. Cholesterol in each fraction was analyzed using Amplex™ Red Cholesterol Assay Kit (ThermoFischer Scientific, #A12216) as per manufacturer’s protocol without modifications.

### Immunoisolation

Synaptosomes (tab. S3, rat brains) were homogenized in SYNPER protein extraction reagent containing protease and phosphatase inhibitors by repeated suspension through a 23G x 1” needle. Synaptosome homogenate was then used for IP experiments using magnetic Sepharose A/G beads (Sigma Aldrich, #GE28-9440-06/). Beads were first coated with respective antibodies for 1 h at RT (**t**ab. S1), washed 3x in PBS and then incubated with material at 4°C for 16 h. Beads were further pelleted down and boiled in 1% SDS + 50 mM Tris + sample buffer, used as *IP eluate* and the remaining supernatant was loaded as *sup*. Beads incubated only with the homogenate was used as the *bead control*. The supernatants exposed to beads only were devoid of non-specific binding substrates and was used further as *loading control*. Similarly, dual immunoisolation protocol, primarily with APP-N terminal ectodomain antibody to deplete FL-APP, and secondly with an APP-CT antibody to enrich APP-CTF pool). Protein-protein interactions to distinguish relative pool of synaptic proteins interacting with full length APP and APP-CTFβ was done using ELISA.

### Computational studies

The protein structure was obtained from an earlier study by Pantelopulos *et al.* (*44, 112*), made available through the author’s GitHub (*113*). The prepared initial structure (modifications explained in SI, initial structure and coarse-graining & fig. S11) was used to prepare three different simulation setups [All atom: realistic and POPC membrane with a single APP-CTFβ (runs #1-3), total-6 simulations; Coarse-grained: realistic and POPC membrane with 1x, 2x, and 3x APP-CTFβ each, in two models each, (runs #1-3), total: 36 simulations]. An all-atom model and two coarse-grained simulations that differ in their choice of modeling the secondary structure of the proteins. Detailed information on modelling parameters, analysis, and calculations etc. are described in detail in the SI.

### Statistical analyses

All graphical illustrations have been prepared using GraphPad Prism 10.4.1.627. Statistical analysis for a particular graph has been mentioned in the respective figure legends. All tests have been performed using the in-built default settings of the module in the software, without modifications, until mentioned otherwise; not significant (ns) *p* > 0.05; * *p* = 0.05; ** *p* = 0.01; *** *p* = 0.001.

## Supporting information

Kapadia et al_SI_Video files

## Acknowledgments

We would like to thank all our lab members for fruitful discussions and helpful feedback. The authors thank Dr. Eric Jansen for sharing AAV.CamK2.GCaMP6; Dr.Thessa.Koorenhof-Scheele and the Radboud Technology Centre Flow cytometry for assistance with FASS; Dr. Jelle Postma for assistance with Imaris; Dr. Hannes Berkhart and Pia Strausberg for assistance with immunogold EM imaging and instrumentation provided by Deutsche Forschungsgemeinschaft, project no. 388171357 (Electron microscope). The authors acknowledge Prof. John Straub (Boston University, Massachusetts) for kindly sharing the *in-silico* structure of APP-CTFβ (C99). This study was supported by the Donders Institute for Brain, Cognition and Behavior and Faculty of Science, Radboud University Nijmegen Netherlands. Schematics were created using Biorender.com.

## Funding

European Research Council (ERC) under the European Union’s Horizon 2020 research and innovation programme - ‘MemCode’, grant 101076961 (AH)

Novo Nordisk Foundation grant NNF18SA0035142 and NNF22OC0079182 (WP)

Independent Research Fund Denmark grant 10.46540/2064-00032B (WP)

Dr.phil. Ragna Rask-Nielsens Grundforskningsfond (FS)

## Author contributions

Conceptualization: AK, ASH

Methodology: AK, FS, ED, JW, IL, NR, WP, ASH

Investigation: AK, FS, ED, IL, NR

Visualization: AK, FS, ED, IL, NR

Funding acquisition: ASH, WP

Project administration: ASH

Supervision: ASH, WP

Writing – original draft: AK, ASH

Writing – review & editing: AK, FS, ED, JW, WP, ASH

## Data and materials availability

All experimental data are available in the main text or the supplementary materials. Computational simulation data, code and materials used in the analysis are deposited in a public database.

**Supplementary Materials**

Materials and Methods

Fig. S1 to S20

Tab. S1 to S4

References (*1–19*)

Movies S1 to S5

Data file 1

