## Supplementary Material file for "Amyloidogenic proteolysis of APP regulates glutamatergic presynaptic function"

**The PDF file includes:**

Materials and Methods

Figs. S1 to S20

Tables S1 to S4

**Other Supplementary Materials for this manuscript include the following:**

Video S1 to S5
Data S1

### Materials

**Table S1. Details of all antibodies used in this study.**

| Target of interest | Specificity (Antibody clone) | Make (Cat. No.) | WB | IHC/ICC | ELISA |
|---|---|---|---|---|---|
| **Amyloid precursor protein (APP) and APP C-terminal fragments** | N-terminal specific | Sigma Aldrich (SAB4200536) | | 1:250 | 1:100 |
| | C-terminal specific (M 3.2) | Covance (SIG-39155) | 1:1000 | 1:500 | 1:200 |
| | C-terminal fragment specific (C1/6.1) | Covance (SIG-39152) | 1:1000 | 1:500 | 1:200 |
| | C-terminal fragment specific (3125) | Generated by Walter lab (Eurogentec)[‡] | 1:1000 | 1:250 | |
| | C-terminal fragment specific (140-rb) | Generated by Walter lab (Eurogentec)[‡] | 1:500 | 1:100 | |
| **Soluble APP species** | sAPPα | Covance (SIG-39139) | | | 1:250 |
| | sAPPβ | Covance (SIG-39138) | | | 1:250 |
| **Amyloid β (Aβ)** | $A\beta_{1-x}$ (82E1) | IBL Int. (JP10323) | 1:500 | 1:200 | |
| | 82E1-Biotin conjugated | IBL Int. (JP10326) | | | 1:1000 |
| | rAβ | Abcam (# ab14220) | 1:500 | 1:250 | 1:1000 |
| | $A\beta_{x-40}$ (BAP-29)[‡] | (Brockhaus et al. 1998)[‡] | | | 1:250 |
| | $A\beta_{x-42}$ (BAP-15)[‡] | (Brockhaus et al. 1998)[‡] | | | 1:100 |

| | | | | | |
|---|---|---|---|---|---|
| | hAβ (2964) | Generated by Walter lab (Eurogentec)[‡] | | 1:500 | |
| **Excitatory synapse** | Vesicular Glutamate Transporter (vGLUT1) | SYSY (135 304) | 1:1000 | 1:2000 | 1:1000 |
| **Inhibitory synapse** | Vesicular GABAergic Transporter (vGAT) Cytoplasmic tail (rb) | SYSY (131 002 and 131 004) | 1:1000 | 1:2000 | 1:1000 |
| **Pan-synaptic marker** | Synapsin | SYSY (106 011 and 106 004) | | | 1:500 |
| **Post synaptic density** | PSD95 | CST (3450), Abcam (ab2723), DSHB (k28/43) | 1:500 | 1:500 | 1:250 |
| **Green fluorescent protein (GFP)** | | Abcam (ab13970) | | 1:1000 | 1:1000 |
| **α-secretase (ADAM10)** | | TFS (MA5-32616) | 1:1000 | 1:500 | 1:500 |
| **β-secretase (BACE1)** | | CST (5606S) | 1:1000 | 1:500 | 1:500 |
| **γ-secretase component Presenilin-1 ( PSEN1)** | Presenilin-1 (D39D1)<br>Presenilin-1 (NTF 3110) | CST (5643)<br>*In house* generated (JW) Eurogentec | 1:1000 | 1:500 | 1:500 |
| **γ-secretase component Nicastrin (NCSTN)** | Nicastrin (D38F9) | CST (5665) | 1:1000 | 1:500 | 1:500 |
| **Histone/Chromatin (Histone H3)** | | Abcam (ab1791) | 1:1000 | | |
| **PIP2** | PtdIns(4,5)P2 (2C11) | Echelon Biosciences (Z-P045) | | | 1:1000 |

| | | | | | |
|---|---|---|---|---|---|
| **Regulating synaptic membrane exocytosis protein 1 (RIM1/2)** | | SYSY (140 217) | 1:1 000 | | 1:500 |
| **Syntaxin-1 (STX1)** | | ProteinTech (83159-6-RR) | 1:1 000 | | 1:500 |
| **Synaptosome Associated Protein 25 (SNAP25)** | | SYSY (111 004) | 1:1 000 | | 1:500 |
| **Synaptotagmin-1 (Syt1)** | C-terminal cytoplasmic domain of Syt1 | SYSY (105 008 and 105 011), DSHB (mAB30- asv30) | 1:1 000 | | 1:500 |
| **Synaptobrevins (Syb)** | | SYSY (104 002) | 1:1 000 | | 1:500 |
| **Vesicle-associated membrane protein-1/2/3 (VAMPs-1/2/3)** | | SYSY (104 102) | 1:1 000 | 1:500 | |
| **Presynaptic cytomatrix protein Bassoon (Bsn)** | | SYSY (141 318) | | | 1:500 |
| **Postsynaptic density-localized scaffolding protein Homer1** | | SYSY (160 004) | | 1:500 | |
| **Puromycin (Puro)** | Recombinant Puromycin (3RH11) | Kerafast (Kf-Ab02366-1.1) | 1:1 000 | | |
| **Flotillin (FLOT1)** | | BDB (610820) | 1:1 000 | | |
| **Transferrin receptor (TFR)** | Transferrin Receptor (YTA 74.4) | IHB (G1-221-12) Abcam (ab22391) | 1:1 000 | 1:200 | |

| | | | | |
|---|---|---|---|---|
| **Glyceraldehyde-3-phosphate dehydrogenase** | GAPDH | SCB (sc-32233) HelloBio (HB9177) | 1:5000 | |
| **Microtubule protein** | | SYSY (188004 and 188 006) | | 1:1000 |
| **Beta-actin** | | SYSY (251 005) | | |
| **Nucleus** | 4′,6-diamidino-2-phenylindole (DAPI) | TFS (D1306) | | |
| **Fluorescent conjugated secondary antibodies** | Goat anti-chicken IgG 405 | Abcam (ab175674) | | 1:500 /1000 |
| | Goat Anti-Chicken IgY H&L A488 | Abcam (ab150169) | | |
| | Goat anti-Mouse IgG A405 | Ozyme (BTM20080-1MG) | | |
| | Goat anti-Mouse IgG A488 | Abcam (ab150113) | | |
| | Goat anti-Mouse IgM (Heavy chain) A568 | Invitrogen (A21043) | | |
| | Goat anti-Mouse IgG A647 | Abcam (ab150115) | | |
| | Goat anti-Mouse IgG Atto647 | Sigma Aldrich (50185-1ML-F) | | |
| | Goat anti-Rabbit IgG A488 | Abcam (ab150077) | | |
| | Goat anti-Rabbit IgG A568 | Abcam (ab175471) | | |
| | Goat anti-Rabbit IgG A568 | Ozyme (BTM20102-1MG) | | |

| | | | | | |
|---|---|---|---|---|---|
| | Goat anti-Guinea Pig IgG A488 | Abcam (ab150185) | | | |
| | Goat anti-Guinea Pig IgG A647 | Abcam (ab150187) | | | |
| | Streptavidin Alexa 568 Conjugate | TFS (S11226) | | 1:1000 | |
| | Streptavidin, Alexa Fluor™ 647 conjugate | Biolegend (405237) | | | |
| **HRP conjugated secondary antibodies** | Goat anti-Mouse IgG (H+L) Secondary Antibody, HRP | TFS (32430) | 1:2500/5000 | 1:1000 | |
| | Goat anti-Rabbit IgG (H+L) Secondary Antibody, HRP | TFS (31460) | | | |
| | Goat anti-Guinea Pig IgG (H+L) Secondary Antibody, HRP | TFS (A18769) | | | |
| | Streptavidin HRP | Biolegend (1474) TFS (N100) | 1:2500 | | 1:1000/2500 |

*WB, western immunoblotting; ICC, immunocytochemistry; IHC, immunohistochemistry; ELISA, enzyme-linked immunosorbent assay; BDB, BD Biosciences; CST, Cell signaling technology; IHB, Iowa Hybridoma bank; SYSY, Synaptic systems; SCB, SantaCruz Biotechnology; TFS, Thermo Fisher Scientific. ‡Gift from Prof. J Walter (University of Bonn, Germany)*

**Table S2. List of antibodies against APP, APP-CTFs and Aβ used in this study.**

| Target of interest | Specificity (Antibody clone) [fig. S10] | Epitope | Species | Remarks |
|---|---|---|---|---|
| **Amyloid precursor protein (APP) and APP C-terminal fragments** | N-terminal specific [AB#3] | N-terminal region of human APP (KLH). | human, rat, mouse | Specific to APP-FL, *no* cross-reactivity to APP-CTFs and Aβ (which lack the N-terminal ectodomain)* |
| | Aβ mid region specific (M 3.2) [AB#2] | residues 10-15 of murine Aβ | rat, mouse | Specific to APP-FL and APP-CTFβ*$ |
| | C-terminal specific (C1/6.1) | conserved carboxyl-terminal 20 residues of APP (residues 676-695 of APP695) | rat, mouse | Specific to APP-CTFs, (*mild* cross-reactivity to APP-FL)*$ |
| | C-terminal specific (3125) ‡ | conserved carboxyl-terminal residues of APP | human (also rat) | cross-reactive between APP-FL and APP-CTFs*$# |
| | C-terminal specific (140-rb) ‡ | conserved carboxyl-terminal residues of APP | human (also rat) | cross-reactive between APP-FL and APP-CTFs*$ |
| **Soluble APP** | sAPPα | soluble fragment resulting from the α-secretase cleavage of APP | | *no* cross-reactivity to sAPPβ β-amyloid or full-length APP*$ |
| | sAPPβ | soluble fragment cleaved N-terminus to the βsecretase (BACE) cleavage site of APP | | *negligible* cross-reactivity to sAPPα or full-length APP*$ |

| | | | | |
|---|---|---|---|---|
| **Amyloid β (Aβ)** | $A\beta_{1-x}$ (82E1) [AB#1] | Asp1 | human (also rat) | Specific to $A\beta_{1-x}$ (cross reactivity to APP- CTFβ)*$ |
| | rAβ | residues 3-16 of rodent Aβ | rat, mouse | specific to murine and rodent, negligible reactivity with human Aβ |
| | $A\beta_{x-40}$ (BAP-29)‡ | Val40 | human (also rat) | Aβ40* |
| | $A\beta_{x-42}$ (BAP-15)‡ | Ala42 | human (also rat) | Aβ42* |
| | hAβ (2964) | residues 1-20 of human Aβ | human specific | hAβ |

Examined by western immunoblotting*, ELISA$. #non-specific band detected at 55kDa. ‡*Gift from Prof. J Walter (University of Bonn, Germany). For details on other antibodies, please visit* [https://www.alzforum.org/antibodies/](https://www.alzforum.org/antibodies/)

### Methods

Immunohistochemistry (IHC) of mouse brain sections Six to eight-week-old wild-type and transgenic vGLUT1$^{GFP}$ mice brains were fixed in 4% (v/v) paraformaldehyde (PFA) and 4% sucrose in 1× phosphate-buffered saline (PBS, pH 7.4) solution. Brains were dissected and sliced to 30 µm. First, a heat-assisted antigen retrieval step (Reveal Decloaker #SKU: RV1000M) was performed to augment and facilitate binding of APP-CT/ APP antibodies for 1 h. Sections were permeabilized (permeabilization buffer; 0.025% Triton X-100 (Merck, #T8787-250ML) in blocking buffer) for 10 min, followed by blocking (2.5% NHS, 2.5% bovine serum albumin (BSA; Carl Roth, #8076-4), 0.0125% Triton X-100 in 1x PBS) for 2 h at RT. Incubation with primary antibodies (respective dilutions in blocking buffer, tab. S1) was performed overnight at 4ºC. Next day, coverslips were washed thrice with PBS and incubated with respective secondary antibodies (respective dilutions in blocking buffer, tab. S1) for 2 h at RT. Removal of the secondary antibody solution was followed by washes of PBS (thrice) and distilled water (once) and sections were incubated with DAPI and mounted under glass coverslips using Moviol mounting reagent (41.67% Glycerol, 16.67% Moviol 4-88 (CarlRoth# 0713.1) in $ddH_2O$).

Microscopy and image processing Confocal microscopy images were acquired on a Nikon SoRa spinning disk super-resolution microscope (Nikon, Japan). Laser power, detector gain, and other parameter settings were kept constant to acquire all comparable samples in the same set. Each immunostaining was performed with a cross-combination of secondary antibodies to rule out non-specific reactivity. Images were acquired using either a 40× or a 100× oil immersion objective, 2048 × 2048 pixels. *z*-stacks were 1x magnification, steps = 16, and step size = 0.3 mm. Imaging was also done with a Leica SP8 Liachroic confocal microscope (Germany) using a 63× oil immersion objective or a Leica widefield fluorescence microscope equipped with Thunder computational clearing module (Germany) using a 20× air objective. Per coverslip, randomly selected 6-10 images were captured, which were further used for quantification, representative of ~30-50 neurons per treatment condition or colocalization analysis. All images depicted herewith were minimally processed using Fiji ImageJ or Imaris software to depict 3D rendering. Mander's overlap coefficient analysis was done using the colocalization processing module of the Fiji ImageJ plugin. Intensity analysis was performed by manually selecting the region of interest (ROI) and measuring the intensity of the individual channels in the selected area. For computing the number of boutons, image masks were created for that particular channel (post thresholding), and selected ROI parameters were analyzed using the analysis module. To normalize the variability in transfection efficiency or the number of neurons in different experimental set-ups, the readings from each experiment were averaged and normalized to the control/untreated neurons from that particular experiment. Average values from each experiment were normalized to the respective controls in the same set. Values from independent experiments were then computed and values are represented as independent data points.

Synaptoneurosome/ synaptosome isolation from cultured neurons/ adult rat brain (fig. S1E, *scheme*, tab. S3) Synaptoneurosomes from cultured neurons were prepared using SYNPER synaptic protein extraction reagent (ThermoFischer Scientific #87793) with smaller modifications. Briefly, treated/untreated neurons were washed with ice-cold PBS and collected in SYNPER reagent supplemented with protease and phosphatase inhibitors. Lysates were homogenized using a 23# needle (21 times) and centrifuged at low speed to remove nuclei and debris (pellet P1; 1200 x g, 5 min). The supernatant (S1) was then subjected to high-speed centrifugation to separate cytosolic material (supernatant, S2) and pellet synaptoneurosomes (P2). The pellet P2 was resuspended in SYNPER reagent and either stored at –20°C for further use or processed immediately. Similar to the protocol for the mice brains, synaptosomes (SYN, tab. S3) from adult rat brains were isolated using the SYNPER protein extraction reagent. To further favor the enrichment of presynaptosome material, ultracentrifugation on sucrose gradients protocol was followed. The collected material between the two sucrose gradients was either stored at –20°C or used further as indicated.

**Table S3. List of synapse preparations used in this study**

| No. | Name (Abbr.) | Notes | Buffer (model) | Scheme reference | Figure reference |
|---|---|---|---|---|---|
| **1.** | Synaptosome (SYN)<br><br>Presorted SYN/ Sorted SYN | synaptosomes constitute resealed presynaptic compartments, sometimes associated with "open" postsynaptic membranes (Whittaker et al. 1964; Luquet et al. 2017; Hafner et al. 2019) | Isotonic buffer (mice brain) | Fig. S2A | Fig. 1, S2 |
| **2.** | Synaptosome fraction (SYN) | (same as 1) | SYNPER reagent (rat brain) | Fig. S2A | Fig. 5, S9B-L |
| **3.** | Synaptoneurosomes (P2 fraction) | synaptoneurosomes constitute resealed pre- and post- compartments (Whittaker et al. 1964)) | SYNPER reagent (dissociated rat neurons) | Fig. S1E | Fig. S1F-G, S2J-K, S5B-D, S9M-O |

ELISA For indirect ELISA, equal protein amounts of different mouse brain fractionated material, synaptosome fraction (SYN), sorted SYN or cellular fractions were then added to the Nunc® MaxiSorp™ 384 well plates (Sigma Aldrich, #P6366) as antigen solutions (0.1-1 µg/ well, 50-75 µl, determined according to the antibody LOD, tab S4) and incubated at 4°C for 16 h. After incubation, residual liquid from the plate was removed by gently tapping the plates and washed with 100 µl of 1x PBS (thrice). 100 µl of blocking buffer (1 mg/ml BSA) was added per well and incubated for 2 h at RT. Wells were further tapped dry and incubated with primary antibodies (respective dilution in blocking buffer- tab. S1, 75 µl). for 2 h at RT or 16 h at 4°C, followed by subsequent washing and incubation with streptavidin-conjugated-HRP complex (1:5000, 50 µl) for another 2 h. Wells were then washed thoroughly four times, tapped dry and filled with 30 µl of 3,3′,5,5′-tetramethylbenzidine substrate (TMB; ThermoFischer Scientific, #34029) in each well and incubated at RT until sufficient blue color developed (time ranged from 2-15 min depending on the different capture antibodies used in the experiment). 30 µl of stop solution (4 M $H_2SO_4$) was added to each well. Absorbance values from each well were read at a Tecan plate reader at a wavelength of 450 nm and a background measurement at 620 nm. Each sample was analyzed in technical duplicate/triplicate wells in each experiment as indicated in the respective graphs. For the sandwich ELISA, plates were precoated with the respective antibody (0.05 – 0.1 mg/well) as capture antibody for 2 h at RT, washed and blocked with 1 mg/ml BSA solution in PBS. Subsequent steps as described above were followed. In certain cases, an ELAST ELISA Amplification System (Revvity, #NEP116001EA) was used as per manufacturer's protocol to augment the signals *via* tyramide signal amplification (indicated in respective figure legends).

**Table S4. LOD of different antibodies examined using in-house ELISA protocol**

| **A.** | **Dilution** | **Protein concentration* (µg)** | **Absorbance (A450)** | | |
|---|---|---|---|---|---|
| **APP-CTFbeta (AB#1)** | | | **Rep#1** | **Rep#2** | **Rep#3** |
| | **1** | 20 | 2,0568 | 1,9887 | 2,1097 |
| | **2** | 10 | 1,9099 | 1,7538 | 1,7733 |
| | **3** | 5 | 1,665 | 1,529 | 1,546 |
| | **4** | 2,5 | 1,2733 | 1,1692 | 1,1822 |

| | | | | | |
|---|---|---|---|---|---|
| | **5** | 1,25 | 0,9305 | 0,8544 | 0,8639 |
| | **6** | 0,63 | 0,5977 | 0,5396 | 0,5456 |
| | ***7*** | ***0,31*** | ***0,2449*** | ***0,2248*** | ***0,2273*** |
| | **8** | 0,16 | 0,1175 | 0,1179 | 0,1091 |
| | **9** | 0,08 | 0,0784 | 0,072 | 0,0728 |
| | **10** | 0,04 | 0,049 | 0,045 | 0,0555 |
| | **11** | 0,02 | 0,0321 | 0,0318 | 0,0336 |
| | **12** | 0,01 | 0,0311 | 0,027 | 0,0273 |
| | **Blank** | 0 | 0,0299 | 0,0118 | 0,0102 |

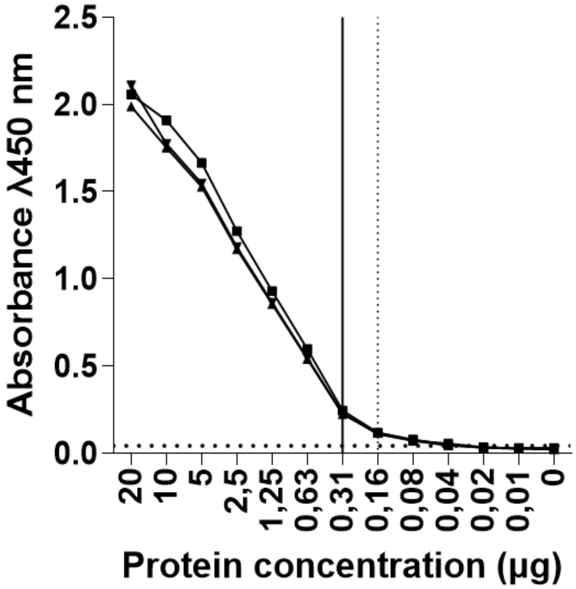

| B. | | Absorbance (A450) | | | |
|---|---|---|---|---|---|
| **Protein** | **LOD concentration (µg)** | **R1** | **R2** | **R3** | **LOD absorbance** |
| **APP-CTF (AB#2)** | 0,31$ | 0,2536 | 0,2723 | 0,2318 | 0,04332 |
| **APP-FL** | 0,63$ | 0,4057 | 0,4356 | 0,3709 | |
| **BACE1** | 0,16$ | 0,1217 | 0,1307 | 0,1113 | |
| **PSEN1** | 0,16$ | 0,1014 | 0,1089 | 0,0927 | |
| **ADAM10** | 1,25$ | 0,9635 | 1,0346 | 0,8808 | |
| **NCSTN1** | 0,31$ | 0,2231 | 0,2396 | 0,2040 | |
| **APP-NT** | 0,32* | 0,1116 | 0,1198 | 0,1020 | |
| **Syb** | 0,63* | 0,4564 | 0,4901 | 0,4172 | |
| **SYT1** | 0,16* | 0,1913 | 0,2280 | 0,2084 | |
| **STX1** | 0,31* | 0,2914 | 0,3089 | 0,3149 | |

*A. Representative values for standard dilution curve for AB#1; B. LOD for different antibodies used in this study. Please note, LOD concentrations were always earmarked to the consecutive higher concentration (dark line) to correct for variability in pipetting errors and to ensure the values would not be below the detection range. Protein samples used were either synaptosome homogenates* or the whole brain homogenates$, prepared as described in respective sections.*

Pharmacological treatments Auxiliary treatments to manipulate APP proteolysis was also performed using GI25423X (1 µM; α-secretase modulator, ASM; Cayman Chemicals #28284-1mg), Compound E (10 nM; γ-secretase inhibition, GSI2; Cayman Chemicals #15579-500ug) and FLI-06 (10 µM; inhibitor of the notch signaling pathway, NSI; Cayman Chemicals #21272-5mg) for 4 h in culture media. Additionally, treatment with CNXQ (10 µM, 30 min; blocking post-synaptic AMPA receptors; Sigma Aldrich #C127-5MG), or AP5 (5 µM, 30 min, post synaptic

NMDA receptor antagonist; HelloBio #HB0225-10mg) or Anisomycin (10 um, 4 h; Tocris #1290) was performed for the specified time durations.

Western blotting / Dot blot analysis Protein content in different material for analysis was estimated using the standard Pierce™ BCA protein assay kit (ThermoFischer Scientific, #23225). For direct comparison between the samples, equal amount of protein was loaded (10-20 µg protein per lane), except Fig. 1B (S1, P2, presort. SYN = 20 µg, Sort. SYN = 1 µg). Samples were dissolved in Laemmli buffer and subjected to boiling at 95°C for 5 min or not for vGLUT1 or vGAT proteins. They were loaded on a precast 4–15% or 4-20% NuPage® Bis-Tris gels (Bio-Rad, #4561083EDU, 4561096EDU). The separated proteins were electro-transferred onto a 0.2-µm nitrocellulose (NT) membrane (Amersham, #1060001) for 10 min using the quick transfer protocol (Trans-Blot turbo transfer system, Bio-Rad). After blotting, for APP/ APP-CT antibodies, membranes were boiled in PBS for 10 min. Ponceau staining was done to examine protein loading of samples. For dot blot, samples (1 mg/ml) were blotted as a single drop on nitrocellulose membranes, allowed to dry and blocked with 2.5 % BSA in TBS-T (TBS [50 mM Tris-HCl, pH 7.5, 150 mM NaCl]. The blots were then incubated in 5% skim milk (Cal Roth, #T145-1) in TBS-T containing 0.1% Tween 20 (ThermoFischer Scientific, #J20605.AP) for 1 h at RT. Blots were incubated overnight at 4°C in the primary antibody solution (respective dilutions in TBS-T, tab. S1). The next day, blots were washed with TBS-T thrice (5 min each) and incubated with respective dilutions secondary antibody solutions in blocking buffer (1:10000, tab. S1) for 2 h at RT. Proteins were detected either by ECL imager (Bio-Rad, Germany) using ECL solutions; equivalent solution A (0.1 M Tris, 0.4 mM coumaric acid, 2.5 mM luminol, pH 8.5) and solution B (0.1 M Tris, 0.018% $H_2O_2$) mixed prior to application on blots, or Li-COR using secondary antibody conjugated to IR 680/800 dyes as reporter to the primary antibody. WB detection reagents were from GE or LiCOR Biosciences. Quantification of band signals was done using ImageJ – Gel processing module or Image Studio processing software (LiCOR Biosciences, Germany). All samples were analyzed in biological duplicates in two or three independent experiments. Values from each sample were normalized to the values from the control cells in that experiment and fold-change with respect to control was indicated as graphs.

Puromycylation assay / Metabolic labelling of nascent proteins To assess active protein synthesis in untreated or pretreated cultured neurons, we used the reported puromycylation assay (*1*). Briefly, cells were incubated with puromycin (5 µM) or FAM-puromycin (0.1 µg/ml) diluted in warm culture medium for 5 min at 37°C under standard culture conditions. Additionally, to confirm puromycin incorporation reflects *de novo* protein synthesis, we also used anisomycin (40 µM, 30 min) before and during the puromycin treatment. Following labelling, cells were immediately placed on ice, washed with ice-cold PBS (thrice) to halt metabolic activity and get rid of extra puromycin, and lysed in STEN lysis buffer supplemented with protease and phosphatase inhibitors for dot blotting. For microscopy, cells were washed with ice-cold PBS (thrice) and fixed with 4% PFA and processed for immunostaining. All experimental conditions were performed in parallel and repeated in at least three independent biological replicates.

Quantification of puromycylation signal intensity was normalized to total protein levels or appropriate housekeeping controls as indicated.

Quantification of SynaptoRed and Synapsin1 positive synapse numbers For SynaptoRed (SR) staining, live neurons (untreated/ pretreated) were loaded with SR with an electrical stimulation to label active presynaptic terminals, as imaged as described earlier. Quantification was performed using ImageJ (Fiji) software for images obtained in the initial 6 s (no stimulation). Subsequently, untreated or treated neurons were processed for ICC using primary antibodies against Synapsin 1 (Syn1), protocol described earlier. For both Syn1 and SR datasets, raw images were background-subtracted, and synaptic puncta were detected using consistent thresholding and particle analysis settings for manually selected ROIs selected 5-10 µm away from the soma of the neurons. Synaptic puncta were defined as discrete, non-nuclear, punctate structures with sizes between 0.2–2.0 $\mu m^2$. Synapse numbers were expressed as average puncta per 10-20 µm dendrite length or per field of view, depending on the experiment. A minimum of 8–15 fields per condition from at least 3 independent cultures were analyzed, and all quantification was performed in a blinded manner.

Preparation of monomeric Aβ peptides Synthetic Aβ peptides (1 mg) were obtained as lyophilized powders and stored at −20°C until use. To prepare monomeric Aβ, lyophilized peptides were first dissolved in 10 mM NaOH (in $ddH_2O$, 0.2 µm prefiltered) to a final concentration of approximately 230 µM. The solution was sonicated in a water bath for 10 min to aid dissolution and disaggregation. Following sonication, the peptide solution was quickly aliquoted in prelabelled low-bind tubes, flash frozen in liquid nitrogen, and stored at −80°C until further use. Prior to each experiment, master aliquots were thawed and further aliquoted to smaller volumes necessary for individual experiment, minimizing freeze–thaw cycles to avoid aggregation. These sister aliquots were further diluted to the desired working concentration in culture medium, just before application to cells. Excess solution was discarded.

Analysis of membrane-bound and extracellular Aβ species To examine membrane-associated and extracellular Aβ species, untreated or γ-secretase inhibitor (GSI) pretreated neurons were treated with or without synthetic Aβ peptides (100 nM) for 5 min at 37°C. Following incubation, cells were washed three times with ice-cold PBS for biochemical assays. For WB analysis, neurons were directly lysed in an ice-cold STEN lysis buffer (supplemented with protease and phosphatase inhibitors). For ICC, neurons and post washes were fixed in 4% PFA. To assess extracellular Aβ levels, conditioned media, collected immediately after treatment (supplemented with 2% sodium deoxycholate, Carl Roth #3484.3), subjected to trichloroacetic acid (TCA, Thermo Fisher Scientific #A11156.30) precipitation (conc. made up to 35%, incubated for 1 h), and the resulting protein pellet (16,100 x g, 10 min) was subjected to ice-cold acetone washes (twice) and resuspended in a Laemmli sample buffer. All samples were processed in parallel to ensure consistency across treatment conditions.

APP knockdown Cultured neurons were transfected with siRNA targeting APP (ThermoFischer Scientific, #4390815; siRNA ID s132407, standard, 20 nmol, species: *Rattus norvegicus*) using Lipofectamine 2000 (ThermoFischer Scientific, #11668027) according to the manufacturer's protocol with minor adaptations. Briefly, siRNA-Lipofectamine complexes were prepared in Opti-MEM (ThermoFischer Scientific, #31985062) and incubated for 20 min at RT. The siRNA-Lipofectamine complexes were then added to the neurons in transfection medium (BrainPhys™ neuronal medium supplemented with NeuroCult™ SM1 neuronal supplement without antibiotics), and incubated for overnight at 37°C. Next day, neurons were washed thrice with transfection medium replaced with the regular neuronal culture medium. Neurons were then allowed to recover for 7 days prior to downstream analysis. Transfection efficiency, assessed *via* ICC, ranged between 70–85% across all analyzed coverslips.

Live antibody labelling experiments Live antibody labelling was performed on cultured neurons to detect surface-exposed epitopes under physiological conditions. Briefly, neurons were gently washed once with pre-warmed culture medium to remove debris. Primary antibodies targeting extracellular epitopes (AB#1-3, table. S1) were diluted in 1:1 conditioned and fresh pre-warmed neuronal culture medium, added directly to the cells for 10–15 min at 37°C in a humidified incubator. Following incubation, cells were carefully washed three times with warm culture medium to remove unbound antibodies and processed for live imaging experiments as discussed earlier. For visualization of surface labeling *via* ICC, neurons were either fixed immediately with 4% PFA and processed for subsequent permeabilization and intracellular staining, as described. Samples for EM experiments were processed as described below.

Electron microscopy Neurons cultured on glass-bottom dishes were live labelled with or without antibody against the extracellular N-terminal domain of APP-CTFβ, N-terminal ectodomain of full-length APP in culture media for 15 min at 37 °C. Neurons were washed with fresh media thrice and fixed with EM grade 4% PFA (Electron Microscopy Sciences #157-4) in 0.2 M HEPES (Sigma Aldrich, #H4034-100G) buffer on ice. Post fixation, neurons were washed thrice in HEPES buffer, blocked with 2% NGS (normal goat serum) for 30 min at RT, followed by incubation with Ultrasmall-gold secondary antibody (1:50; Aurion, #800.022/11) in blocking buffer. Excess antibody was washed away with HEPES buffer thrice, followed by second fixation step [2% glutaraldehyde (Carl Roth #4157.1) in 0.2M HEPES pH 7.2] on ice for 1 h. Neurons were washed and quenched with $NaBH_4$ (Sigma Aldrich, #213462-25G; 1mg/ml in 0.2M HEPES) for 15 min at RT. Once the bubbling stopped, secondary quenching with 200mM glycine (Merck, #GE17-1323-01) and 200mM ammonium chloride (Sigma Aldrich, #A4514-100G) in 0.2M HEPES for 15 min RT. After washes with HEPES, additional 3x washing with ECS (Enhancement Conditioning Solution, Aurion, #500.055) was performed (5 min each). Silver amplification was performed using protocol indicated in the manufacturer's protocol (Aurion, #500.044). Post-amplification after 90 min, neurons were washed with ECS solution (3x, 5 min each) and $ddH_2O$ (3x, 5 min each), followed by osmification $OsO_4$ (Info, 0.2% in $ddH_2O$) for 30 min at RT. Osmified neurons were washed and stained with uranyl acetate (Info, 0.25% in $ddH_2O$, in dark for

1 h at RT), following 5x washes with ddH2O.Neurons were then dehydrated in a series of increasing ethanol solutions (twice 30, 50, 70, 90, twice 95, and twice 100% ethanol 10 min each) at RT. Cells were then infiltrated with Epon epoxy resin (Sigma Aldrich, #45359-1EA-F) in increasing ratios of ethanol: Epon (1:1 for 1 h, 1:2 overnight at RT), and finally twice with 100% Epon for 1 h at RT. Epon was polymerized by curing at 60°C for 48 h. After polymerization, samples were unmounted from the glass-bottom dishes using freeze-thaw cycles of liquid nitrogen and a heating plate set to 60°C. The retrieved block face was trimmed to fit on an EM grid (formvar and copper-coated copper slot grid, Science Services, #EFCF2010-Cu-50) and ultra-thin sections 50-nm-thick were collected. Sections were counterstained with 1% aqueous uranyl acetate for 25 min and lead citrate Ultrastain solution (Leica, #16707235) for 7 min with thorough washing and drying in between. Sections were imaged with a Zeiss Crossbeam 550 (acceleration voltage: 30 kV, probe current: 150 pA, high resolution mode) using a STEM detector. For all examined samples, multiple regions were imaged (>10) with resolution between 1.8 - 8 nm per pixel, across each EM grid. This ensured classification and avoided inadvertent production of a biased/subjective data selection. Best representative images have been illustrated in fig. S6A, and additional images have been depicted in S10A, all images were depicted as is or minimally processed to adjust brightness/contrast using Fiji ImageJ software.

Computational modelling and simulations

*Initial Structure and coarse-graining* The protein structure was obtained from an earlier study by Pantelopulos *et al. (2)* that was made available through the author's GitHub (*3*). A visual inspection of the structure revealed that the side chain of arginine 76 protrudes through the ring of proline 73, a situation that does not occur naturally. To remedy the situation, we manually rotated the side chain of arginine76 by –90 degrees around the axis provided by its C$a$ and C$\beta$ atom, i.e. the axis linking the side chain to the backbone. The side chain had to be rotated clockwise to avoid collision with the rest of the protein structure, most severely histidine77. The generous amount of the rotation was then checked throughout a first simulation, which was eventually used as the first production simulation. In the simulation, the arginine's side chain freely rotates about coming as close as 3 Å to the proline. The dihedral angle spanning its *N* C*A* C*G* C*Z* atom to encapsulate the rotation of the whole side chain spans almost the whole spectrum (Please see, fig. S11).

*All-Atom* The all-atom simulation was set up using CHARMM force field and was set up using CHARMM-GUI (*4–14*). A single copy of the protein was placed in a 16x16 nm² membrane. The different membrane configurations were created with different lipid compositions as shown in the table. S3. The membranes are denoted as 'Realistic' and 'POPC'. Only the realistic membrane is asymmetric, with differences in the lipid compositions of the inner and outer monolayers. Following the CHARMM-GUI workflow, the membrane protein complex was submerged in TIP3P water and solvated to a concentration of 0.15 M NaCl.

**Table S5:** Composition of the different membrane types in %. Only the realistic membrane differs in the composition of its outer and inner membrane leaflet. 3 Lipids and POPC are symmetric membranes.

| | **Realistic presynaptic** | | **POPC** |
|---|---|---|---|
| | *Inner leaflet* | *Outer leaflet* | |
| **POPC** | 30 | 45 | 100 |
| **CER160** | 0 | 10 | |
| **CHL1** | 25 | 25 | |
| **POPE** | 25 | 7 | |
| **CER3** | 2 | 2 | |
| **POPS** | 10 | 0 | |
| **PSM** | 3 | 11 | |
| **POPI** | 3 | 0 | |
| **PI(4,5)P2 (SAPI24)** | 2 | 0 | |

*Coarse-Grained* The refined protein structure (see above) was used to coarse-grain using Martini 3 model (*15*) with the vermouth and martinize2 framework (*16*). The first protein structure used the elastic network approach (*17*) with an elastic force constant of 700 kJ/mol/nm². The second protein structure employs the Go model (*18*). The contact map was calculated using martinize2 with a structure bias of 9.414 kJ/mol. The initial structures were then placed in a membrane system. Starting configurations for both protein models with one protein in a 20x20 nm² membrane and two and three proteins in a 30x30 nm² membrane were created using TS2CG 2.0. The membrane compositions followed the analog scheme to the all-atom case with lipids obtained from the Martini3 lipidome (*19*).The systems were solvated in Martini3 water beads and NaCl salt concentration of 0.15 M was introduced to the system using the solvation tool in TS2CG.

**Table S6:** Lipid composition for coarse-grained membranes showing the corresponding all-atom lipid names.

| **AA** | **CG** | **Realistic presynaptic** | **POPC** |
|---|---|---|---|

| | | *Inner leaflet* | *Outer leaflet* | |
|---|---|---|---|---|
| POPC | **POPC** | 30 | 45 | 100 |
| CER160 | **XCER** | 0 | 10 | |
| CHL1 | **CHOL** | 25 | 25 | |
| POPE | **POPE** | 25 | 7 | |
| CER3 | **PCER** | 2 | 2 | |
| POPS | **POPS** | 10 | 0 | |
| PSM | **SSM** | 3 | 11 | |
| POPI | **POPI** | 3 | 0 | |
| PI(4,5)P2 (SAPI24) | **POP6** | 2 | 0 | |

*Simulations* The initial setup was minimized and equilibrated in multiple steps for the all-atom and coarse-grained simulations. Whenever applicable, the temperature was set to 310 K and the pressure to 1 bar.

All-atom: The equilibration was performed in seven steps with gradually reducing the strength of restraints on the backbone and side-chain atoms. The time step was gradually increased from 1 fs to 2 fs. After two equilibration steps the ensemble was switched from NVT (number of particles, volume, and temperature constant) to NPT (number of particles, pressure, and temperature constant). The chosen barostat was C-rescale and the thermostat was V-rescale. The production simulation was 500 ns for each replica.

Coarse-grained: Depending on the system a different number of equilibration steps were employed, similarly to the all-atom, restraints were gradually lifted. As before, during equilibration the ensemble was switched from NVT to NPT and the timestep was increased from 10 fs to 20 fs. In NPT equilibrations, the Berendsen barostat was used; for the production simulation Parrinello-Rahman was used. The thermostat was v-rescale. The final production simulations were conducted for 5 μs with a timestep of 20 fs.

Analysis Methods

*Fourier Fitting* Throughout the fitting, MD analysis was used to extract positions from the GROMACS (version 2023.3) simulation trajectory. The upper and lower monolayers were separated into two index groups based on the position of the PO4 or P beads/atoms. The two resulting point clouds were then fitted to a grid. The fitting was calculated using the Fourier least-squares fitting [CITE SHIGA]. This allows the generation of a design matrix, in which each row corresponds to a (x,y) tupel, while each column contains the corresponding Fourier terms.

$$z(x,y) = \sum_{n=-N_x}^{N_x} \sum_{m=-N_y}^{N_y} A_{nm}\left(\cos[p_n x + q_m y] + \sin[p_n x + q_m y]\right)$$

The Fourier sum contains three harmonic oscillators in each directions, which corresponds to the wavenumber cutoff off 2 $nm$^{−1}.

Now the system of equations can be solved through the least square solver to obtain the Fourier coefficients $A_{nm}$:
$A \cdot coeffs \ = \ b$,

with A being the design matrix, and b describing the z-values from the input data. The least square solver then minimizes the difference between solutions obtained using the coefficients and the real z-values. Thus, the vector coefficients then contain the Fourier coefficients $A_{nm}$ that can be mapped back onto the x,y grid. From these coefficients the middle of the membrane is constructed as the pairwise average of the coefficients for each monolayer, which provide a function for each monolayer.

*Thickness and Radial Thickness* The thickness is given for each grid point as the difference between the fitted surface of the upper and lower monolayer. The average thickness over the considered time frame is then plotted. The same average thickness then used to derive the radial thickness. In sequential steps, the radius is increased and traversed. In specified intervals the thickness is interpolated (the polar coordinates considered do not necessarily match the grid). The thickness is then averaged over each circle and plotted. The radius of the circles is increased by increments of .05 nm until the circle exceeds the simulation box size. The number of angles considered is influenced by the radius, such that a bigger circle is traversed with more angles. Angles are equally spaced from 0 until, but not including 360 degrees. The number of angles considered is $max(10 \cdot 2 \cdot \ \pi \cdot \ radius, 2)$.

Membrane thickness analysis - The thickness of each membrane was measured and plotted. Each box has the width and height of the simulation box. A grid is placed over the box, and the thickness is the difference between the Fourier approximations interpolating the grid. The thicknesses are also shown for membranes without any proteins. If proteins are present in the simulation, a representative protein location is shown through white markers indicating the position of carbon alpha (Cα) atoms - CA atoms (all-atom) or backbone beads (coarse-grained).

Radial thickness -_Considering the average thickness over a certain simulation time as shown in the thickness plots above allows us to judge the influence of the proteins on the membrane thickness. Note that this analysis is only performed if a single protein cluster emerged in the simulation. Single protein simulations by default thus form a single cluster. Beginning at the center, the average thickness on a circle of a specified radius is calculated and plotted.

Height Residue C-Asn99 (GO model) - The distance of the C-terminal for clustered systems for the Go Martini Model is shown. The heights for all-atom and elastic network model are provided in the main manuscript.

*Interaction energy analyses* The interaction energies are calculated analogously for the interaction bar charts and the interaction energy network demonstrated in Fig. 5 of the main manuscript. In each case the energy groups are defined and a rerun in GROMACS is initiated. For the bar charts, the energy groups consist of each present Arg76 residue individually and each lipid type forming a group. The all-atom energy interaction matrix considers each lipid and each amino acid residue as their own energy group. As the number of allowed energy groups per rerun in GROMACS is limited to 63, the system was split up into blocks of 30 residues. The groups were then iteratively rerun to achieve all possible combinations without considering the order. The analysis of non-bonded interaction energies from the Arg76 residue to the different lipid types was only performed for the realistic membrane (Fig. S14-15). The energy contributions are divided into Lennard-Jones and Coulomb interactions, indicated in different colors as mentioned, respectively. For systems containing multiple proteins, thus multiple Arg76, the data is shown for each protein individually (Fig. S15).

**Figure S1.**

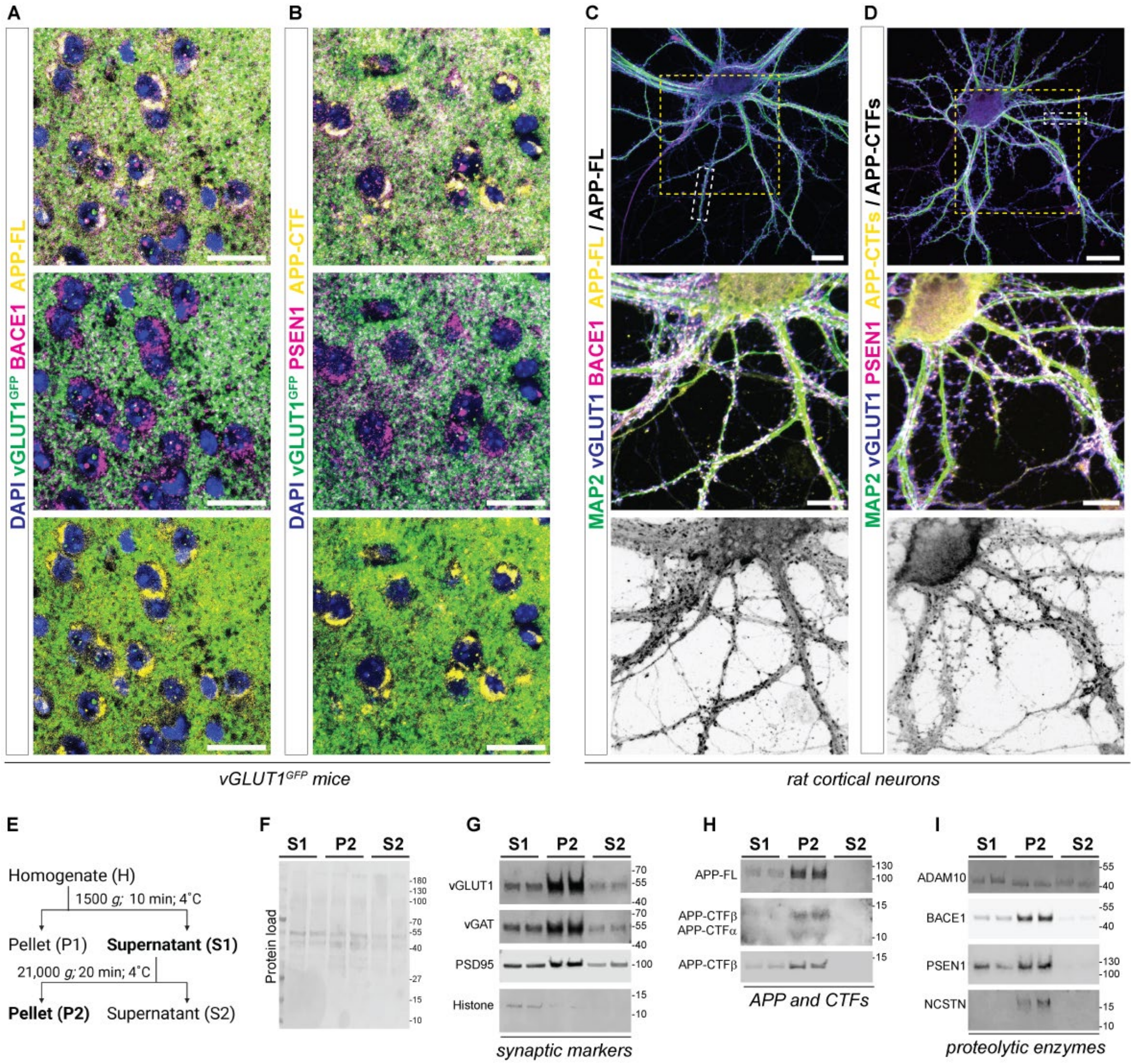


**Figure S1. Localization and distribution of APP, APP-CTFs, and processing enzymes.** (**A-B**) Immunohistochemical analyses of vGLUT1$^{GFP}$ transgenic mice forebrain sections (m, 6 w) showing localization of APP-FL (*yellow,* **A**) and BACE1 (*magenta,* **A**) along with APP-CTFs (*yellow,* **B**) and PSEN1 (*magenta,* **B**) alongside GFP positive vGLUT1 synapses (*green*) signals in the cortex. Sections were co-stained for DAPI to visualize the nucleus (*blue*). Scale bar = 20 µm, N = 2 animals/experiments. (**C-D**) Immunocytochemical analyses of rat cortical neurons (DIV21) showing localization of APP-FL (*grayscale, yellow-merge,* C) and BACE1 (*magenta,* **C**) along with APP-CTFs (*grayscale, yellow-merge,* D) and PSEN1 (*magenta*, **D**). Neurons were co-stained for vGLUT1 (excitatory synapse, *blue*) and MAP2 (dendrites, *green*), respectively. White dotted box depicts the ROI depicted in Fig. 1O-P. Yellow dotted box depicts the enlarged ROI depicted in the panels below to show the localization of APP and APP-CTFs also within the somatic-dendritic compartments. Scale bar = 20 µm, N = 3. (**E**) Schematic representation depicting synaptosome enrichment from rat critical neurons. Pellet P1 contains mostly nuclear material. The supernatant (S1 fraction) upon further centrifugation results in the enrichment of synaptoneurosomes in pellet P2 (synaptoneurosomes containing both pre- and post-synaptic components, tab. S3) wherein the supernatant S2 is synaptosome depleted material. (**F-I**) Western immunoblots depicting the relative levels of protein load (Ponceau staining, **F**) in S1, P2 and S2 fraction. and probed with antibodies against vGLUT1, vGAT, PSD95, Histone H3 (**G**); APP-FL, APP-CTFs, APP-CTFβ (**H**); and its processing enzymes ADAM10, BACE1, PSEN1 and NCSTN (**I**). Samples presented here represent biological duplicates, N = 3.

**Figure S2.**

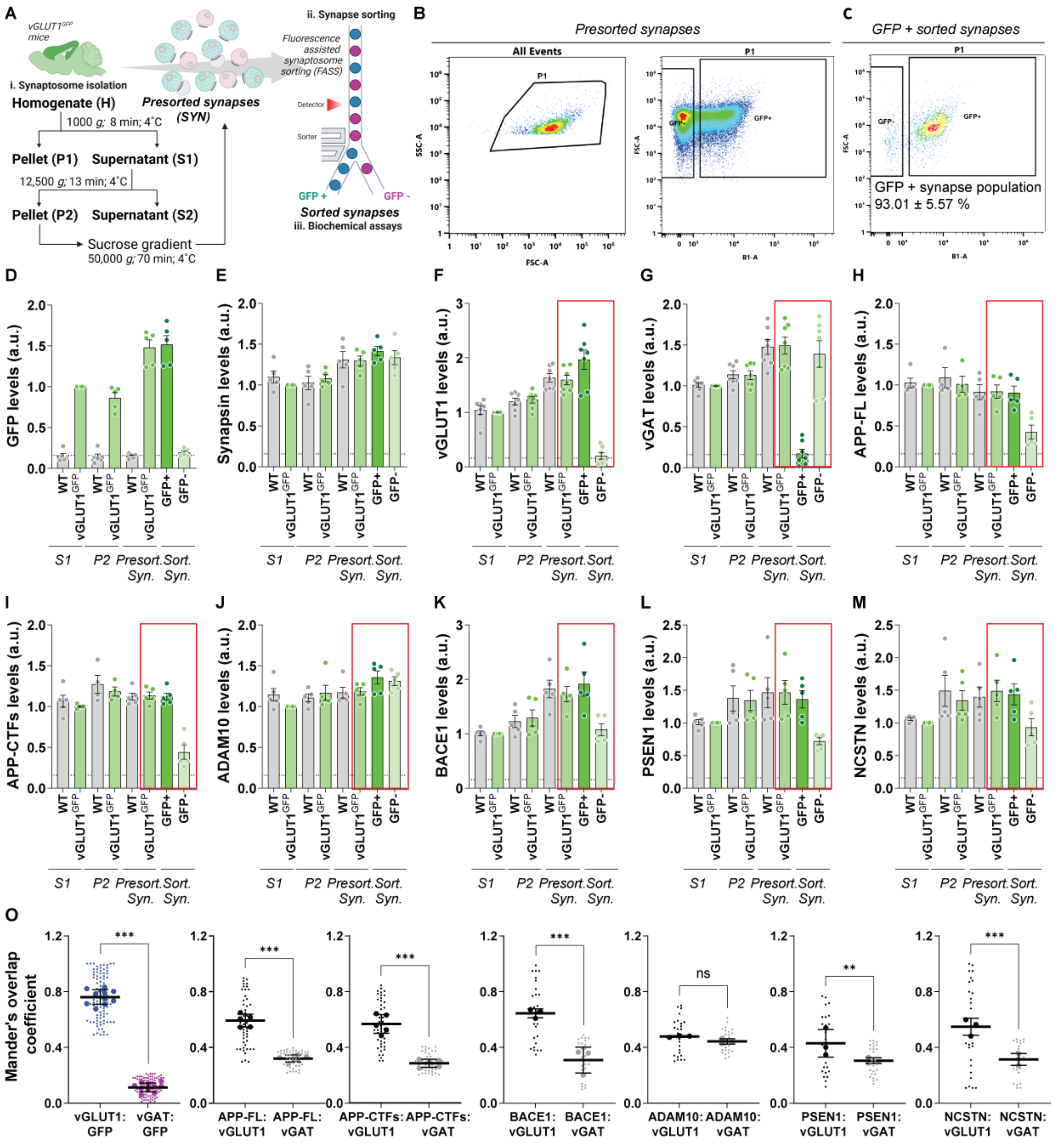


**Figure S2. FASS and biochemical analyses of GFP+ and GFP– synapses from vGLUT1$^{GFP}$ mice.** (**A**) Schematic representation depicting synaptosome isolation (i) and FAS sorting (ii) of GFP+ (GFP positive) and GFP – (GFP negative) synapses from vGLUT1$^{GFP}$ mice forebrain (m, 1y) for further biochemical analysis (iii) using ICC and ELISA. (**B-C**) Sorting of synapses (singlets, all events and gating of GFP+ and GFP– synapse populations). Sorted GFP+ synapses were rerun to determine purity of sorted population (C). Values indicate the mean and standard deviation of 5 experimental replicates, N = 5 animals/ experiments. (**D-M**) Indirect ELISA analyses depicting levels of GFP (D), synapsin (E), vGLUT1 (F), vGAT (G), APP-FL (H), APP-CTFs (I), ADAM10 (J), BACE1 (K), PSEN1 (L) and NCSTN (M) in all fractions starting from S1, P2, unsorted synapses and sorted synapses. Values were normalized to S1 fraction of $vGLUT1^{GFP}$ mice. Dotted line indicates average values from blank wells. As seen in the graphs, GFP signals are specific to the transgenic mice and the sorted population of the synapses. Red box highlights data depicted in Fig. 1 D, E, G, H, J, K, M, N, respectively. n = 10-15, N = 5. (**O**) Mander's overlap coefficient of overlap determined from ICC analysis of sorted synapses. vGLUT1/vGAT: GFP n = 20, N = 5; APP-FL/APP-CTFs: vGLUT1/vGAT, n = 10, N = 5; ADAM10/BACE1/PSEN1/NCSTN1: vGLUT1/vGAT, n = 6, N = 3. ns ($p > 0.05$), * ($p \leq 0.05$), ** ($p \leq 0.01$) , *** ($p \leq 0.01$); One-way ANOVA.

**Figure S3.**

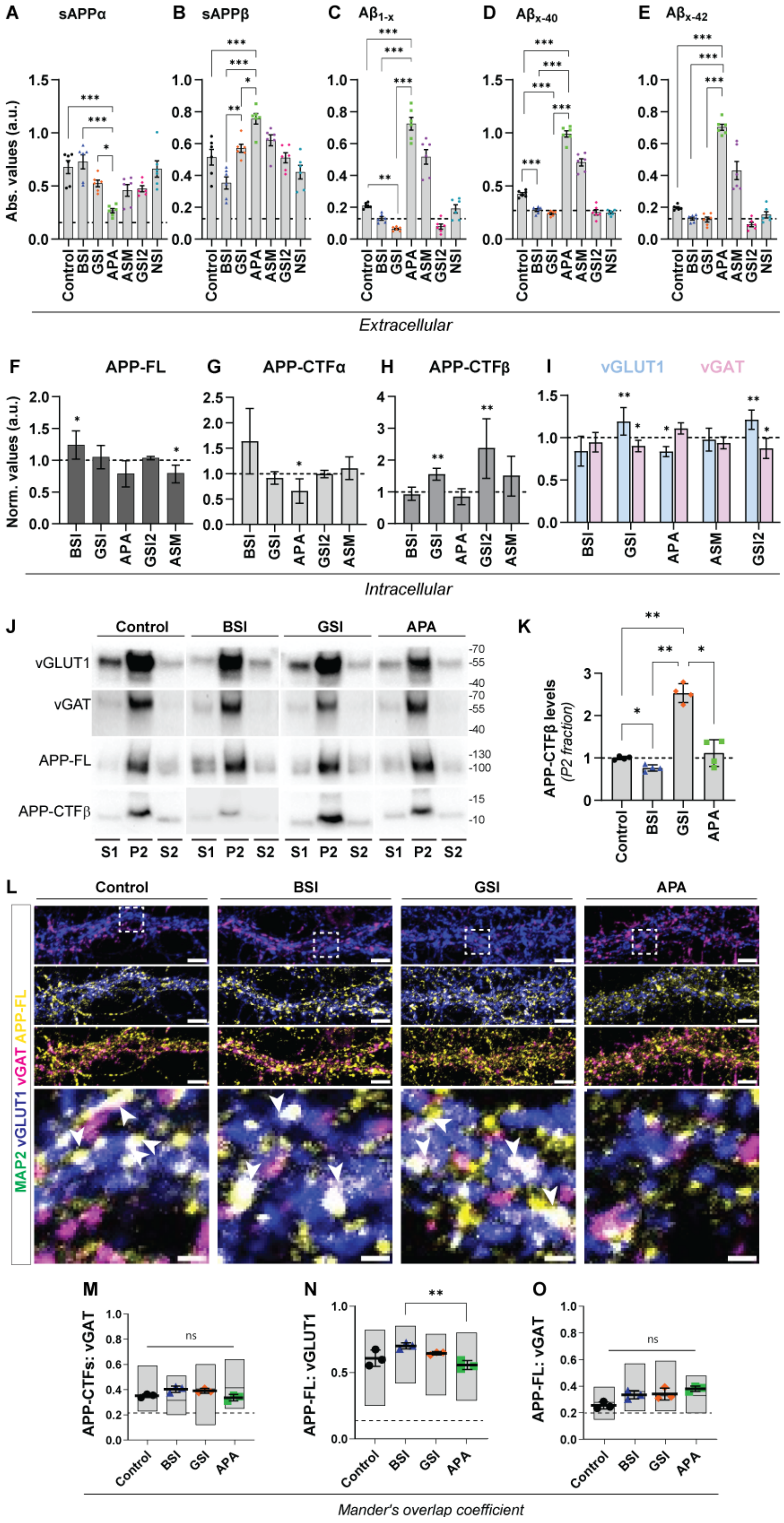
A sAPPα
B sAPPβ
C Aβ1-x
D Aβx-40
E Aβx-42
Abs. values (a.u.)
Control BSI GSI APA ASM GSI2 NSI
Extracellular
F APP-FL
G APP-CTFα
H APP-CTFβ
I vGLUT1 vGAT
Norm. values (a.u.)
BSI GSI APA GSI2 ASM
Intracellular
J
Control BSI GSI APA
vGLUT1
vGAT
APP-FL
APP-CTFβ
S1 P2 S2
K
APP-CTFβ levels (P2 fraction)
Control BSI GSI APA
L
Control BSI GSI APA
MAP2 vGLUT1 vGAT APP-FL
M
APP-CTFs: vGAT
N
APP-FL: vGLUT1
O
APP-FL: vGAT
ns
Mander's overlap coefficient

**Figure S3. Pharmacological modulation of APP processing alters distribution of synaptic protein in cultured rat neurons.** (**A-I**) Rat cortical neurons (DIV21) were treated with LY2886721 (1 µM; BACE1 inhibition, BSI), DAPT (10 µM; γ-secretase inhibition, GSI) or Aftin-4 (5 µM; activator of APP processing, APA) for 4 h, to pharmacologically manipulate different stages of APP proteolysis. Additionally, we tested GI25423X (1 µM; α-secretase modulator, ASM), Compound E (10 nM; γ-secretase inhibition, GSI2) and FLI-06 (10 µM; inhibitor of the notch signaling pathway, NSI) for 4 h. Post-treatment media was used to detect levels of sAPPα (**A**), sAPPβ (**B**), Aβ (**C**), $A\beta_{x\text{-}40}$ (**D**) and $A\beta_{x\text{-}42}$ (**E**) using indirect ELISA. Neuron homogenate was subjected to western blotting to examine changes in the level of APP-FL (**F**), APP-CTFα (**G**) and APP-CTFβ (**H**) along with vGLUT1 and vGAT (**I**), respectively. n = 6, N = 3. (**J-K**) Neurons post-treatment was subjected to synaptosome enrichment, loaded on western blots to examine changes in levels of proteins between the fractions. While APP-FL remains relatively comparable in the P2 fraction, differences in levels of APP-CTFβ are significant. Dotted line represents the average levels of APP-CTFβ within P2 (synaptic) fraction in untreated conditions. N = 2. (**L**) ICC depicting relative changes in the levels of full-length APP (APP-FL) upon treatment with different compounds (*gray, yellow-merge*). Neurons were also co-stained with vGLUT1 (excitatory synapses, *blue*), vGAT (inhibitory synapses, *magenta*) and MAP2 (dendrites, *green*). Scale bar = 5 µm. Box selection depicts the zoomed in panel. Arrows depict colocalization/overlap of APP-FL with vGLUT1+ synapse. Scale bar = 1 µm, N = 4. (**M-O**) Scatter plot depict Mander's overlap coefficient of colocalization between APP-FL: vGLUT1 (**N**). MOC values between APP-FL (**O**) and APP-CTFs (**M**, images depicted in Fig. 2F) with vGAT positive boutons show no significant changes among treatment groups. N = 4. ns ($p > 0.05$), * ($p \leq 0.05$), ** ($p \leq 0.01$) , *** ($p \leq 0.01$); One-way ANOVA.

**Figure S4.**

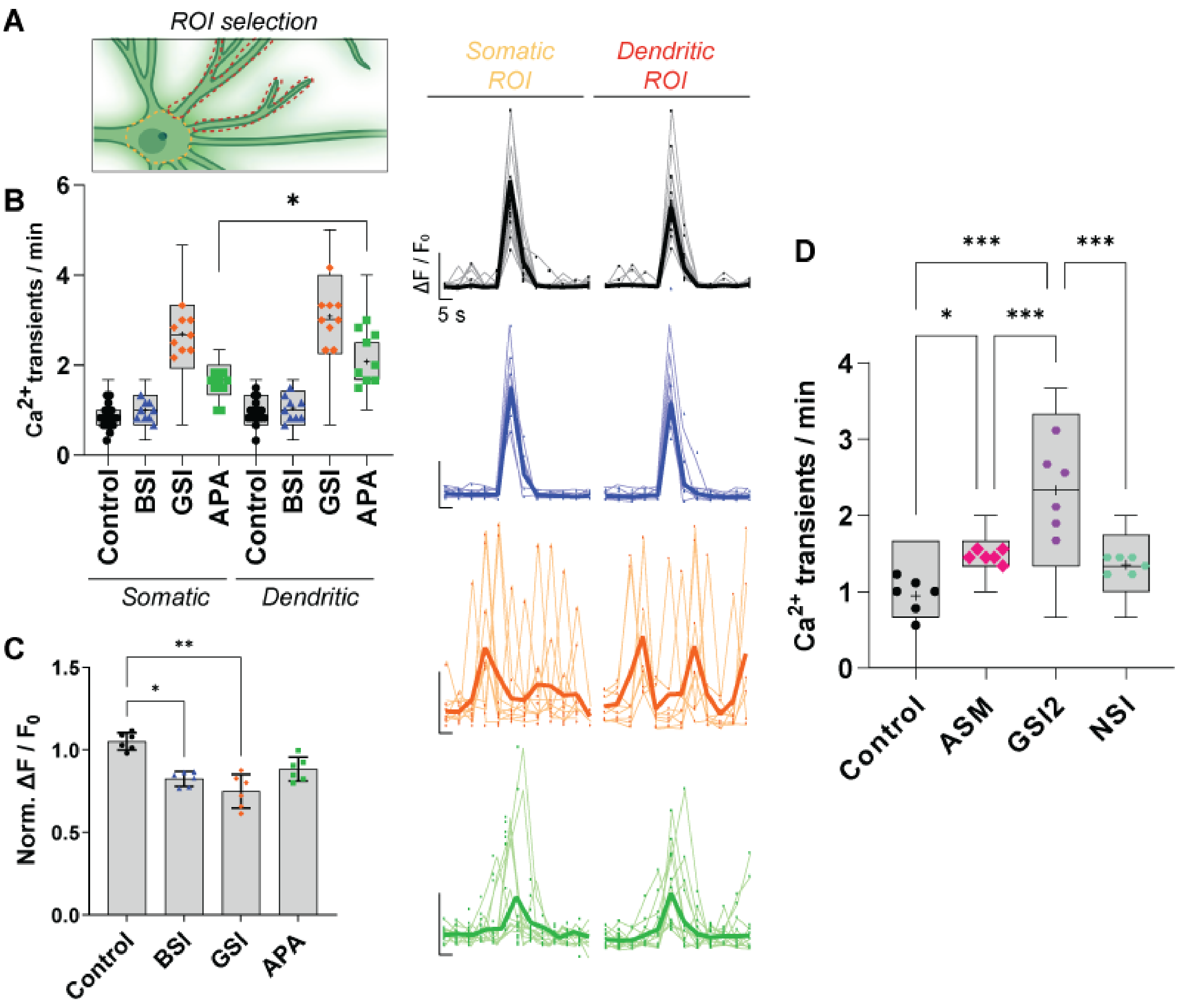


**Figure S4. Spontaneous $Ca^{2+}$ transients in neurons after pharmacological modulation of APP proteolysis.** (**A**) Spontaneous calcium transients were recorded in the absence and presence of $Ca^{2+}$ in Tyrode buffer from Camk2-GcAMP6 expressing neurons post pharmacological manipulation of APP proteolysis (data depicted in Fig. 2I-J represent total neuronal $Ca^{2+}$ changes). ROIs were further segmented to differentiate between soma and dendrite (*schematic representation*). Somatic and dendritic $Ca^{2+}$ traces ($\Delta F/F_0$) per minute have been depicted respectively as (*individual traces- thin lines; average- thick lines*). (**B**) Box plots depict average number of spontaneously evoked somatic or dendritic $Ca^{2+}$ transients per minute, following pharmacological modulation of APP proteolysis. APA treatment shows a possible decoupling between somatic and dendritic $Ca^{2+}$ changes. The differences in values of soma and dendrite in GSI treatment group were not statistically significant. Each data point indicates an experimental replicate. Control n = 18; BSI, GSI, APA, n = 9; N = 3. (**C**) Bar plots depict the $\Delta F/F_0$ normalized to baseline, spontaneously evoked $Ca^{2+}$ transients, following pharmacological modulation of APP proteolysis. Each data point indicates an experimental replicate. N = 3. (**D**) Box plot depicts average number of spontaneously evoked somatic or dendritic $Ca^{2+}$ transients per minute, following auxiliary treatments to manipulate APP proteolysis - GI25423X (1 µM; α-secretase modulator, ASM), Compound E (10 nM; γ-secretase inhibition, GSI2) and FLI-06 (10 µM; inhibitor of the notch signaling pathway, NSI) for 4 h. Modulation of α-secretase shows an increase *wrt* control, comparable to APA treatment conditions (Fig. 2I-J). GSI2 shows a significant increase in number of $Ca^{2+}$ transients when compared to control (comparable effect as GSI, depicted in Fig. 2I-J). GSI-induced effect on APP-CTF accumulation is independent of the Notch signaling pathway. Each data point indicates an experimental replicate, n = 6; N = 3. ns ($p > 0.05$), * ($p \leq 0.05$), ** ($p \leq 0.01$) , *** ($p \leq 0.01$); One-way ANOVA.

**Figure S5.**

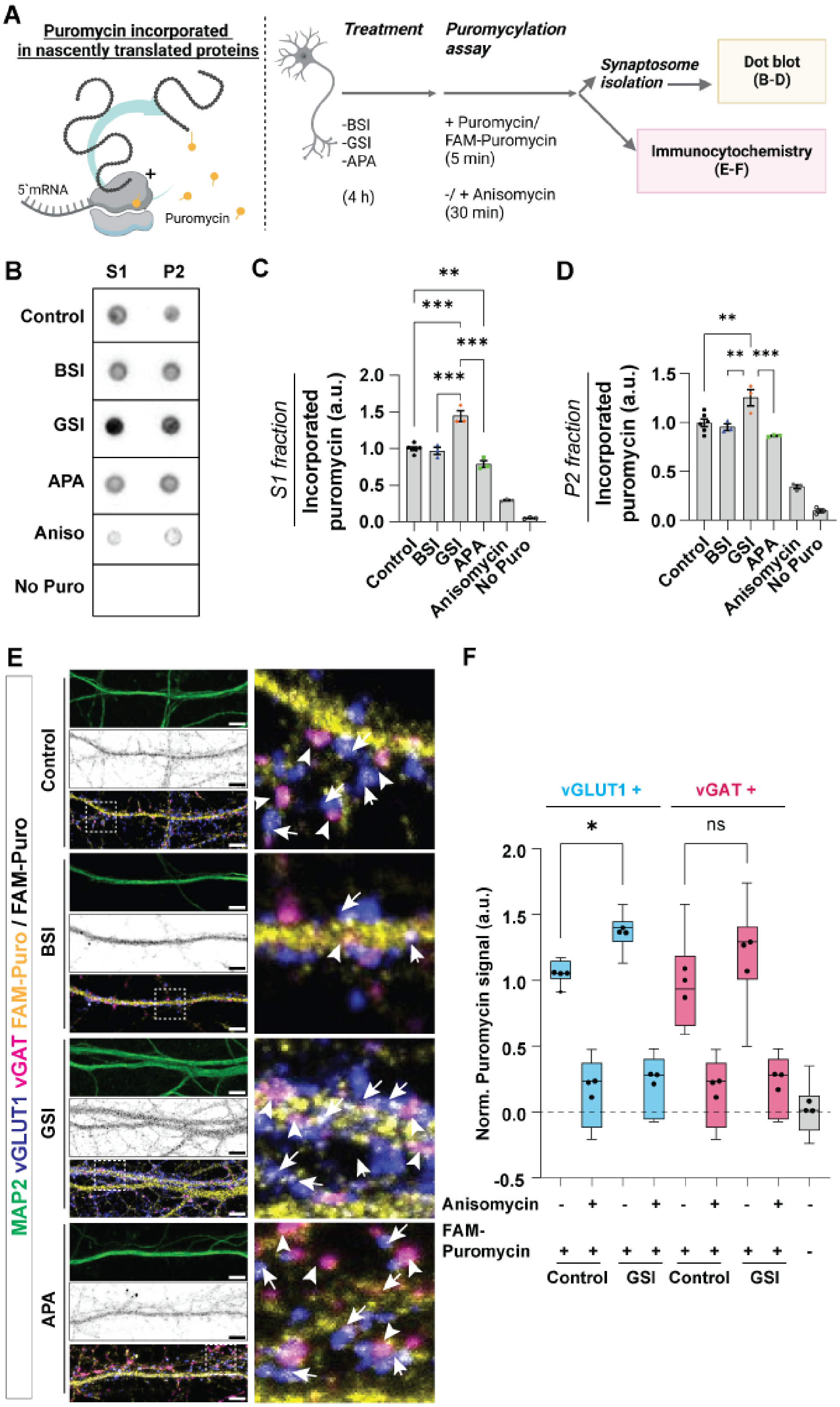

A
Puromycin incorporated in nascently translated proteins
5`mRNA
Puromycin
Treatment
Puromycylation assay
-BSI
-GSI
-APA
(4 h)
+ Puromycin/ FAM-Puromycin (5 min)
-/ + Anisomycin (30 min)
Synaptosome isolation
Dot blot (B-D)
Immunocytochemistry (E-F)
B
S1
P2
Control
BSI
GSI
APA
Aniso
No Puro
C
S1 fraction
Incorporated puromycin (a.u.)
Control
BSI
GSI
APA
Anisomycin
No Puro
D
P2 fraction
Incorporated puromycin (a.u.)
E
MAP2 vGLUT1 vGAT FAM-Puro / FAM-Puro
Control
BSI
GSI
APA
F
vGLUT1 +
vGAT +
ns
Norm. Puromycin signal (a.u.)
Anisomycin
FAM-Puromycin
Control
GSI
Control
GSI

**Figure S5. Puromycylation assay for nascent protein synthesis in neuronal fractions and synapses following APP proteolysis manipulation.** (**A**) Schematic representation depicting the flow for puromycylation assay to examine effects on nascent protein synthesis upon pharmacological manipulation of APP proteolysis. Puromycin (1 µM, 5 min; FAM-Puromycin 500 nM, 5 min) incorporated into the proteome was detected using western immunoblotting or ICC analysis. Anisomycin (40 µM, 30 min) was used to inhibit protein synthesis. (**B-D**) Puromycin incorporated in neuronal S1 (**C**) and synaptoneurosomes enriched P2 (**D**) fractions were detected using dot blot assay using an antibody against puromycin. Samples not treated with puromycin was used to correct the background (*blank values*) and normalized to values of control samples. (**E-F**) Nascently translated proteins were imaged using FAM-tagged puromycin and imaged using ICC, wherein neurons were co-stained with vGLUT1, vGAT and MAP2 to visualize the excitatory, inhibitory synapses and dendrites, respectively. Arrows and arrowheads indicate localization of FAM-Puro within VGLUT1 and vGAT synapses, respectively. Scale bar = 5 µm. ROI selection was created using vGLUT1 and vGAT signals, superimposed on FAM puro images and extracted values were computed to compare changes in local protein synthesis in excitatory (*blue*) and inhibitory synapses (*pink*), respectively (**F**). In each treatment condition, anisomycin (40 µM, 30 min) was used as control. Values are summarized from three independent experiments. n = 6, N = 3. ns ($p > 0.05$), * ($p \leq 0.05$), ** ($p \leq 0.01$) , *** ($p \leq 0.01$); One-way ANOVA.

**Figure S6.**

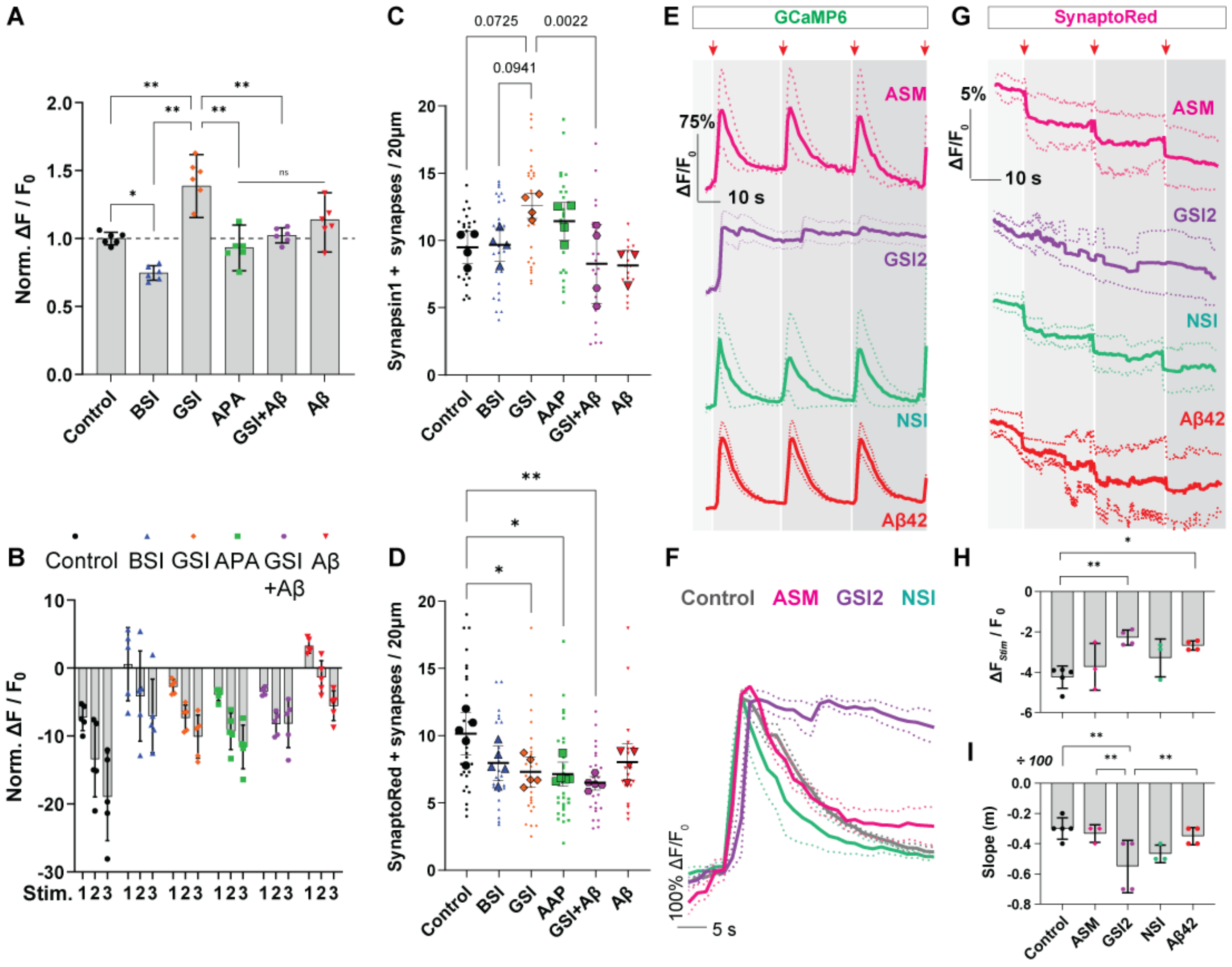


**Figure S6. Effect of pharmacological modulation of APP proteolysis on synaptic vesicle release, synaptic density, and neuronal $Ca^{2+}$ responses.** (**A**) Bar plot depicts the normalized $\Delta F/F_0$ of $Ca^{2+}$ transients, following pharmacological modulation of APP proteolysis. Each data point indicates an experimental replicate. Normalized data from each stimulation. N = 3. (**B**) Bar graph depicts % decrease in fluorescence intensity of SR (ΔF/F) demonstrating synaptic vesicle release (just after stimulation, first-1; second-2, third-3), (**C-D**) Neurons treated with different modulators were either stained with Synapsin1 (**C**) or loaded with SR dye (**D**) and fold change in synaptic density (per um) was quantified and depicted as scatter dot plots. Although GSI-treated conditions show an increase in synaptic density seen in ICC (primarily vGLUT1 positive synapses), these synapses show deficiency in SR dye loading capacity. For control, BSI, GSI, APA, GSI+A$\beta_{42}$, N = 4; A$\beta_{42}$, N = 3. Live labeling of SR dye, control, BSI, GSI, APA, GSI+A$\beta_{42}$, n = 15, N = 5; A$\beta_{42}$, n = 12, N = 4. (**E-I**) Neurons expressing AAV.Camk2.GCaMP6 (DIV3, transduction) were subjected to pharmacological treatment to manipulate APP proteolysis - GI25423X (1 µM; α-secretase modulator, ASM), Compound E (10 nM; γ-secretase inhibition, GSI2) and FLI-06 (10 µM; inhibitor of the notch signaling pathway, NSI) for 4 h. Post-treatment, neurons were loaded with SR dye and subjected to dual imaging of neuronal $Ca^{2+}$ response and synaptic vesicle release events (*scheme 3A*). Overall changes in neuronal $Ca^{2+}$ response (**E, F**) and SR fluorescence intensity (**G-I**, *quantified from multiple selected ROIs*) are depicted as line plots, respectively. Traces (A, C) depict mean (*bold*) and S.E.M (*dotted lines*). Control, n = 15, N = 5; GSI2, $A\beta_{42}$, n = 12, N = 4; ASM, NSI, n = 9, N = 3. Electrical stimulation after GSI2 treatment resulted in stronger effects on the neurons, altering neuronal health. Rescue of network hyperexcitation by $A\beta_{42}$ treatment resulted in neuronal blebbing, thus, further experiments were done only with GSI (DAPT) treatment. In comparison to DAPT, Compound E is an irreversible inhibitor of gamma-secretase, resulting in the observed severe phenotype. (**H**) Bar graph depicts % decrease in fluorescence intensity of SR (ΔF/F) before and just vesicle after stimulation, Control, n = 15, N = 5; GSI2, $A\beta_{42}$, n = 12, N = 4; ASM, NSI, n = 9, N = 3. (**I**) SR fluorescence values recorded during the initial 6 s before stimulation was linear fitted, slopes calculated and plotted as bar graphs. Control, n = 15, N = 5; GSI2, $A\beta_{42}$, n = 12, N = 4; ASM, NSI, n = 9, N = 3. ns ($p > 0.05$), * ($p \leq 0.05$), ** ($p \leq 0.01$) , *** ($p \leq 0.01$); One-way ANOVA.

**Figure S7.**

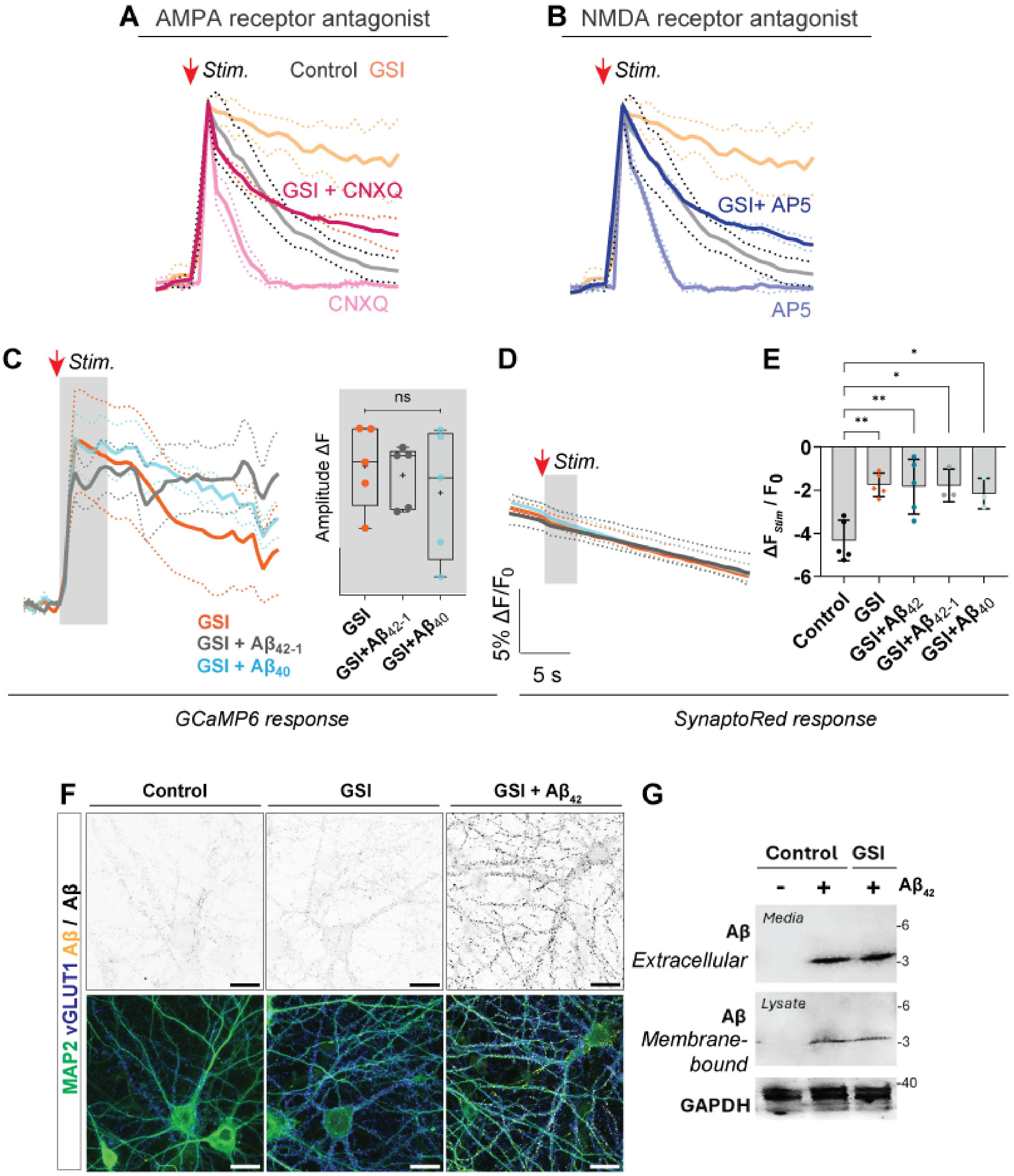


**Figure S7. Effects of post-synaptic receptor antagonism, protein synthesis inhibition, and Aβ42 binding to neuronal membranes.** (**A-B**) Neurons expressing Camk2-GCaMP6s were treated with GSI (4 h) were further treated with CNXQ (10 µM, 30 min; blocking post-synaptic

AMPA receptors), or AP5 (5 μM, 30 min, post synaptic NMDA receptor antagonist) to examine whether changes at postsynaptic signaling influence GSI induced network hyperactivity. APMA and NMDA receptor antagonists were added 30 min before the imaging experiments concluding the GSI treatment time point. Experimental values were compared to neurons treated with the inhibitors alone (without GSI) for the respective durations. Traces (A-B) depict mean (*bold*) and S.E.M (*dotted lines*). n = 6, N = 3. (**C-E**) As observed in Fig. 3, exposure to 100 nM of $A\beta_{42}$ (5 min) results in rescue of GSI-induced network hyperactivity. Additionally, in a similar experimental setting we examined $A\beta_{40}$ as well as $A\beta_{42\text{-}1}$ (*reverse sequence, control peptide*), and recorded changes in calcium (**C**) and SR fluorescence response (**D-E**). wherein no significant change in GCaMP6s response, amplitude of the $Ca^{2+}$ transient, or change in SR fluorescence intensity was detected. (**F-G**) Control or GSI-treated or untreated (control) neurons additionally treated with or without $hA\beta_{42}$ (100 nM, 5 min) were further analyzed *via* immunocytochemistry (F) or western blotting (G) with antibody against Aβ epitope. Neurons were further co-stained with antibodies against vGLUT1 (excitatory synapses, *blue*) and MAP2 (dendrites, *green*). $hA\beta_{42}$ is mainly detected extracellularly bound to the neuronal membranes. Western immunoblot analyses of protein enriched samples show that $A\beta_{42}$ in the treatment media and bound to neuronal membranes is monomeric in nature. Also, GSI-induced APP-CTF accumulation did not impact the binding of $A\beta_{42}$ to neuronal membranes. GAPDH was used as a loading control. N = 2. ns ($p > 0.05$), * ($p \leq 0.05$), ** ($p \leq 0.01$) , *** ($p \leq 0.01$); One-way ANOVA.

**Figure S8.**

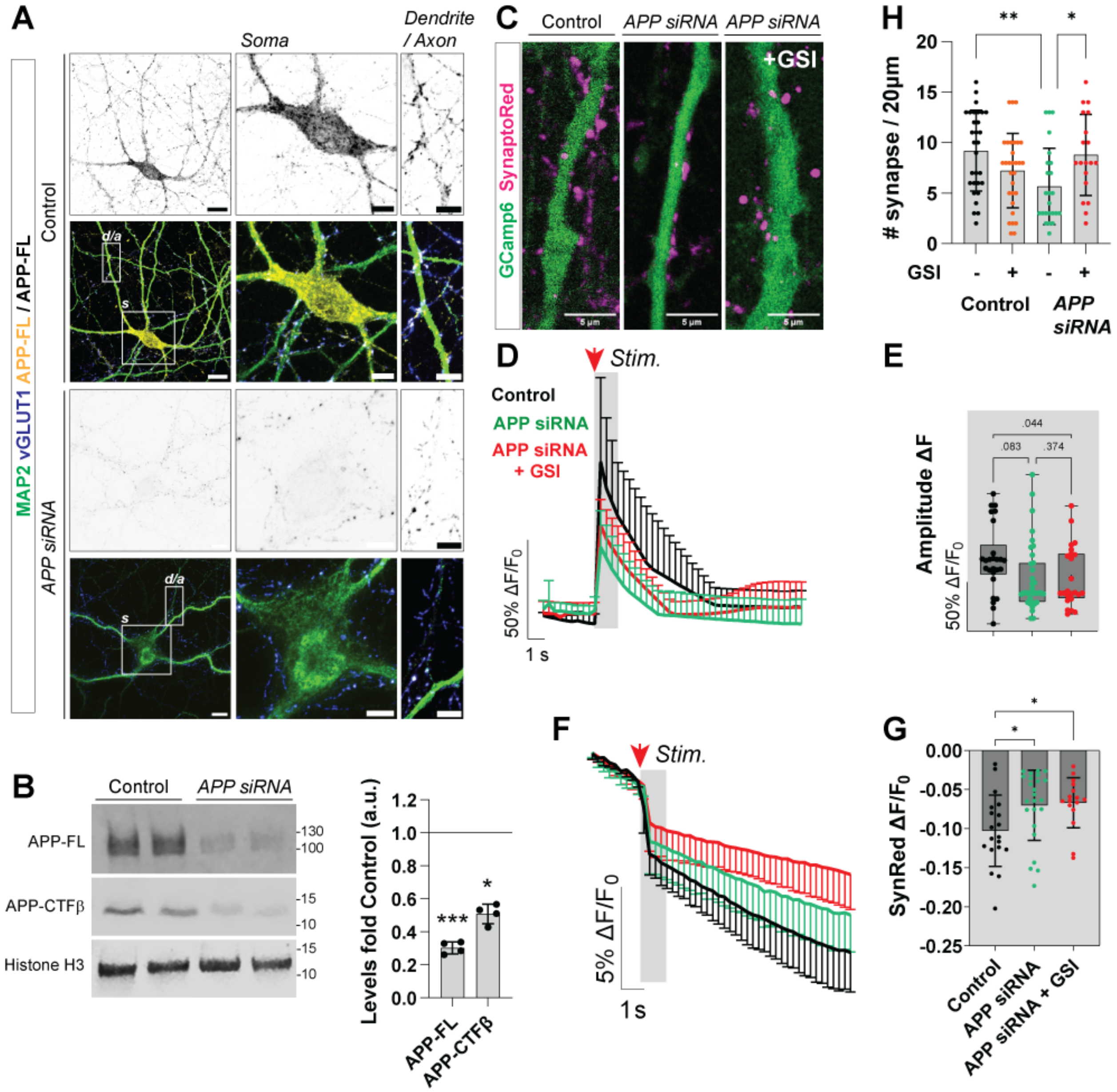


**Figure S8. Effect of loss of APP on synaptic vesicle release and dendritic calcium transients.** (**A-B**) Rat cortical neurons were transfected with an *APP siRNA* (DIV14) and fixed for immunocytochemistry (A) or processed western immunoblot (B) analysis of APP/APP-CTFβ expression in neurons (DIV21). Excitatory synapses were stained with vGLUT1 (*blue*) and dendrites with MAP2 (*green*), respectively. Scale bar = 10 µm. APP-FL levels were compared in soma (s) and dendritic / axons (d/a). Zoomed-in panels are depicted in a white box, scale bar = 5 µm. APP-FL and APP-CTFβ levels were compared in the total neuronal homogenate, wherein histoneH3 was used as loading control. Values were normalized to the control neurons (*line*) and depicted as a bar graph. n = 4, N = 2. (**C-G**) Control and *APP siRNA* treated neurons were further subjected to dual- calcium (GCaMP6s, *green*, **C**) and synaptic vesicle release events (SR measurements, *magenta*, **C**). Neuronal $Ca^{2+}$ traces were quantified and depicted (**D-E**). Loss of APP results in lower amplitude of the $Ca^{2+}$ transient upon neuronal stimulation. GSI treatment does not result in significant differences between neuronal calcium changes. Each data point depicts neuronal soma or dendritic ROI, N = 2. (**F-G**) Changes in levels of SR dye were quantified and depicted as traces (*left,* **F**) and bar blots (*right,* **G**). Loss of APP results in significantly lower decrease in SR fluorescence as compared to the control. GSI treatment does not have any effect on release probability. Each data point depicts a single ROI. N = 2 experimental replicates. (**H**) Number of synapses (per 20 μm) loaded with SR dye were quantified and depicted as a bar plot. Loss of APP results in a significant decrease in the number of synapses but increases upon treatment with GSI. An increase in synapse numbers due to loss of APP and inhibition of γ-secretase indicates that changes in synapse number is independent of APP/APP-CTFβ expression. Of note, there was no significant change in synapse size (APP siRNA, with or without GSI), quantified by the area of SR positive particles. N = 2. ns ($p > 0.05$), * ($p \leq 0.05$), ** ($p \leq 0.01$) , *** ($p \leq 0.01$); One-way ANOVA.

**Figure S9.**

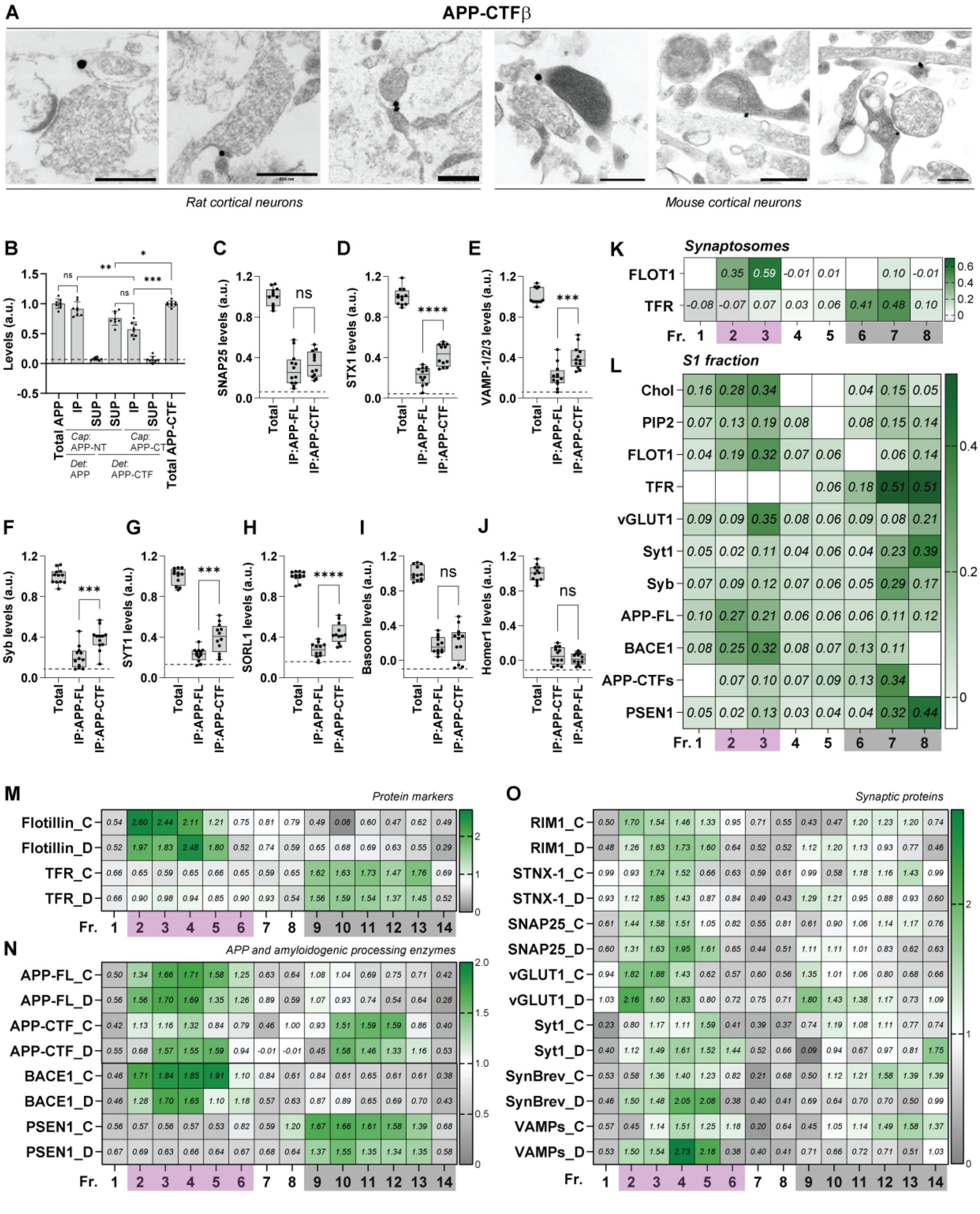


**Figure S9. Subcellular localization and molecular interactions of APP-CTFβ.** (A) Immunogold EM images of APP C-terminal fragment-β (APP-CTFβ) in cultured rat and mouse cortical neuronal synapses. (**B-J**) Dual immunoisolation of presynapse enriched homogenate, first with α-APP-N terminus antibody and later with α-APP-C terminus antibody was done to examine the pool of synaptic proteins interacting with either the full-length APP or APP-CTFβ, respectively. Bar plot (**B**) and box plots (**C-J**) depict absolute values computed from ELISA experiments. Dotted lines represent background values, n = 6, N = 3 independent experiments from different animals. ns ($p > 0.05$), * ($p \leq 0.05$), ** ($p \leq 0.01$) , *** ($p \leq 0.01$); One-way ANOVA.(**K**) Heat map depicts the normalized percentage of flotillin (FLOT1) and transferrin receptor (TFR) within synaptosome material fractionated into PIP2 + cholesterol enriched and perisynaptic fractions. n = 8, N = 4. (**L**) Subsequently, S1 fraction was also subjected to sucrose gradient ultracentrifugation to segregate lipid ordered (cholesterol rich) and disordered fractions (cholesterol depleted). Heat maps depict the normalized percentage of individual lipid material, relative pool of synaptic proteins as well as APP, APP-CTFs and its processing enzymes within the two distinct pools. n = 8, N = 4. **M-O.** Synaptosomes from cultured primary neurons untreated (Ctrl, C) or GSI treated (DAPT, 10 μM, 4 h; D), were subjected to density gradient centrifugation using a similar protocol to fractionate Chol and PIP2 enriched and perisynaptic fractions. Heat map depicts the normalized percentage of flotillin (FLOT1, M) and transferrin receptor (TFR, M), APP, APP-CTFs and processing enzymes (N), along with various synaptic proteins (O) within the different fractions. n = 4, N = 2.

**Figure S10.**

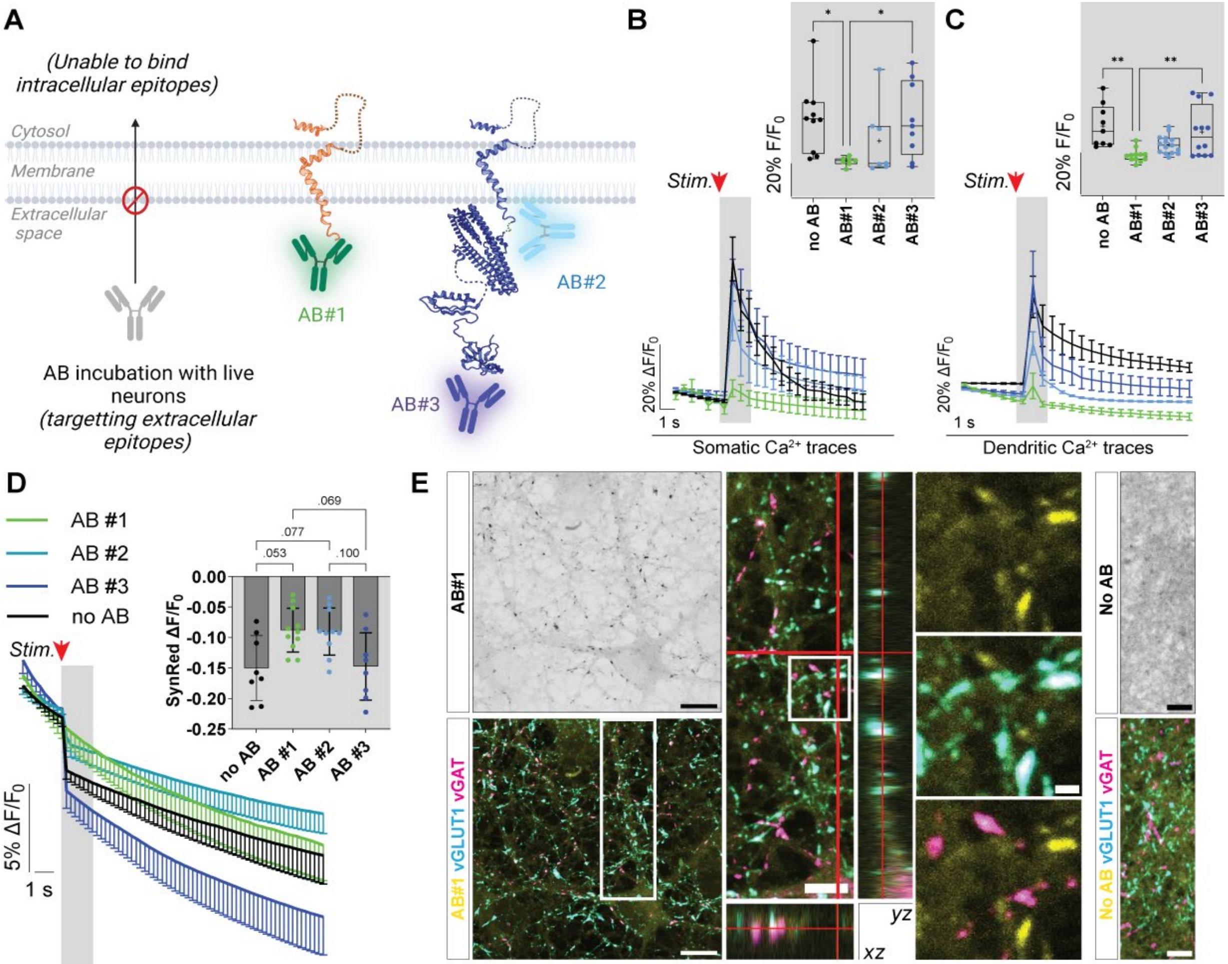


**Figure S10. Live antibody labeling, calcium imaging, synaptic vesicle release, and synaptic localization analyses in cultured neurons.** (**A**) Schematic representation depicting antibodies when incubated with live neurons (1:1000, 10 min) bind to extracellular, but not intracellular epitopes. (**B-D**) Neurons post-treatment was further used for dual-calcium (GCaMP6s **B-C**) and synaptic vesicle release events (SR fluorescence measurements, **D**, inset depicts the quantified F/F0 ratio of SR FL intensity). Somatic (**B**) and dendritic (**C**) $Ca^{2+}$ traces were quantified and depicted. Antibody targeting the N-terminal domain of APP-CTFβ (AB#1) show significant changes in $Ca^{2+}$ transients as compared to AB#3, which targets the extracellular ectodomain of full-length APP. AB#2 which binds within the Aβ sequence shows similar trend when compared to AB#1, but the differences are not significant. (**E**) Neurons treated with AB#1 were fixed and co-stained for excitatory (vGLUT1, *cyan*) and inhibitory (vGAT, *magenta*) synapses, scale bar = 10 µm. Orthogonal images depict the fluorescent signals seen within vGLUT1 positive synapses (rectangular zoom, scale bar = 5 µm). White boxes depict zoomed-in panels, scale bar = 2 µm. ICC of neurons stained only with Streptavidin-Alexa Fluor 488 (primary delete) was used to compare for non-specific background binding (*right panels*). N = 2. ns ($p > 0.05$), * ($p \leq 0.05$), ** ($p \leq 0.01$) , *** ($p \leq 0.01$); One-way ANOVA.

**Figure S11.**

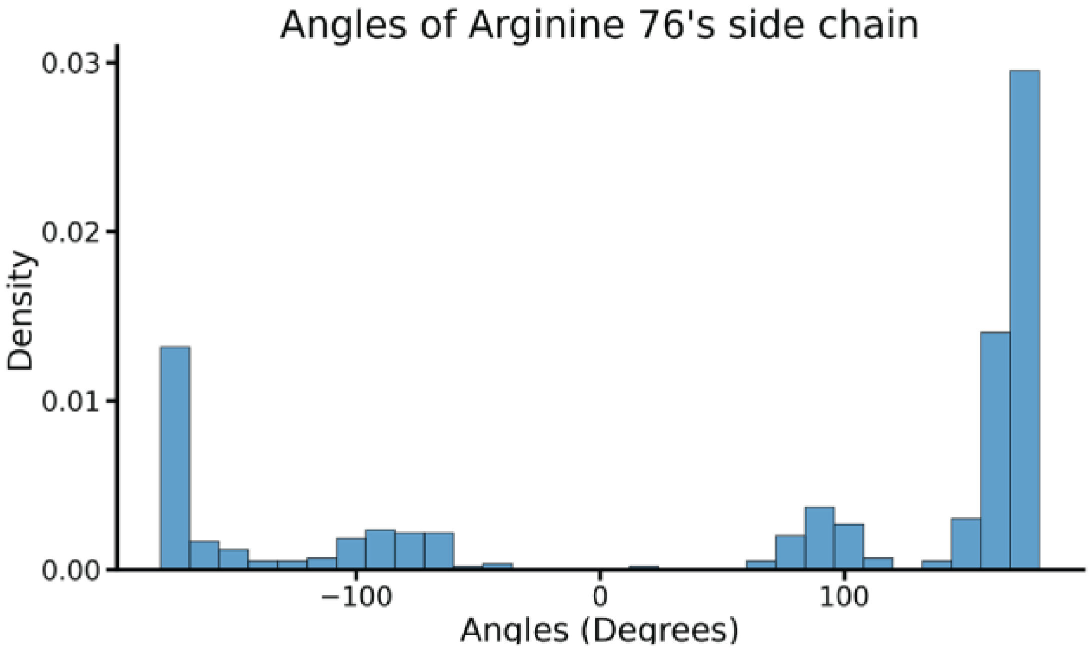


**Figure S11. Dihedral angles of the side chain of APP-CTFβ Arg76.** The dihedral angle spanning its *N* C*A* C*G* C*Z* atom to encapsulate the rotation of the whole side chain spans almost the whole spectrum. Dihedral angles of the side chain of Arg76 of APP-CTFβ in the first simulation to validate the choice of rotation angle. As the arginine's side chain explores the whole space anyway, the initial choice of rotation is meaningless for the simulation.

### Figure S12

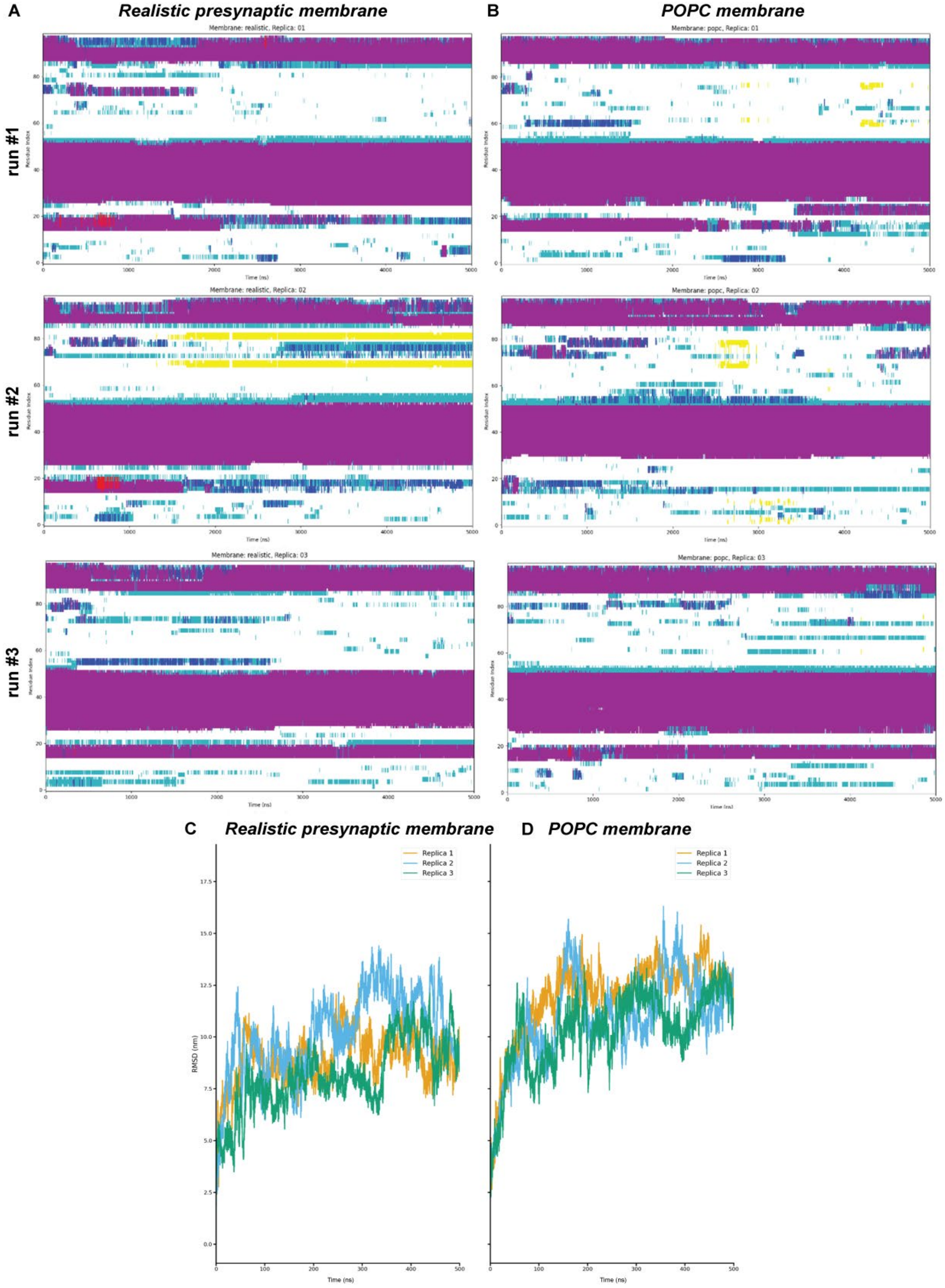


**Figure S12. Secondary structure analysis and APP-CTFβ RSMD plots.** (**A-B**) The secondary structure of the protein is shown over time for either the realistic presynaptic membrane (A) or

POPC only membrane (B), and all replica simulations (run #1-3). Alpha helices, purple; beta sheets, yellow, 3-10 helices, blue; pi helices, red; turns, cyan and random coils, white. (**C, D**) Line plots show the root mean square deviation (RMSD) for all simulations (runs #1-3) for the two different membrane types (C, realistic; D, POPC). It is noticeable that the protein is the most stable, albeit with some fluctuations, in the realistic membrane.

**Figure S13.**

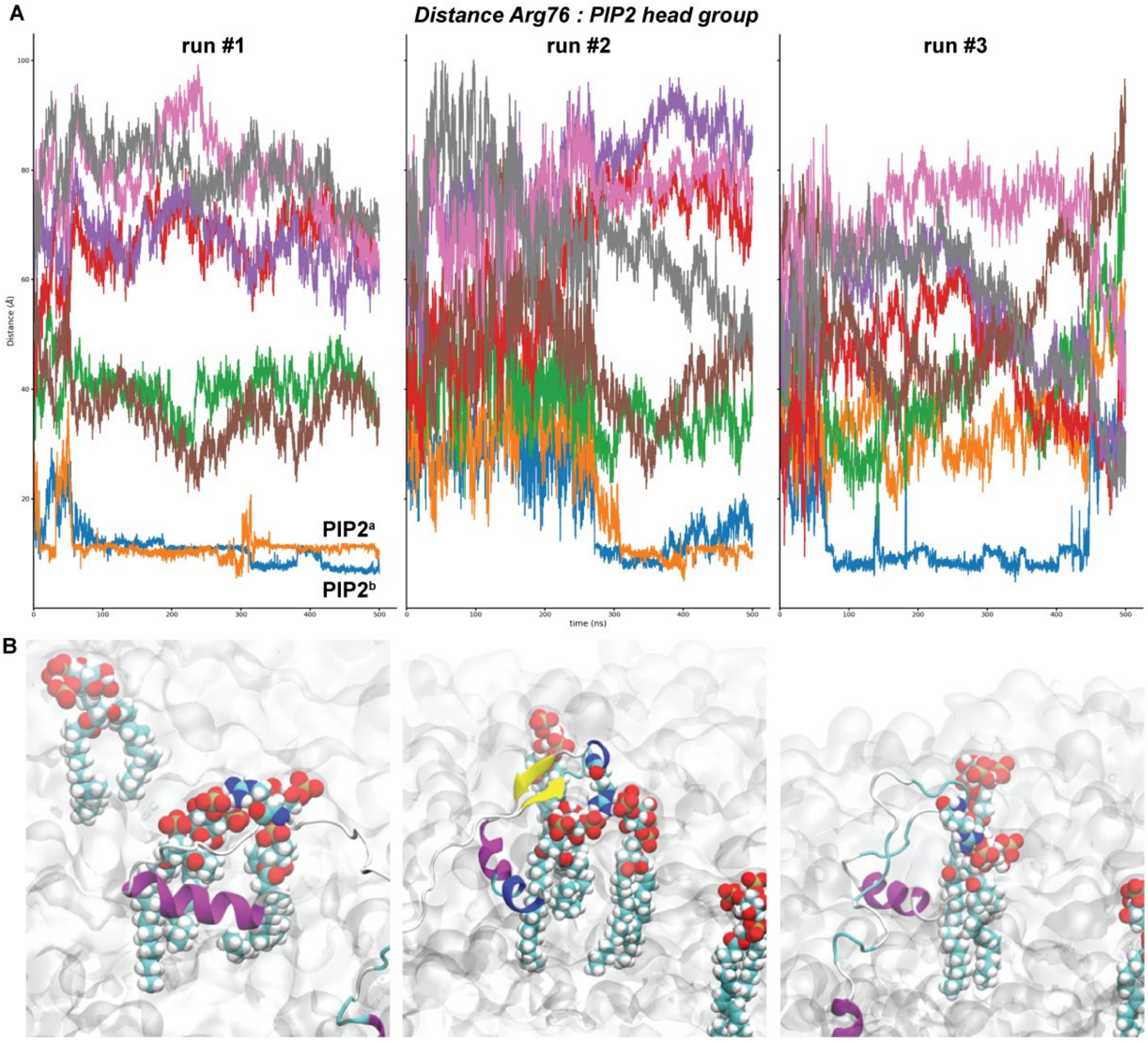


**Figure S13. APP-CTFβ Arg76 interacts with PIP2 head group.** (**A**) For each replica simulation for the realistic membrane, the distance of the final carbon atom in the side chain of arginine 76 is shown to the phosphorus of the PIP2 lipids (8 PIP2 lipids per simulation). Notably, in two replicas, the PIP2 lipids (A, run#1, *a and b*, Fig. 5B) remain stably close to the arginine and are held in place. In the third replica, only one of the PIP2 lipids stays close to the arginine for a considerable time and then departs again. (**B**) The visible parts of the protein are shown in the secondary structure representation with the alpha-helical C-terminal visible in the left image. Arginine 76 is shown in a ball-and-stick representation. The only colored lipids are two PI(4,5)P2 lipids close to the protein. During the simulation, the Arginine not only stabilized the location of the one PIP2 lipid but also facilitated the movement of the second lipid.

**Figure S14**

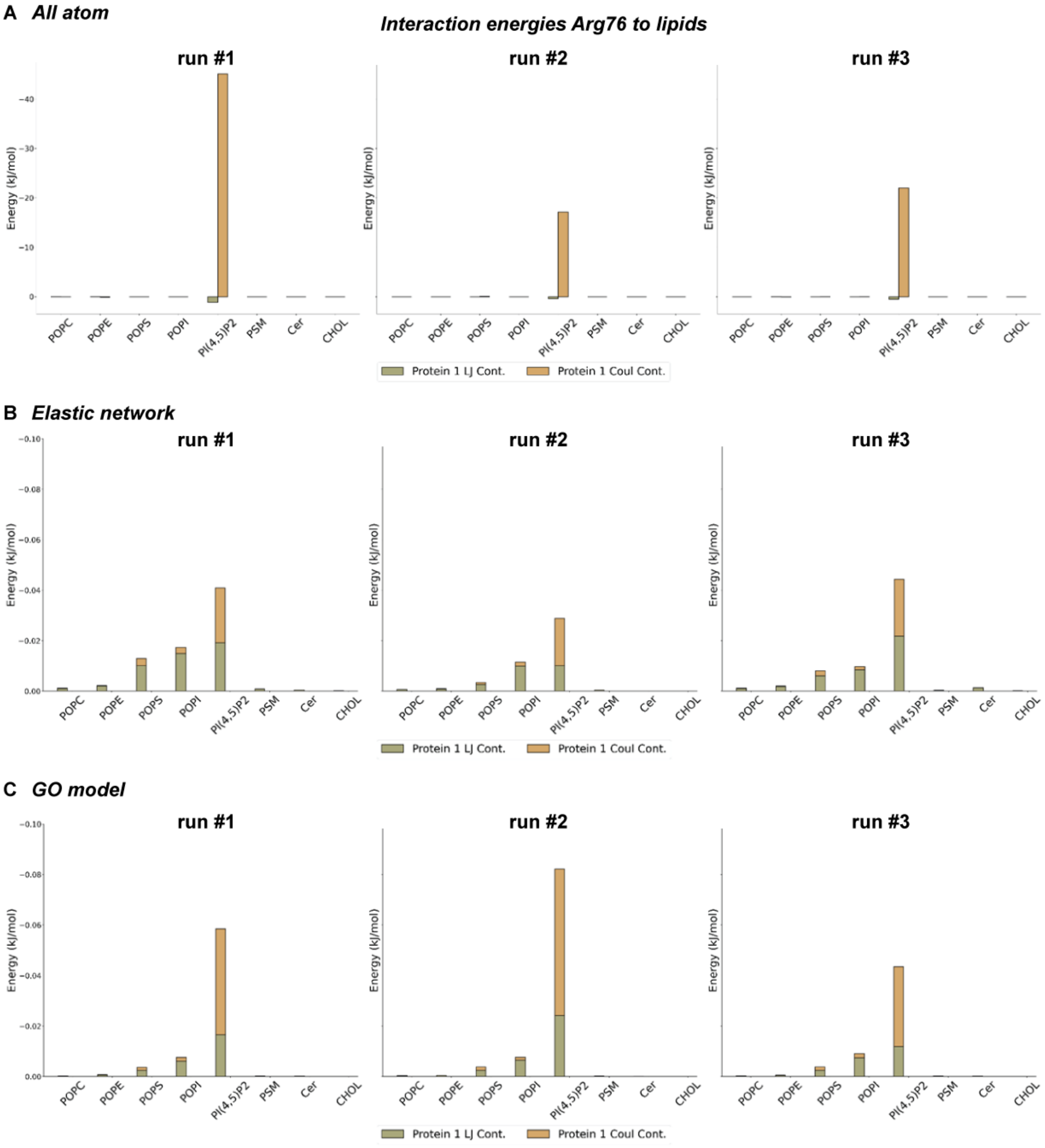


**Figure S14. Computed interaction energy between APP-CTFβ Arg76 and lipid types.** (**A-C**) Total average interaction energy contributions of Arg76 to all lipids of a specific type split into Van der Waals (LJ, Lennard-Jones) and electrostatic (Coulomb) contributions. The contributions have been scaled/averaged by the abundance of the lipids, respectively; for one protein system modelled in the realistic presynaptic membrane for both all-atom (A) and coarse-grained simulations [elastic network (B) and GO model (C)].

**Figure S15.**

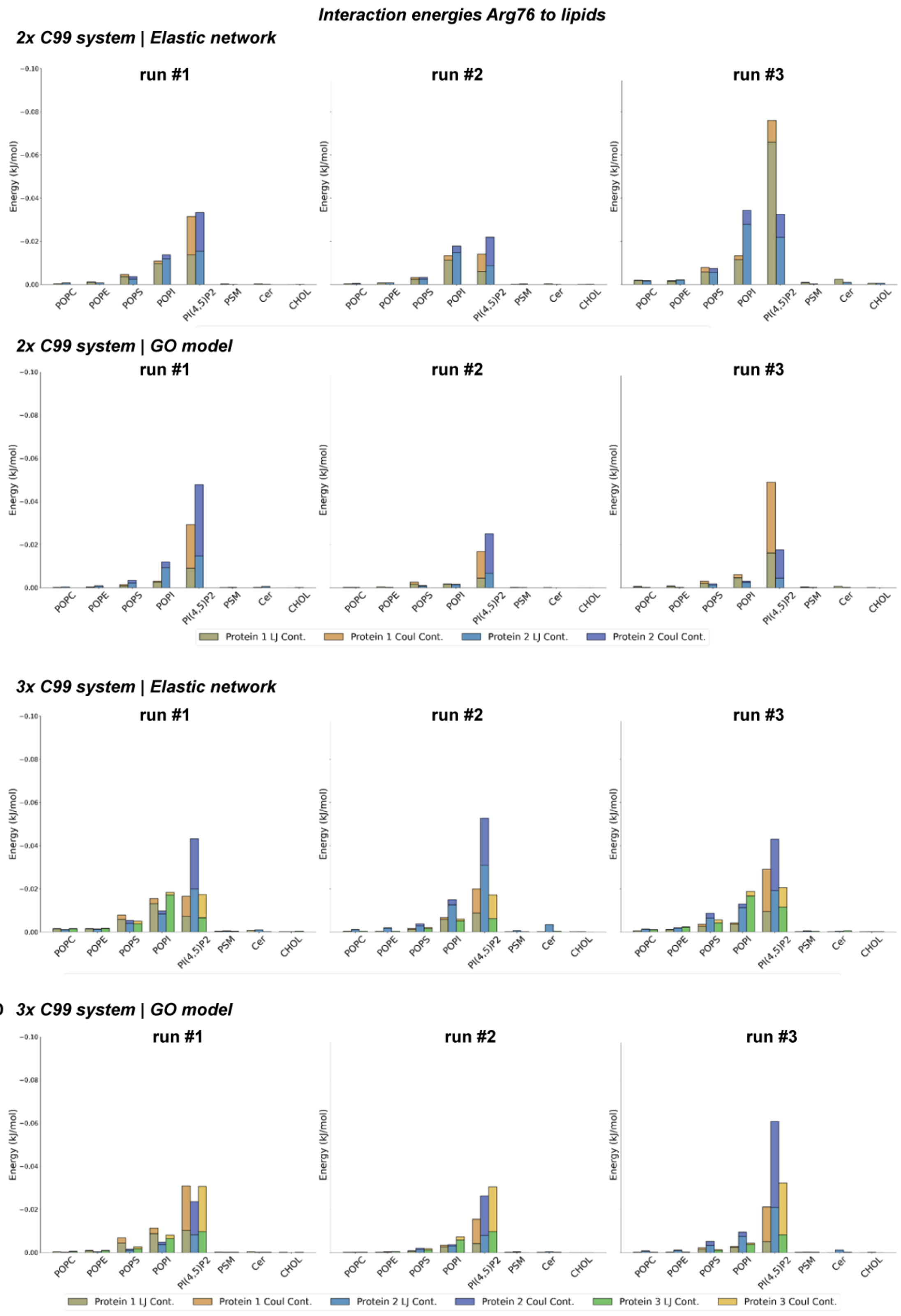

Interaction energies Arg76 to lipids
A 2x C99 system | Elastic network
B 2x C99 system | GO model
C 3x C99 system | Elastic network
D 3x C99 system | GO model
run #1
run #2
run #3
Energy (kJ/mol)
POPC
POPE
POPS
POPI
PI(4,5)P2
PSM
Cer
CHOL
Protein 1 LJ Cont.
Protein 1 Coul Cont.
Protein 2 LJ Cont.
Protein 2 Coul Cont.
Protein 3 LJ Cont.
Protein 3 Coul Cont.

**Figure S15. Effect of molecular crowding dependent on interaction energies between APP-CTFβ Arg76 and lipid types.** (**A-D**) Similar to earlier, total average interaction energy contributions of Arg76 to all lipids of a specific type, split into Van der Waals (LJ, Lennard-Jones) and electrostatic (Coulomb) contributions. The contributions have been scaled/averaged by the abundance of the lipids, respectively; for two (A-B) or three (C-D) protein systems modelled in the realistic presynaptic membrane for coarse-grained simulations [elastic network (A, C) and GO model (B, D)]. Data from C, run#2; depicted in Fig. 5G

**Figure S16.**

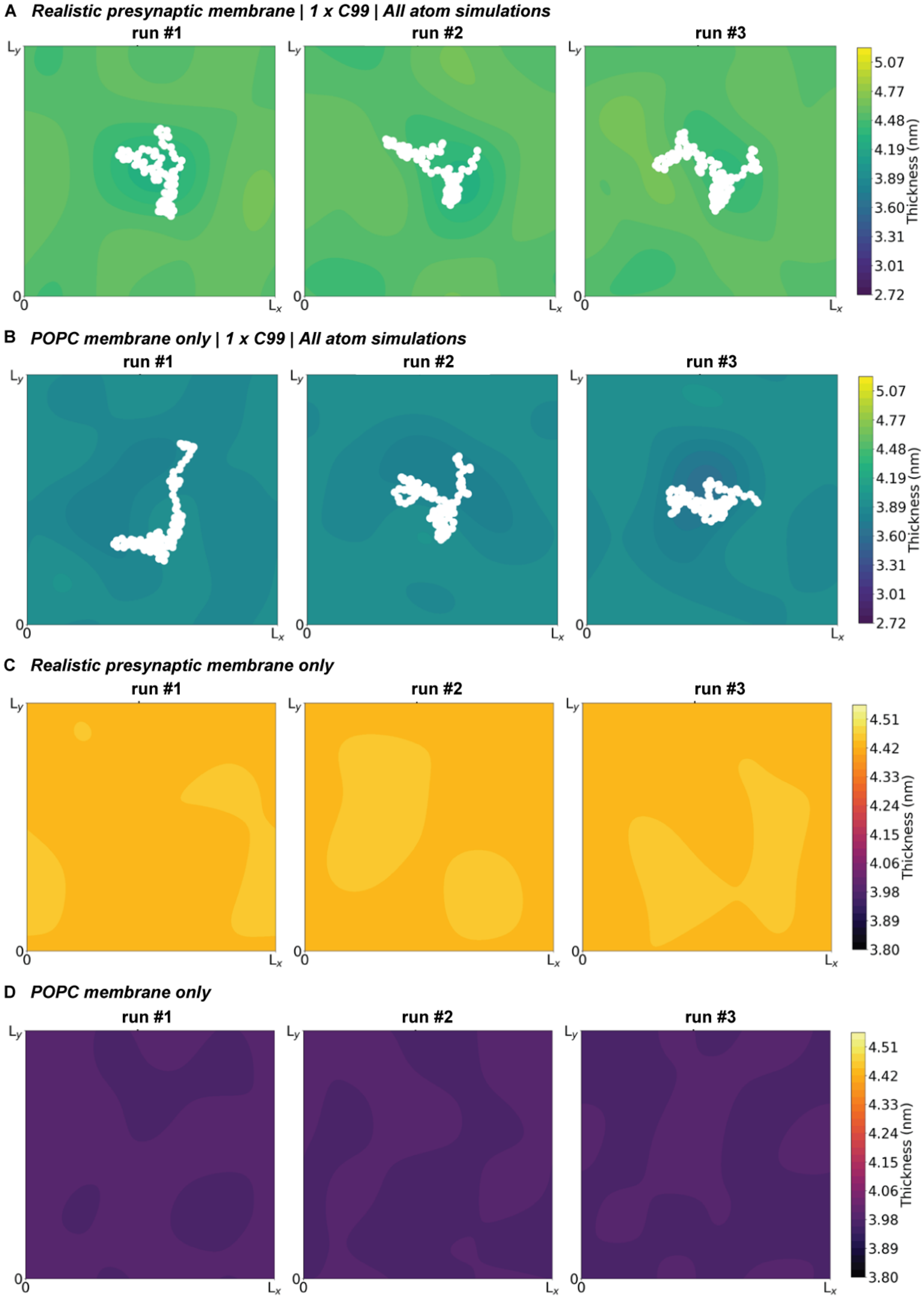


**Figure S16. Effect of APP-CTFβ on membrane thickness.** Heat maps (**A-D**) depict the changes in membrane thickness with (A, B; All atom simulations) or without (C, D, coarse-grained simulations) a single APP-CTFβ in two different membrane systems (A, C, realistic; B, D, POPC membrane). White regions show protein location (CA atoms).

**Figure S17.**

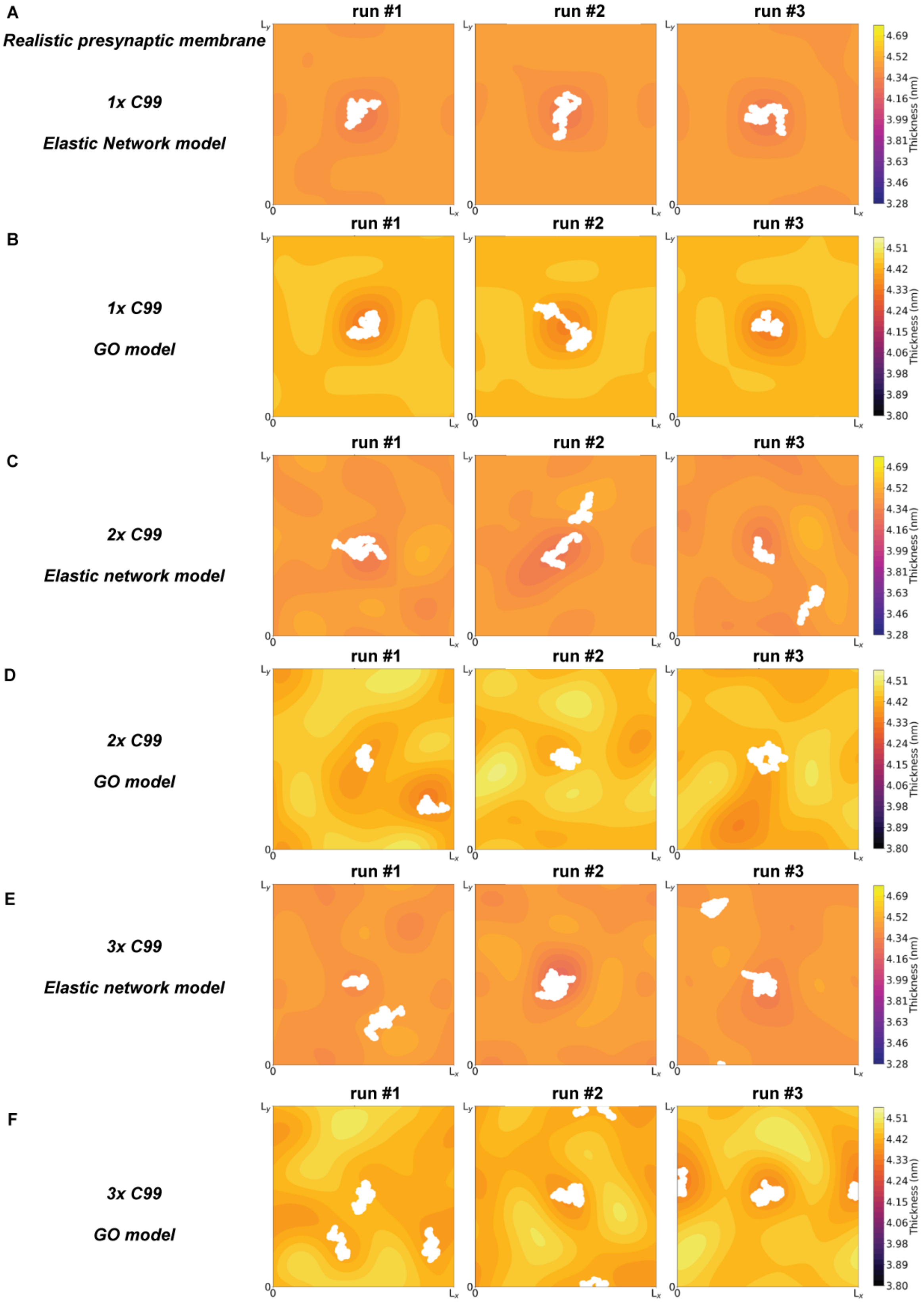
A
Realistic presynaptic membrane
1x C99
Elastic Network model
B
1x C99
GO model
C
2x C99
Elastic network model
D
2x C99
GO model
E
3x C99
Elastic network model
F
3x C99
GO model
run #1
run #2
run #3
Thickness (nm)
4.69
4.52
4.34
4.16
3.99
3.81
3.63
3.46
3.28
4.51
4.42
4.33
4.24
4.15
4.06
3.98
3.89
3.80

**Figure S17. Effect of APP-CTFβ on realistic presynaptic membrane thickness.** Heat maps (**A-F**) depict the changes in membrane thickness in one (A, B), two (C, D) or three (E, F) APP-CTFβ in the realistic presynaptic membranes (A, C, E, elastic network; B, D, F, GO model). White regions show the protein(s) location (backbone beads). Data from A, run#1, C, run#1, and E, run#2; depicted in Fig. 5H panels, respectively.

**Figure S18.**

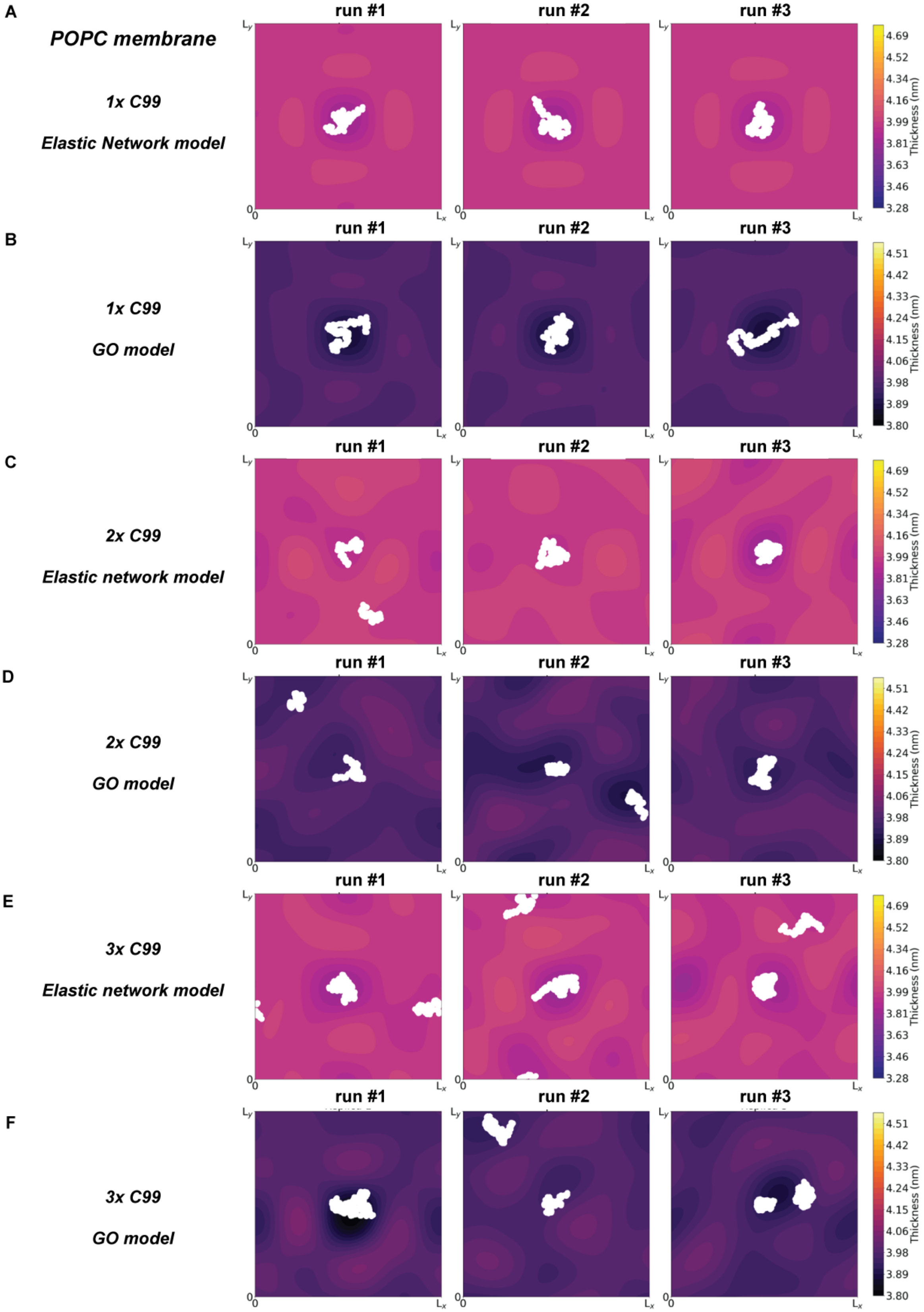
A
POPC membrane
1x C99
Elastic Network model
B
1x C99
GO model
C
2x C99
Elastic network model
D
2x C99
GO model
E
3x C99
Elastic network model
F
3x C99
GO model
run #1
run #2
run #3
Thickness (nm)
4.69
4.52
4.34
4.16
3.99
3.81
3.63
3.46
3.28
4.51
4.42
4.33
4.24
4.15
4.06
3.98
3.89
3.80
$L_y$
$L_x$
0

**Figure S18. Effect of APP-CTFβ on POPC membrane thickness.** Heat maps (**A-F**) depict the changes in membrane thickness in one (A, B), two (C, D) or three (E, F) APP-CTFβ in the realistic presynaptic membranes (A, C, E, elastic network; B, D, F, GO model). White regions show the protein(s) location (backbone beads).

**Figure S19.**

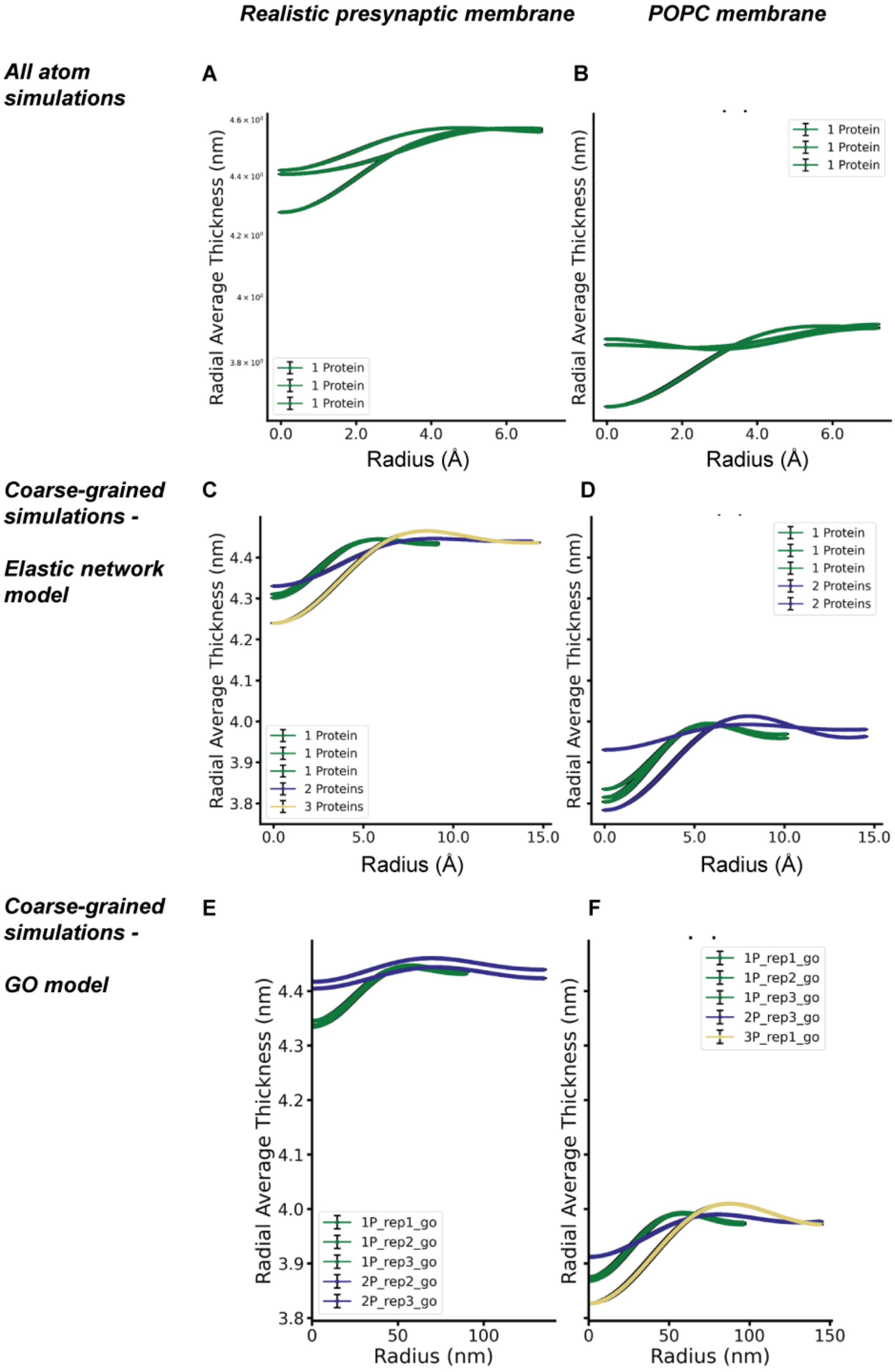

Realistic presynaptic membrane
POPC membrane
All atom simulations
Coarse-grained simulations -
Elastic network model
Coarse-grained simulations -
GO model
A
B
C
D
E
F
Radial Average Thickness (nm)
Radius (Å)
Radius (nm)
1 Protein
2 Proteins
3 Proteins
1P_rep1_go
1P_rep2_go
1P_rep3_go
2P_rep2_go
2P_rep3_go
3P_rep1_go

**Figure S19. Effect of APP-CTFβ transmembrane domain on membrane thinning. A-F.** Thickness profile of the lipid membrane quantified radially (see methods section) from the APP-CTFβ transmembrane domain (1, 2 or 3 protein systems) computed from all-atom simulations (A, B); coarse-grained simulations (C-F) *via* elastic network (C, D) or GO model (E, F). Part of graph C is depicted in Fig. 5K.

**Figure S20.**

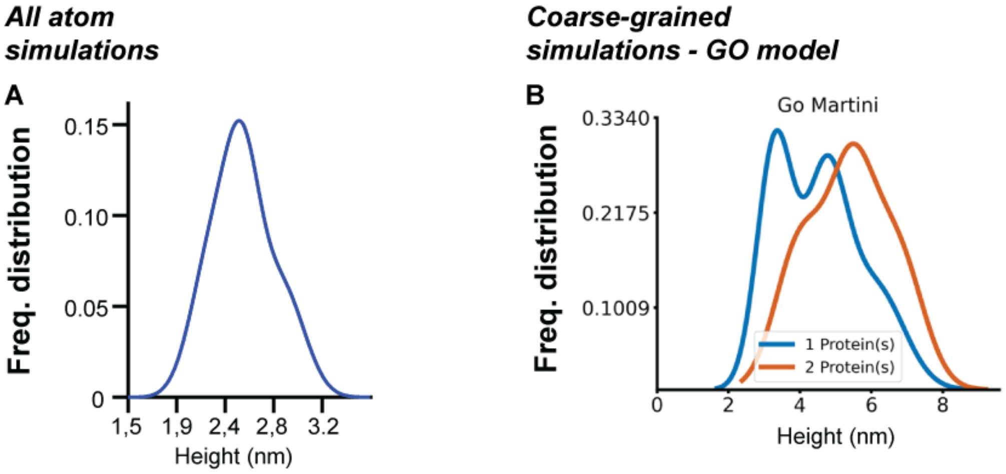


**Figure S20. APP-CTFβ C-terminal Asn99 explores cytosolic lumen.** Line plots (**A, B**) show the height of the last backbone bead (Asn99) of the C-terminal to/from the center of the membrane leaflet (obtained from fitting scheme described above) for all atom (A) and coarse-grained GO model (B) simulations. Data from coarse-grained elastic model simulations are depicted in Fig. 5I.

**Video S1. Spontaneous calcium activity in Camk2-GCaMP6-expressing neurons following pharmacological modulation of APP processing**. Primary neurons (DIV21) transduced with AAV-Camk2-GCaMP6 were treated for 4 h with LY2886721 (1 µM; BACE1 inhibitor, BSI), DAPT (10 µM; γ-secretase inhibitor, GSI), or Aftin-4 (5 µM; APP processing activator, APA) to target distinct stages of APP proteolysis. Spontaneous calcium transients were recorded under $Ca^{2+}$-containing Tyrode buffer.

**Video S2. Experimental workflow for dual imaging of neuronal $Ca^{2+}$ dynamics and synaptic vesicle release.** Neurons expressing AAV.Camk2.GCaMP6 (transduced at DIV3) were treated with pharmacological modulators at DIV21 for 4 h. Synaptic vesicles were then labeled with SynaptoRed (SR), followed by washout and a 5-min rest period. Neurons were stimulated (30 Hz, 4-5 trains starting at 7 s, 30 s recovery), and dual imaging of GCaMP6 (*green*) and SR (*magenta*) signals was performed for ~2 min at 125 ms intervals. Fluorescence changes in individual ROIs (synapses-*magenta* and dendrites-*green*) were quantified.

**Video S3. APP-CTF–induced neuronal hyperexcitation *via* presynaptic dysfunction.** Neurons expressing AAV.Camk2.GCaMP6 were treated with pharmacological modulators to alter APP processing (as above). Dual imaging of neuronal $Ca^{2+}$ dynamics (GCaMP6, *green*) and synaptic vesicle release (SR, *magenta*) was performed as indicated above. GSI treatment resulted in enhanced $Ca^{2+}$ responses and impaired vesicle release consistent with presynaptic dysfunction.

**Video S4. Aβ rescues γ-secretase inhibitor (GSI)-induced hyperactivity via postsynaptic mechanisms.** Neurons were treated with γ-secretase inhibitor (GSI) to increase APP-CTF accumulation. GSI pretreated or untreated neurons were subjected to acute treatment with Aβ for 5 min, before imaging. Dual imaging of neuronal $Ca^{2+}$ dynamics (GCaMP6, *green*) and synaptic vesicle release (SR, *magenta*) was performed as indicated above. Data indicate that Aβ treatment normalizes GSI-induced hyperactivity, implicating a postsynaptic mechanism of rescue.

**Video S5. Molecular dynamics simulation of APP-CTFβ (C99) in a realistic presynaptic membrane.** The video depicts APP-CTFβ (C99) embedded in a lipid bilayer that approximates the composition of a presynaptic membrane. This simulation models C99 in a physiologically relevant environment, providing insight into its potential role in membrane lipid signaling.

### Appendix: vGLUT1$^{GFP}$ transgenic mice

Generation and characterization of the knock in mouse model by CRISPR/Cas-mediated genome editing technique was outsourced to Cyagen (https://www.cyagen.com/)

1. Schematic depicting the targeting strategy

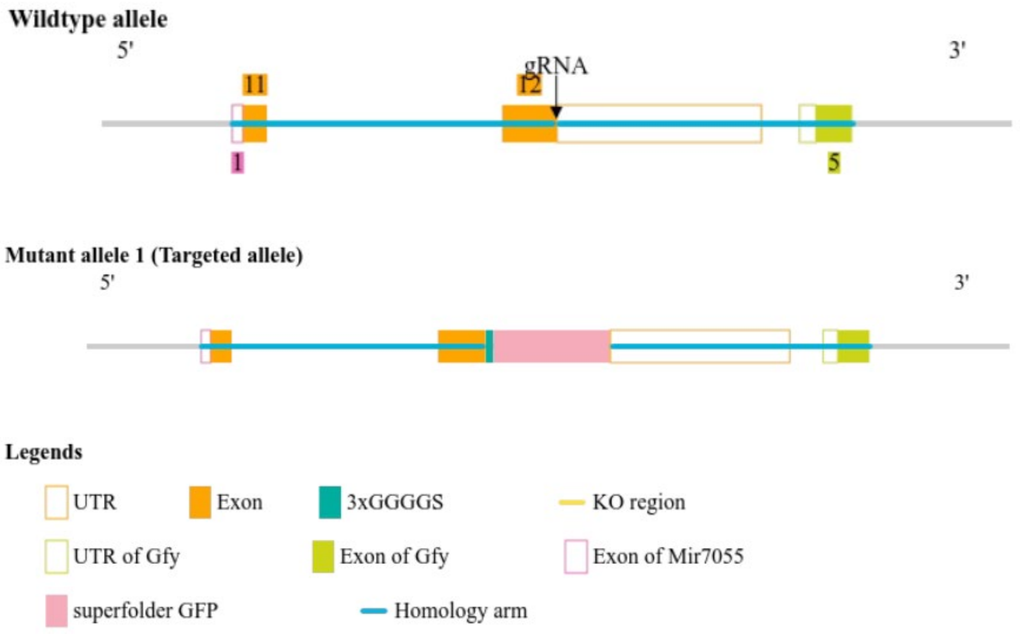

### 2. Gene and protein information

**Slc17a7 solute carrier family 17 (sodium-dependent inorganic phosphate cotransporter), member 7 [ *Mus musculus* (house mouse) ]**

Gene ID: 72961, updated on 16-Mar-2021

**Gene summary**

| | |
|---|---|
| **Official Symbol** | Slc17a7provided by MGI |
| **Official Full Name** | solute carrier family 17 (sodium-dependent inorganic phosphate cotransporter), member 7provided by MGI |
| **Primary source** | MGI:MGI:1920211 |
| **See related** | Ensembl:ENSMUSG00000070570 |
| **Gene type** | protein coding |
| **RefSeq status** | VALIDATED |
| **Organism** | Mus musculus |
| **Lineage** | Eukaryota; Metazoa; Chordata; Craniata; Vertebrata; Euteleostomi; Mammalia; Eutheria; Euarchontoglires; Glires; Rodentia; Myomorpha; Muroidea; Muridae; Murinae; Mus; Mus |
| **Also known as** | Vglu; Vglut1; AI851913; 2900052E22Rik |
| **Expression** | Biased expression in cortex adult (RPKM 249.8), frontal lobe adult (RPKM 216.5) and 3 other tissues See more |
| **Orthologs** | human all |
| **NEW** | **Try the new Gene table** |
| | **Try the new Transcript table** |

**Genomic context**

**Location:** 7; 7 B3 See Slc17a7 in Genome Data Viewer

**Exon count:** 12

| Annotation release | Status | Assembly | Chr | Location |
|---|---|---|---|---|
| 109 | current | GRCm39 (GCF_000001635.27) | 7 | NC_000073.7 (44813345..44825563) |
| 108.20200622 | previous assembly | GRCm38.p6 (GCF_000001635.26) | 7 | NC_000073.6 (45163921..45176139) |
| Build 37.2 | previous assembly | MGSCv37 (GCF_000001635.18) | 7 | NC_000073.5 (52419291..52431509) |

**Chromosome 7 - NC_000073.7**

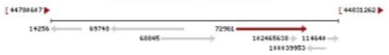

### 3. Transcripts information

Gene: Slc17a7 ENSMUSG00000070570

| | |
|---|---|
| **Description** | solute carrier family 17 (sodium-dependent inorganic phosphate cotransporter), member 7 [Source:MGI Symbol;Acc:MGI:1920211] |
| **Gene Synonyms** | 2900052E22Rik, Vglut1 |
| **Location** | Chromosome 7: 44,813,373-44,825,566 forward strand.<br>GRCm39:CM001000.3 |
| **About this gene** | This gene has 5 transcripts (splice variants), 322 orthologues, 12 paralogues and is associated with 28 phenotypes. |
| **Transcripts** | |

| Name | Transcript ID | bp | Protein | Translation ID | Biotype | CCDS | UniProt Match | Flags |
|---|---|---|---|---|---|---|---|---|
| Slc17a7-201 | ENSMUST00000085374.7 | 2915 | 560aa | ENSMUSP00000082489.6 | Protein coding | CCDS52244 | Q3TXX4 | TSL:1GENCODE basicAPPRIS P1 |
| Slc17a7-202 | ENSMUST00000209634.2 | 1860 | 585aa | ENSMUSP00000147661.2 | Protein coding | - | A0A1B0GRU0 | TSL:5GENCODE basic |
| Slc17a7-205 | ENSMUST00000211652.2 | 3080 | No protein | - | Retained intron | - | - | TSL:2 |
| Slc17a7-204 | ENSMUST00000210540.2 | 2722 | No protein | - | Retained intron | - | - | TSL:1 |
| Slc17a7-203 | ENSMUST00000210498.2 | 2346 | No protein | - | Retained intron | - | - | TSL:NA |

Ensembl release 103 - February 2021 ©

4. Gene expression/activity chart

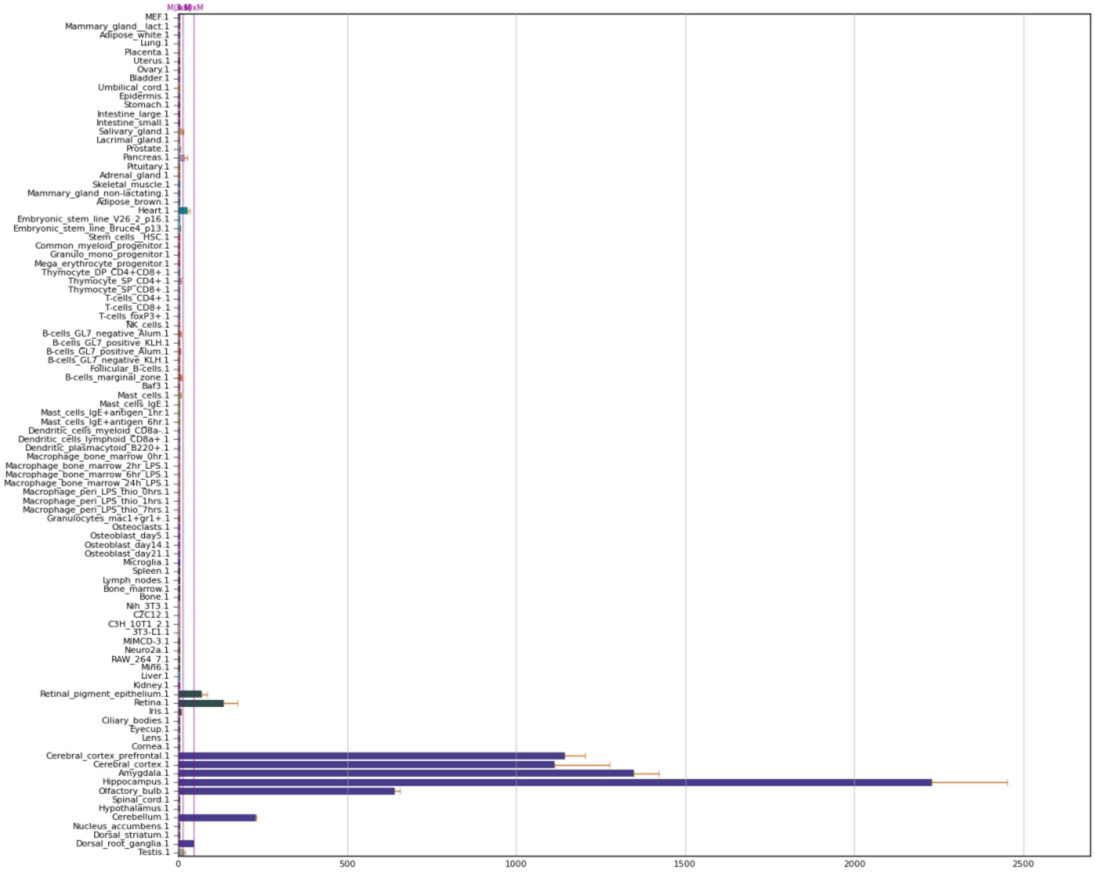

**Data S1. (separate file)**

Raw numerical values used for preparing graphical representations in the main manuscript Fig. 1-5 have been provided in a separate excel file.
